# Precision calcium imaging of dense neural populations via a cell body-targeted calcium indicator

**DOI:** 10.1101/773069

**Authors:** Or A. Shemesh, Changyang Linghu, Kiryl D. Piatkevich, Daniel Goodwin, Howard J. Gritton, Michael F. Romano, Cody Siciliano, Ruixuan Gao, Chi-Chieh (Jay) Yu, Hua-An Tseng, Seth Bensussen, Sujatha Narayan, Chao-Tsung Yang, Limor Freifeld, Ishan Gupta, Habiba Noamany, Nikita Pak, Young-Gyu Yoon, Jeremy F.P. Ullmann, Burcu Guner-Ataman, Zoe R. Sheinkopf, Won Min Park, Shoh Asano, Amy E. Keating, James S. Trimmer, Jacob Reimer, Andreas Tolias, Kay M. Tye, Xue Han, Misha B. Ahrens, Edward S. Boyden

## Abstract

Methods for one-photon fluorescent imaging of calcium dynamics in vivo are popular due to their ability to simultaneously capture the dynamics of hundreds of neurons across large fields of view, at a low equipment complexity and cost. In contrast to two-photon methods, however, one-photon methods suffer from higher levels of crosstalk between cell bodies and the surrounding neuropil, resulting in decreased signal-to-noise and artifactual correlations of neural activity. Here, we address this problem by engineering cell body-targeted variants of the fluorescent calcium indicator GCaMP6f. We screened fusions of GCaMP6f to both natural as well as engineered peptides, and identified fusions that localized GCaMP6f to within approximately 50 microns of the cell body of neurons in live mice and larval zebrafish. One-photon imaging of soma-targeted GCaMP6f in dense neural circuits reported fewer artifactual spikes from neuropil, increased signal-to-noise ratio, and decreased artifactual correlation across neurons. Thus, soma-targeting of fluorescent calcium indicators increases neuronal signal fidelity and may facilitate even greater usage of simple, powerful, one-photon methods of population imaging of neural calcium dynamics.

## Introduction

In recent years, methods for one-photon fluorescent imaging of calcium dynamics in vivo, including epifluorescent, endoscopic, and light-sheet methods, have become popular techniques for neural activity mapping in living larval zebrafish, mice, and other species. In conjunction with fluorescent calcium indicators, these techniques capture, at high speeds (e.g., 20 Hz or more), the dynamics of hundreds of neurons across large fields of view, at a low equipment complexity and cost (Alivisatos et al., 2013; Grienberger and Konnerth, 2012; Keller et al., 2015). For the purposes of neural spike extraction, neuroscientists typically focus on analyzing the data from cell bodies of neurons being imaged. Both two-photon (Chen et al., 2013; Helmchen and Denk, 2005; Raichle et al., 1998) and one-photon imaging modalities have resolution limits that will typically mix signals from cell bodies with those from closely passing axons and dendrites, resulting in artifactual correlations of measured neural activity (**Fig. 1A-1B**). This crosstalk between neuropil signals and cell body signals can be somewhat mitigated in two-photon microscopy by restricting measurements to the interiors of cell bodies; the crosstalk problem is worse for one-photon epifluorescent methods, where contamination of somatic signals by neuropil signals may be impossible to overcome (Chen et al., 2013; Harris et al., 2016; Peron et al., 2015).

**Figure 1.**
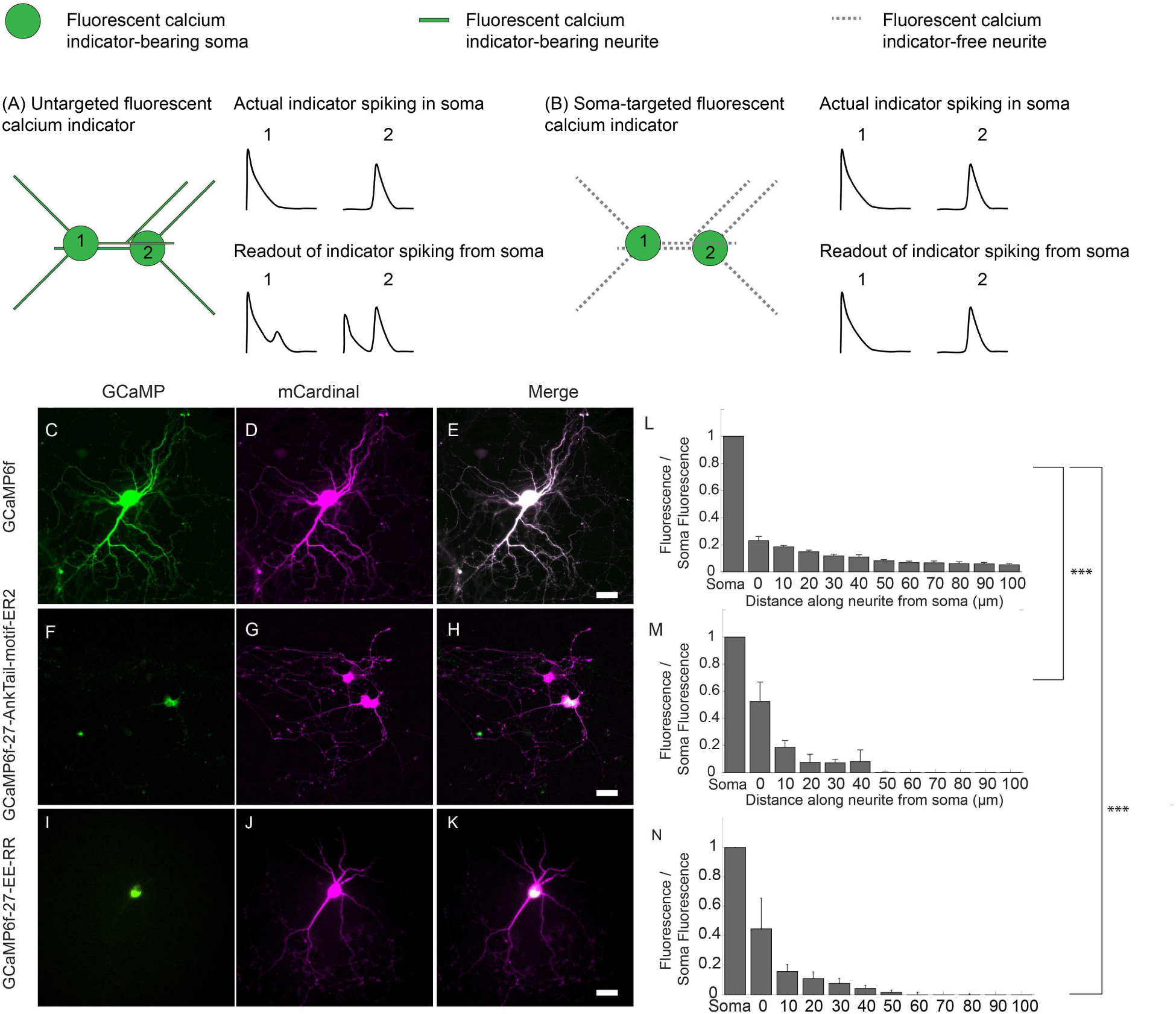
Somatic GCaMP6f variants. Untargeted GCaMP expresses throughout the neural cytosol. One can image several cells, but each cell body is surrounded by GCaMP-bearing neurites from other cells (**A**), which can bleed into the signals attributed to a given cell body (compare “actual” to “readout”). Restricting GCaMP expression to the cell body would enable imaging at single cell resolution (**B**), because neurites cannot contribute bleedthrough signal to a cell body of interest. (**C-K**) Representative images are presented for cultured hippocampal neurons expressing wild-type vs. selectively soma-targeted GCaMP6f variants, as well as the countermarker mCardinal. (**C**) A hippocampal neuron in culture expressing GCaMP6f and mCardinal, seen in the GFP channel. (**D**) The neuron of C, seen in the mCardinal channel (magenta). (**E**) Merge of **C** and **D**. (**F-H**) As in **C-E**, for a neuron expressing GCaMP6f-27-AnkTail-motif-ER2 (termed SomaGCaMP6f1). (**I-K**) As in **C-E**, for a neuron expressing GCaMP6f-27-EE-RR (termed SomaGCaMP6f2). Scale bars for **E, H, K**: 20 µm. (**L**) Bar plot of GCaMP6f brightness versus position along a neurite, normalized to GCaMP6f brightness at the soma, extracted from neurites of cultured hippocampal neurons expressing GCaMP6f (n = 8 neurites from 8 cells from 3 cultures). (**M**) As in **L**, for neurons expressing GCaMP6f-27-AnkTail-motif-ER2 (SomaGCaMP6f1; n = 5 neurites from 5 cells from 2 cultures). ***P < 0.001, Kruskal-Wallis analysis of variance of neurite brightness followed by post-hoc test via Steel’s test with GCaMP6f as a control group; see **Supplemental Table 3** for full statistics for **Figure 1**). (**N**) As in **L**, for neurons expressing GCaMP6f-27-EE-RR (SomaGCaMP6f2; n = 5 neurites from 5 cells from 3 cultures). ***P < 0.001, Kruskal-Wallis analysis of variance of neurite brightness followed by post-hoc test via Steel’s test with GCaMP6f as a control group; see **Supplemental Table 3** for full statistics for **Figure 1**).

As a result, many studies use computational methods to attempt to clean up the in vivo calcium signals, algorithmically correcting somatic signals for the neuropil contribution (Andilla and Hamprecht, 2014; Mukamel et al., 2009; Pinto and Dan, 2015; Pnevmatikakis et al., 2014, 2016). Although such algorithms are widely used in two-photon calcium imaging, one-photon calcium imaging is subject to higher neuropil contamination levels, which remains an open problem for ongoing computational research (Zhou et al., 2016). Furthermore, the contribution of neuropil to observations of a given cell body of interest is only estimated, not exactly measured, through such computational strategies. Accordingly, a second strategy has emerged, namely localizing genetically encoded calcium indicators to the nucleus by fusing them to well-known nuclear localization sequences (NLSs) or histones (H2B), which effectively eliminates the neuropil signal (Bengtson et al., 2010; Kim et al., 2014; Nguyen et al., 2016; Schrödel et al., 2013; Vladimirov et al., 2014). While such nuclear localized calcium indicators do indeed enable low crosstalk imaging of neural populations, even in one-photon microscopy settings, there is a concern that the requirement for calcium to enter the nucleus greatly slows the temporal precision of such imaging, compared to classical cytosolic calcium imaging.

We here confirm that nuclear localized versions of the popular genetically encoded fluorescent calcium indicator GCaMP6f exhibit, in cultured mouse neurons, on and off time constants 3-5x slower than those of cytoplasmic GCaMP6f. We hypothesized that if we could localize a genetically encoded calcium indicator such as GCaMP6f to the cytosol near the cell body, we could greatly reduce neuropil fluorescence, similar to the effect of nuclear localized GCaMP6f, while not sacrificing kinetics as occurs with nuclear localization. While soma-targeting of membrane proteins such as optogenetic actuators has been done for many years (Baker et al., 2016; Forli et al., 2018; Greenberg et al., 2011; Pégard et al., 2017; Shemesh et al., 2017; Wu et al., 2013a) to decrease crosstalk in the context of single-cell precision optogenetics, this strategy has not been adapted for genetically encoded calcium indicators. We screened through a diversity of peptides, both natural and engineered, and discovered two such small motifs that, when fused to GCaMP6f, enabled it to express primarily within 50 microns of the cell body. The kinetics of response were similar to those mediated by conventional GCaMP6f. We found that in intact brain circuits, such as in living larval zebrafish and mice, these soma-targeted GCaMP6f molecules were able to greatly reduce the number of neuropil contamination spikes mistakenly attributed to a given neural cell body. Because of these effects, use of soma-targeted GCaMP6f greatly reduced artifactual correlations between nearby neurons in live zebrafish and mouse brain. Thus soma-targeted calcium indicators may be useful in a diversity of situations where high speed one-photon calcium population neuron imaging is desired.

## Results

### Designing and screening cell-body targeted GCaMP6f variants

As a test case to realize the strategy of cell body targeting of genetically encoded calcium indicators, we chose GCaMP6f, which is currently popular due to its high calcium sensitivity and ability to report single action potentials (Chen et al., 2013). We first searched the literature for proteins known to express somatically. We chose 8 such proteins (see **Supplemental Table 1** for a list of the proteins, as well as the various fragments and fusions, and **Supplemental Table 13** for the sequences of the fragments) for further consideration. These were the kainate receptor subunit KA2 (Shemesh et al., 2017; Valluru et al., 2005), the potassium channel K_V_2.1 (Lim et al., 2000), the sodium channels Na_V_1.2 and Na_V_1.6 (Garrido et al., 2003), the adaptor protein Ankyrin_G_ (Zhang and Bennett, 1998), and the rat small conductance calcium-activated potassium channel rSK1 (Bowden et al., 2001). In addition, we explored de novo designed coiled-coil proteins that self-assemble into complexes, hypothesizing that their mutual binding could potentially slow their diffusion from the cell body; interestingly, two of these self-assembling protein fragments, EE-RR (Moll et al., 2001; Selgrade et al., 2013) and AcidP1-BaseP1 (Oakley and Kim, 1998), did indeed (see below) result in somatic localization, raising the possibility that such fundamental protein engineering building blocks might find applicability in neuroengineering.

For some of these soma-restricted proteins, in earlier work cell body expression was analyzed by fusing the full-length proteins to reporters – specifically, Na_V_1.2, Na_V_1.6, Ankyrin_G_, and rSK1 were fused to fluorescent proteins (FPs) (Garrido et al., 2003; Moruno Manchon et al., 2015; Schäfer et al., 2010; Zhang and Bennett, 1998), KA2 to a Myc-tag (Valluru et al., 2005), and K_V_2.1 to an HA-tag (Lim et al., 2000). In some cases, earlier work showed that key fragments were sufficient to cause soma targeting of a reporter (**Supplemental Table 1**). For Na_V_1.2 and Na_V_1.6, 326- and 27-amino acid segments within intracellular loops between transmembrane domains, termed Na_V_1.2(I-II) and Na_V_1.6(II-III) respectively (see **Supplemental Table 13** for sequences), were sufficient for somatic localization (Garrido et al., 2001, 2003). For K_V_2.1, a 65–amino acid segment within the intracellular loop between transmembrane domains IV and V (K_V_2.1-motif, see **Supplemental Table 13** for sequences) sufficed (Lim et al., 2000; Wu et al., 2013b). For rSK1, the tail region (rSK1-tail, see **Supplemental Table 13** for sequences) sufficed (Fletcher et al., 2003). For Ankyrin_G_ it was found that the spectrin-binding domain (AnkSB-motif, see **Supplemental Table 13** for all Ankyrin subsequences), the tail domain (AnkTail-motif), the membrane-binding domain (AnkMB-motif), the COOH-terminal domain (AnkCT-motif) and the serine-rich domain (AnkSR-motif) were all targeted to the axon and the cell body of neurons (Zhang and Bennett, 1998).

We made over 30 fusions between GCaMP6f and the protein fragments reported above (see the different fusions screened in **Supplemental Table 2** and the sequences of localization fragments in **Supplemental Table 13**). For Na_V_1.2, Na_V_1.6, K_V_2.1, and rSK1 we performed fusions in which the previously characterized localization fragment was attached to the C-terminus of GCaMP6f. In a recent study (Shemesh et al., 2017), we fused the channelrhodopsin CoChR (Klapoetke et al., 2014) to the first 150 amino acids of the KA2 receptor subunit (KA2(1-150)) thereby creating a somatic CoChR. Since both N and C terminal fusions of KA2(1-150) with CoChR caused somatic localization, we made similar upstream and downstream fusions of this fragment with GCaMP6f (**Supplemental Table 2**). In the present study, we also found that the first 100 amino acids of KA2 (KA2(1-100)) were sufficient to introduce somatic localization of GCaMP6f, therefore we made additional upstream and downstream fusions of KA2(1-100) with GCaMP6f (**Supplemental Table 2**). Since the length of the linker between parts of a fusion protein can affect the ultimate function of the fusion, we tested the effect of different linker lengths between GCaMP6f and trafficking sequences on soma localization (**Supplemental Table 2**). In some cases, we inserted into the construct a superfolder GFP (sfGFP; (P?delacq et al., 2006), which contains three mutations to EGFP in order to enhance folding), with a mutation to abolish its fluorescence (here called nullsfGFP, see STAR Methods for the full sequence of nullsfGFP; **Supplemental Table 2**). This was done to explore whether better folding, facilitated by sfGFP, might help improve expression of the final fusion protein. For Ankyrin_G_ fragments, we made fusions both upstream and downstream of GCaMP6f (**Supplemental Table 2**). For *de-novo* coiled-coil proteins, we made downstream fusions only.

We expressed each of these GCaMP6f fusion protein in cultured mouse hippocampal neurons (**Supplemental Table 2**). Thereafter, using wide-field fluorescence microscopy we performed a preliminary screen to sort through the fusions and prioritize them for more detailed characterizations. In this screen, we assessed the expression level (fluorescence under baseline conditions), the somatic localization of the GCaMP6f fluorescence, the toxicity (assessed as the percentage of dead fluorescent cells out of all expressing cells), and whether there was a fluorescent change over the baseline fluorescence (termed here df/f_0_, see STAR Methods for explanation of calculation) indicative of spontaneous neural activity. We found that five constructs did not result in obvious toxicity, exhibited somatic localization, and displayed dynamic activity with a df/f_0_ similar to that of GCaMP6f (**Supplemental Table 2**). These were GCaMP6f fused to the fragments mentioned below (integers in the construct names denote the length of the linker; see **Supplemental Table 2** for fusions tested and for the sequences of different linkers): Na_V_1.2(I-(GCaMP6f-27-Na_V_1.2(I-II)-ER2); GCaMP6f fused upstream to nullsfGFP and to KA2(1-100) (GCaMP6f-24-nullsfGFP-24-KA2(1-100)-ER2); GCaMP6f fused downstream to a zero-photocurrent CoChR mutant called nullCoChR followed by the K_V_2.1-motif (nullCoChR-12-GCaMP6f-K_V_2.1-motif); GCaMP6f fused to AnkTail-motif (GCaMP6f-27-AnkTail-motif-ER2); and finally GCaMP6f fused to the coiled-coil peptide set EE-RR (GCaMP6f-27-EE-RR).

We screened these five somatic GCaMP6f candidates for expression in mouse brain circuitry, incubating mouse cortical slices expressing these five candidates with 4-aminopyridine (4-AP) to induce a low level of spiking to screen for physiological function (see STAR Methods). 1 mM 4-AP resulted in repeated transient upshoots in GCaMP6f brightness, with approximately 5-20 GCaMP transients occurring per minute in neurons in such a slice (**Figure S1**), which we estimated to represent spikes, and therefore we refer to such transients abbreviatedly as GCaMP-spikes from here onward; essentially no GCaMP-spikes were seen in slices not exposed to 4-AP. For each somatic GCaMP6f candidate, we assessed the df/f_0_, calculating the ratio between the df/f_0_ at the cell body and the df/f_0_ in the neuropil (see STAR Methods for how we determined somata vs. neuropil in mouse brain slices). We set as our screen criteria that a good somatic GCaMP6f would have a df/f_0_ similar to or larger than conventional GCaMP6f at the soma, and also exhibit a ratio of soma df/f_0_ to neuropil df/f_0_ larger than non-targeted GCaMP6f. The latter ratio we used as a measure, during this screen, of soma-localization, reasoning that for a soma-localized GCaMP6f, the neuropil df/f_0_ would begin to get lost in the noise; while not precise, our intention at this phase of the project was simply to do a fast screen in brain slices, and measuring exact falloff of fluorescence along neurites is hard to measure without actually tracing the neurites. We found that GCaMP6f-24-nullsfGFP-24-KA2(1-100)-ER2 expressed in the neurites of pyramidal neurons in the cortex, indicating impaired somatic localization when tested in an in vivo context, and we did not further pursue this construct in **Fig. S1** or beyond. The remaining four constructs had a soma df/f_0_ to neuropil df/f_0_ significantly higher than that of GCaMP6f (**Figure S1** and **Supplemental Table 7**). GCaMP6f-27-AnkTail-motif-ER2 and GCaMP6f-27-EE-RR had the highest df/f_0_ at the soma (103±13% and 135±29% respectively; mean ± standard deviation reported throughout; n = 20 cells from 2 slices from 2 mice for each SomaGCaMP6f variant, and GCaMP6f **Figure S1**). Because of their high sensitivity, we chose to pursue these two for more detailed characterization. We named these two constructs SomaGCaMP6f1 and SomaGCaMP6f2, respectively.

### Characterization of SomaGCaMP6f variants in mouse hippocampal cultures

We co-expressed GCaMP6f, GCaMP6f-27-AnkTail-motif-ER2 (SomaGCaMP6f1) or GCaMP6f-27-EE-RR (SomaGCaMP6f2) with the red fluorescent protein mCardinal to serve as a cellular tracer, using cultured mouse hippocampal neurons (**Figure 1C-K**). In cultured neurons, the number of transfected neurons was sparse (approximately one transfected neuron per 200 non-transfected neurons) and therefore we were able to trace single neurites. Thus, we traced neurites and looked at fluorescence as a function of distance down each neurite in individual cells. We found that the fluorescence fell off in neurites much faster in SomaGCaMP6f1 (**Figure 1L,M**) and SomaGCaMP6f2 (**Figure 1L,N**) expressing cells compared to GCaMP6f expressing cells (see **Supplemental Table 3** for full statistics). The baseline fluorescence of GCaMP6f, SomaGCaMP6f1, and SomaGCaMP6f2 expressing cells in culture were all similar to each other, and to that of the nuclear localized GCaMP6f-NLS (**Figure 2A**, see **Supplemental Table 13** and STAR Methods for the nuclear localization sequence). We next compared the fluorescent response of each molecule to a single action potential in cultured hippocampal neurons (**Figure 2B**), and found comparable responses (**Figure 2C**; see **Supplemental Table 4** for full statistics). We found that SomaGCaMP6f1 and SomaGCaMP6f2 had an SNR (defined as the magnitude of the fluorescence change caused by a single action potential divided by the standard deviation of the baseline fluorescence) similar to GCaMP6f, whereas GCaMP6f-NLS had an SNR significantly lower than that of GCaMP6f (**Figure 2D**; see **Supplemental Table 4** for full statistics). We found that SomaGCaMP6f1 and SomaGCaMP6f2 had rise (τ_on_) and decay (τ_off_) times, for a single action potential, similar to those of GCaMP6f, and that, as expected from previous work, GCaMP6f-NLS had rise and decay times significantly slower than those of GCaMP6f (**Figure 2E, F** and **Supplemental Table 4** for full statistics). When we analyzed the resting potential, membrane capacitance, holding current, and membrane resistance of cultured hippocampal neurons, they did not differ for cells expressing SomaGCaMP6f1 or SomaGCaMP6f2 vs. GCaMP6f (**Figure S2**).

**Figure 2.**
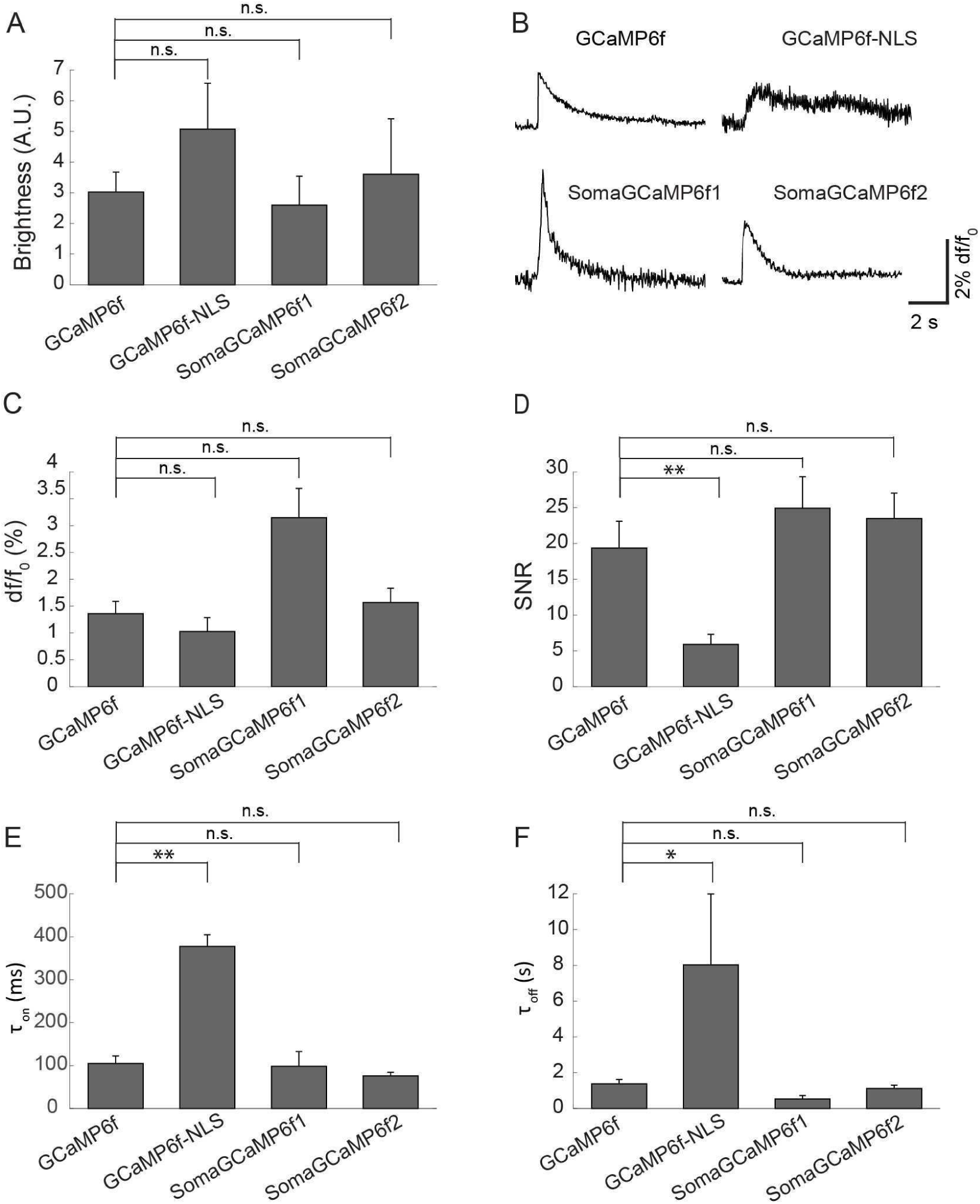
Kinetics and sensitivity of SomaGCaMP6f1 and SomaGCaMP6f2, as compared to conventional and nuclear-targeted GCaMP6f. GCaMP6f, GCaMP6f-NLS, SomaGCaMP6f1 and SomaGCaMP6f2 were transfected into hippocampal neurons for patch clamp and imaging. (**A**) Average baseline brightness values for GCaMP6f, GCaMP6f-NLS, SomaGCaMP6f1 and SomaGCaMP6f2 (n = 8 cells from 2 cultures for GCaMP6f; n = 7 cells from 2 cultures for SomaGCaMP6f1; n = 5 cells from 2 cultures for SomaGCaMP6f2; n = 7 cells from 2 cultures for GCaMP6f-NLS). n.s., not significant, Kruskal-Wallis analysis of variance followed by post-hoc test via Steel’s test with GCaMP6f as control group; see **Supplemental Table 4** for full statistics for **Figure 2**. (**B**) A representative fluorescence response for one action potential in the cell body for GCaMP6f, GCaMP6f-NLS, SomaGCaMP6f1 and SomaGCaMP6f2. (**C**) df/f_0_ for GCaMP6f, GCaMP6f-NLS, SomaGCaMP6f1 and SomaGCaMP6f2 (n = 8 cells from 2 cultures for GCaMP6f; n = 5 cells from 2 cultures for SomaGCaMP6f1; n = 7 cells from 2 cultures for SomaGCaMP6f2; n = 8 cells from 2 cultures for GCaMP6f-NLS). n.s., not significant, Kruskal-Wallis analysis of variance followed by post-hoc test via Steel’s test with GCaMP6f as control group; see **Supplemental Table 4** for full statistics for **Figure 2**. (**D**) Signal to noise ratio (SNR), defined as the magnitude of the fluorescence change caused by a single action potential divided by the standard deviation of the baseline fluorescence, for GCaMP6f, GCaMP6f-NLS, SomaGCaMP6f1 and SomaGCaMP6f2 (n = 8 cells from 2 cultures for GCaMP6f; n = 5 cells from 2 cultures for SomaGCaMP6f1; n = 7 cells from 2 cultures for SomaGCaMP6f2; n = 8 cells from 2 cultures for GCaMP6f-NLS). **P < 0.01, Kruskal-Wallis analysis of variance followed by post-hoc test via Steel’s test with GCaMP6f as control group; see **Supplemental Table 4** for full statistics for **Figure 2**. (**E**) Time constant for signal rise (Ton) during a single action potential for GCaMP6f, GCaMP6f-NLS, SomaGCaMP6f1 and SomaGCaMP6f2 (n = 8 cells from 2 cultures for GCaMP6f; n = 5 cells from 2 cultures for SomaGCaMP6f1; n = 6 cells from 2 cultures for SomaGCaMP6f2; n = 8 cells from 2 cultures for GCaMP6f-NLS). **P < 0.01, Kruskal-Wallis analysis of variance followed by post-hoc test via Steel’s test with GCaMP6f as control group; see **Supplemental Table 4** for full statistics for **Figure 2**. (**F**) Time constant for signal decay (Toff) after a single action potential for GCaMP6f, GCaMP6f-NLS, SomaGCaMP6f1 and SomaGCaMP6f2 (n = 7 cells from 2 cultures for GCaMP6f; n = 5 cells from 2 cultures for SomaGCaMP6f1; n = 7 cells from 2 cultures for SomaGCaMP6f2; n = 8 cells from 2 cultures for GCaMP6f-NLS). *P < 0.05, Kruskal-Wallis analysis of variance followed by post-hoc test via Steel’s test with GCaMP6f as control group; see **Supplemental Table 4** for full statistics for **Figure 2**.

### SomaGCaMP6f1 enables low-crosstalk imaging of neural activity in brain slices

We assessed whether soma targeting of GCaMP6f could reduce neuropil contamination by comparing patch-reported spikes to GCaMP-reported spikes in mouse brain slices. We randomly chose SomaGCaMP6f1 for this experiment; later we explored the use of SomaGCaMP6f2 in living mouse brain (see below). The idea was to patch cells in brain slices and electrophysiologically record from them while simultaneously imaging the cell bodies, so that we could count how many fluorescent GCaMP6f-reported spikes were detected in the cell body in the absence of corresponding patch-reported action potentials, and thus were the result of neuropil contamination. Using identical imaging parameters for histological analysis (see STAR Methods), we measured the density of labeled cells. Although SomaGCaMP6f1 is dimmer than GCaMP6f in the living brain (see below), we were able to easily identify the cells with expression and count them. We found that slices expressing either GCaMP6f or SomaGCaMP6f1 contained cells expressing the indicators at a density of 18±7 cells per 10^6^ µm^3^ and 21±5 cells per 10^6^ µm^3^, respectively (mean +/- standard error of the mean; n = 3 slices from 3 mice for GCaMP6f; n = 3 slices from 3 mice for SomaGCaMP6f1; **Figure 3A, B** and **Supplemental Table 5** for full statistics). To be conservative in our analysis, we manually traced cell bodies and made sure the region of interest (ROI) for calcium trace analysis was inside the cell, to avoid choosing an ROI that contained the contour of the cell, thereby decreasing the probability of including GCaMP6f fluorescence coming from processes of other cells (see STAR Methods). We found that the baseline brightness of the cell body of SomaGCaMP6f1-expressing neurons was about 4.8-fold lower than that of GCaMP6f-expressing neurons in live brain slices (**Figure S3** and **Supplemental Table 9** for full statistics), indicating a potential difference in level of expression between the in vitro (**Fig. 2**) and in vivo (**Fig. 3**) contexts, not uncommon for genetically encoded reagents given the different transfection protocols, gene dosages, and cellular contexts. Thus, in order to compare GCaMP6f and SomaGCaMP6f1 to each other fairly, in terms of quality of somatic targeting, change in fluorescence (df/f_0_) and signal-to-noise (SNR), and crosstalk, we increased the excitation light power in SomaGCaMP6f1 experiments to match the baseline brightness to GCaMP6f slices (**Figure 3C** and **Supplemental Table 5** for full statistics), for all further experiments reported in **Figure 3**. In such conditions, we found that despite similar brightness of cell bodies, fluorescence in the neurites fell off significantly faster down the neurite for SomaGCaMP6f1 than for GCaMP6f (**Figure 3D**). We found that df/f_0_ of transients per single patch-reported spikes observed during 4-aminopyridine evoked activity was similar between GCaMP6f and SomaGCaMP6f1 expressing cells in such slices (**Figure 3E** and **Supplemental Table 5** for full statistics), while the df/f_0_ of the transient driven by a burst (50 Hz current injections of 500 pA, 5 ms, in trains of 5, 10, or 20 pulses) was significantly higher in SomaGCaMP6f1 vs. GCaMP6f expressing cells (**Figure S4** and **Supplemental Table 10** for full statistics). The SNR for imaged action potentials was also comparable between GCaMP6f- and SomaGCaMP6f1-expressing neurons in slice (**Figure 3F** and **Supplemental Table 5** for full statistics). We compared the amount of crosstalk, as indicated by the number of fluorescent GCaMP-reported spikes that lack an associated patch-reported spike in brain slices expressing GCaMP6f vs. SomaGCaMP6f1 (**Figure 3G** vs. **Figure 3H**, respectively). Patch-reported spike rates were similar between GCaMP6f- and SomaGCaMP6f1-expressing cells in slices under the low-frequency spiking induced by 4-aminopyridine (**Figure 3I** and **Supplemental Table 5** for full statistics). Low-frequency spiking allowed detailed counting of spikes in the fluorescence vs. patch-reported traces. We found under such conditions that neurons in the GCaMP6f slices exhibited a roughly 2:3 ratio of erroneous spikes to actual spikes, meaning that for every three GCaMP-spikes that were corroborated by patch-reported APs, there were two erroneous GCaMP6f spikes. In contrast, in SomaGCaMP6f1 slices the ratio was cut to 1:6 (**Figure 3J**), a 75% decrease in artifact ratio.

**Figure 3.**
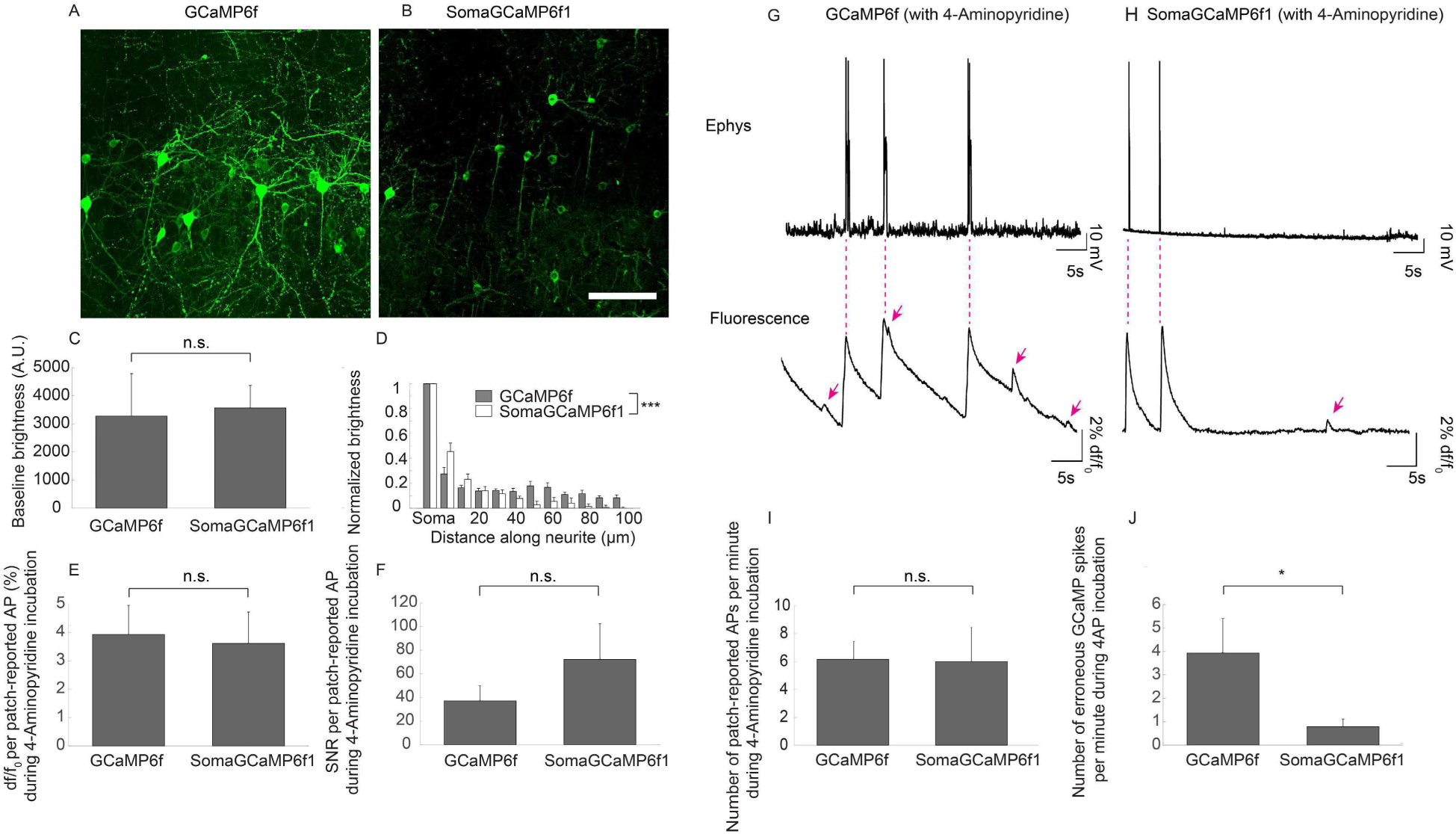
Decreased neuropil crosstalk in mouse brain slices expressing SomaGCaMP6f1. (**A, B**) Representative slices expressing GCaMP6f (**A**) and SomaGCaMP6f1 (**B**). Scale bar, 100 µm. (**C**) A bar chart showing average baseline brightness values for cells expressing GCaMP6f or SomaGCaMP6f1 in brain slice, following light power tuning so that the baseline recorded brightness from GCaMP6f or SomaGCaMP6f1 slices were similar (n = 7 neurons from 2 slices from 2 mice for GCaMP6f; n = 22 neurons from 6 slices from 3 mice for SomaGCaMP6f1). n.s., not significant, Wilcoxon rank sum test of the brightness between GCaMP6f and SomaGCaMP6f1; see **Supplemental Table 5** for full statistics for **Figure 3**. (**D**) A bar plot of brightness versus position along a neurite, normalized to brightness at the soma, extracted from neurites of neurons from slices expressing GCaMP6f or SomaGCaMP6f1 (n = 11 neurites from 6 neurons from 2 slices from 2 mice for GCaMP6f; n = 11 neurites from 6 neurons from 2 slices from 2 mice for SomaGCaMP6f1). ***P < 0.001, Kolmogorov-Smirnov test of neurite brightness between GCaMP6f and SomaGCaMP6f1; see **Supplemental Table 5** for full statistics for **Figure 3**. (**E**) A bar chart showing the average df/f_0_ of somata of neurons in slices expressing GCaMP6f or SomaGCaMP6f1 during an action potential (n = 14 APs from 3 neurons from 3 slices from 2 mice for GCaMP6f; n = 6 APs from 3 neurons from 3 slices from 3 mice for SomaGCaMP6f1). n.s., not significant, Wilcoxon rank sum test of the df/f_0_ between GCaMP6f and SomaGCaMP6f1; see **Supplemental Table 5** for full statistics for **Figure 3**. (**F**) A bar chart showing the average signal to noise ratio (SNR) of somata of neurons in slices expressing GCaMP6f or somaGCaMP6f1 following an action potential (n = 14 APs from 3 neurons from 3 slices from 2 mice for GCaMP6f; n = 6 APs from 3 neurons from 3 slices from 3 mice for SomaGCaMP6f1). n.s., not significant, Wilcoxon rank sum test of the SNR between GCaMP6f and SomaGCaMP6f1; see **Supplemental Table 5** for full statistics for **Figure 3**. (**G, top**) Representative electrophysiological recording of a cell expressing GCaMP6f in a slice, under 4-AP stimulation. (**G, bottom**) The GCaMP6f fluorescent signal in the cell recorded in G, top. Magenta arrows denote peaks in the GCaMP fluorescent signal that do not have a corresponding patch-reported action potential (AP). (**H, top**) representative electrophysiological recording of a cell expressing SomaGCaMP6f1 in a slice, under 4-AP stimulation. (**H, bottom**) The SomaGCaMP6f1 fluorescent signal in the cell recorded from in H, top. Magenta arrows denote peaks in the SomaGCaMP6f1 fluorescent signal that do not have a corresponding action potential. (**I**) A bar chart showing the average number of patch-reported APs per minute in neurons in slices expressing GCaMP6f or somaGCaMP6f1 following an action potential (n = 8 neurons from 8 slices for GCaMP6f from 4 mice; n = 6 neurons from 6 slices for SomaGCaMP6f1 from 3 mice). n.s., not significant, Wilcoxon rank sum test of the average number of APs per minute between GCaMP6f and SomaGCaMP6f1; see **Supplemental Table 5** for full statistics for **Figure 3**. (**J**) A bar chart showing the number of erroneous GCaMP-spikes per minute in neurons expressing either GCaMP6f or SomaGCaMP6f1 in slice (n = 8 neurons from 8 slices from 4 mice for GCaMP6f; n = 6 neurons from 6 slices from 3 mice for SomaGCaMP6f1). *P < 0.05, Wilcoxon rank sum test of the number of fluorescent peaks minus the number of APs between GCaMP6f and SomaGCaMP6f1 expressing neurons; see **Supplemental Table 5** for full statistics for **Figure 3**.

We measured the decay times of the fluorescent GCaMP spikes, using two different stimulation protocols. In the first, we electrophysiologically injected current into GCaMP6f- or SomaGCaMP6f1-expressing cells to induce single action potentials in single cells in brain slices. In the second stimulation protocol, we used 0.1 mM 4-aminopyridine to induce action potentials throughout the slice. The τ_off_ was similar for GCaMP6f and SomaGCaMP6f1 for single action potentials evoked by electrophysiology (**Figure S4** and **Supplemental Table 10** for full statistics). For 4-aminopyridine evoked APs, GCaMP6f-expressing cells showed increased τ_off_ compared to the values recorded after electrophysiology-evoked spikes. We found that the calcium spike rate was similar between GCaMP6f and SomaGCaMP6f1 expressing slices (10.4±2.2 GCaMP-spikes per minute for GCaMP6f and 6.7±3.0 GCaMP-spikes per minute for SomaGCaMP6f1; mean +/- standard error of the mean; n = 8 neurons from 8 slices from 4 mice for GCaMP6f; n = 6 neurons from 6 slices from 3 mice or SomaGCaMP6f1; **Figure S4D** and **Supplemental Table 10** for full statistics).

### SomaGCaMP6f1 reduces crosstalk between neurons in larval zebrafish brain

To assess the performance of GCaMP6f vs. SomaGCaMP6f variants in zebrafish, we first transiently and sparsely expressed these molecules in the brains of larval zebrafish by DNA injection into embryos at 1–2 cell stages (**Figure 4A**). We observed no transient expression of SomaGCaMP6f2 in the injected fish, in initial studies, so for our immediate purposes we chose to compare GCaMP6f vs. SomaGCaMP6f1 in zebrafish.

**Figure 4.**
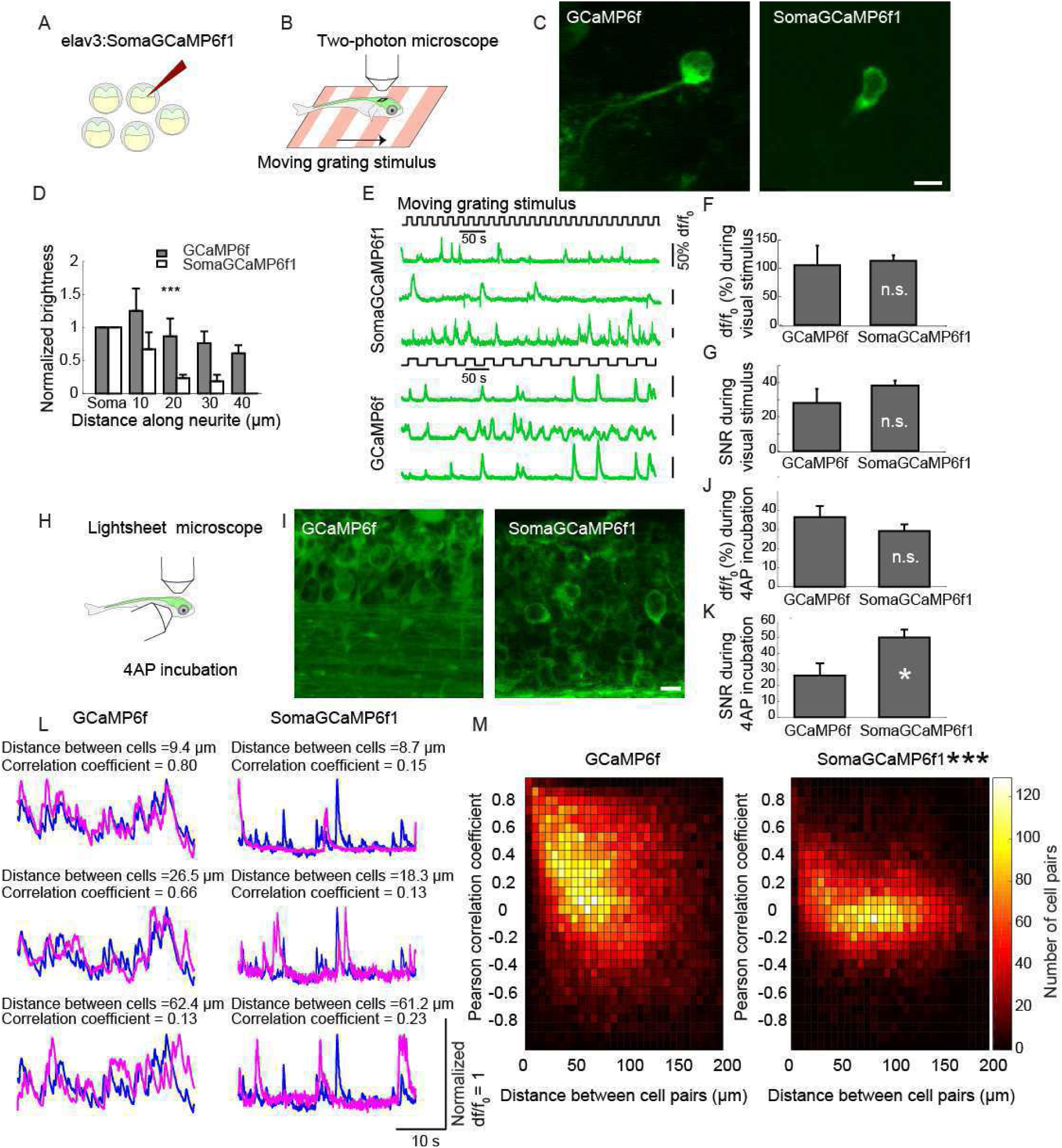
Decreased neuropil crosstalk in SomaGCaMP6f1-expressing larval zebrafish. (**A**) Embryos (1-2 cell stage) were injected with 20 ng/μl of elavl3:SomaGCaMP6f1. (**B**) Fish exhibiting transient expression in the brain were selected and imaged under the 2-photon microscope. A forward moving grating was used as a stimulus as GCaMP6f or SomaGCaMP6f1 expressing cells were imaged at 15 Hz. For the two-photon experiments (panels E, F, and G): for GCaMP6f experiments, 20s on / 20s off stimulus periods were used; for SomaGCaMP6f1, 10s on / 10s off (the difference in frequencies between GCaMP6f and SomaGCaMP6f1 was inadvertent). (**C**) Representative images of neurons transiently expressing GCaMP6f and SomaGCaMP6f1 in zebrafish larvae at 5 dpf. Scale bar: 5µm. (**D**) Bar plot of GCaMP6f brightness versus position along a neurite, normalized to GCaMP6f brightness at the soma, extracted from neurites of zebrafish neurons expressing GCaMP6f (n = 5 neurites in 4 cells in 2 fish) and SomaGCaMP6f1 (n = 8 neurites in 8 cells in 4 fish). ***P < 0.001, Wilcoxon rank sum test of neurite brightness between GCaMP6f and SomaGCaMP6f1; see **Supplemental Table 6** for full statistics for **Figure 4**. (**E**) Representative calcium traces for SomaGCaMP6f1-expressing cells and for GCaMP6f-expressing cells in response to the moving grating. (**F**) Bar chart showing the average df/f_0_ of somata of neurons in the optic tectum of zebrafish expressing GCaMP6f or SomaGCaMP6f1 in response to the moving grating (n = 6 neurons from 3 fish for GCaMP6f; n = 5 neurons from 3 fish for SomaGCaMP6f1). n.s., not significant, Wilcoxon rank sum test of the df/f_0_ between GCaMP6f and SomaGCaMP6f1; see **Supplemental Table 6** for full statistics for **Figure 4**. (**G**) Bar chart showing the average signal-to-noise ratio (SNR, see definition in STAR Methods) of somata of neurons in the optic tectum of zebrafish expressing GCaMP6f or SomaGCaMP6f1 in response to the moving grating (n = 6 neurons from 3 fish for GCaMP6f; n = 5 neurons from 3 fish for SomaGCaMP6f1). n.s., not significant, Wilcoxon rank sum test of the SNR between GCaMP6f and SomaGCaMP6f1; see **Supplemental Table 6** for full statistics for **Figure 4**. (**H**) Fish exhibiting stable pan-neuronal expression in the brain were selected and imaged using a lightsheet microscope. 4-AP stimulation was used for the experiments described in panels J, K, L, and M. (**I**) A z-projection image of neurons expressing GCaMP6f (left) or SomaGCaMP6f1 (right) in the spinal cord of zebrafish. Scale bar: 5µm. (**J**) Bar chart showing the average df/f_0_ of calcium events in the somata of zebrafish neurons in the forebrain expressing GCaMP6f or SomaGCaMP6f1 and stimulated with 4-AP (n = 5 neurons from 2 fish for GCaMP6f; n = 5 neurons from 2 fish for SomaGCaMP6f1). n.s., not significant, Wilcoxon rank sum test of the df/f_0_ between GCaMP6f and SomaGCaMP6f1. (**K**) Bar chart showing the average signal-to-noise ratio (SNR) of somata of zebrafish neurons in the forebrain expressing GCaMP6f or SomaGCaMP6f1 and stimulated with 4-AP (n = 5 neurons from 2 fish for GCaMP6f; n = 5 neurons from 2 fish for SomaGCaMP6f1). *P < 0.05, Wilcoxon rank sum test of the SNR between GCaMP6f and SomaGCaMP6f1; see **Supplemental Table 6** for full statistics for **Figure 4**. (**L**) Traces, normalized to their respective maxima for clarity, of representative cell pairs in the forebrain expressing GCaMP6f (left) or SomaGCaMP6f1 (right) that are ∼10 µm (top row), ∼20 µm (middle row) and ∼50 µm (bottom row) apart, during 4-AP stimulation. Pearson correlation coefficients between the traces are denoted above them. (**M**) A density plot showing the Pearson correlation coefficients of cell pairs in the forebrain as a function of the distance between cell pairs for GCaMP6f (n = 426 cells from 5 fish) and SomaGCaMP6f1 (n = 340 cells from 4 fish), during 4-AP stimulation. ***P < 0.001, two-dimensional Kolmogorov-Smirnov test between GCaMP6f and SomaGCaMP6f1; see **Supplemental Table 6** for full statistics for **Figure 4**.

We imaged neurons expressing either GCaMP6f or SomaGCaMP6f1 using two-photon microscopy (**Figure 4B,C**) and found that the fluorescent signal in neurites fell off significantly faster down neurites in SomaGCaMP6f1-expressing cells compared to GCaMP6f-expressing cells (**Figure 4D**). We imaged the tectum of the fish brain with a two-photon microscope while presenting a visual stimulus consisting of a moving grating (**Figure 4B**, see STAR Methods for description of the stimulus) and found that cells expressing GCaMP6f or SomaGCaMP6f1 exhibited fluorescent transients during the presentation of the visual stimulus (**Figure 4E**), and that the df/f_0_ and SNR measured at the cell bodies were similar in GCaMP6f and SomaGCaMP6f1-expressing fish (**Figure 4F,G**, see STAR Methods for calculation of df/f_0_ and SNR). For the following experiments, we generated stably expressing fish lines (see STAR Methods) expressing pan-neuronally. We imaged these fish with a one-photon lightsheet microscope (**Figure 4H**), and found that in GCaMP6f-expressing fish, GCaMP6f-filled neurites touched upon GCaMP6f-filled cell bodies, resulting in the kind of situation that could result in crosstalk, but in SomGCaMP6f1 fish this phenomenon was less pronounced, when both were evaluated for the same region of interest and analyzed with the same software package (Pnevmatatakis et al 2016) (**Figure 4I**). In these fish lines, as in mice (see above), we found that the baseline fluorescence of SomaGCaMP1-expressing cells was approximately 4.7 fold lower compared to GCaMP6f (n = 25 cells from 5 GCaMP6f fish, n = 25 cells from 5 SomaGCaMP6f1 fish, see **Supplemental Table 6** for values and statistics). For this reason, we increased the laser power approximately 4.5-5 fold in SomaGCaMP6f1 experiments (see STAR Methods), to cause similar brightness as GCaMP6f. To induce neural spiking, we immersed the fish in 4-aminopyridine and imaged their brains over 10-minute-long periods (see STAR Methods). The df/f_0_ for GCaMP6f and SomaGCaMP6f1 cells in the forebrain were similar, and the SNR for SomaGCaMP6f1 was twice that of GCaMP6f (**Figure 4J,K**). We then calculated the Pearson correlation coefficients between all the possible neuron pairs in the field of view (**Figure 4L**) and plotted them against the distance between these neural pairs (**Figure 4M**), to see whether crosstalk was more pronounced for nearby neurons in the GCaMP6f case. Indeed, we found that that in GCaMP6f-expressing brains, the shorter the distance between neuron pairs, the higher the correlation between their GCaMP-spikes, while in SomaGCaMP6f1 expressing neurons this dependency of pairwise correlation on the distance was significantly lower (**Figure 4M** and **Supplemental Table 6** for full statistics). This suggests that the artifactual contamination of cell body signals with neuropil signals can manifest as an artifactual increase in correlation between neural activity patterns, which could lead in turn to artifactual conclusions about neural connectivity, oscillatory dynamics, synchrony, and neural codes.

We found that for GCaMP-spikes, τ_on_ and τ_off_ were similar between GCaMP6f and SomaGCaMP6f1 fish (**Figure S5** and **Supplemental Table 11** for full statistics). We counted the number of GCaMP spikes in both GCaMP6f and SomaGCaMP6f1 fish, and detected approximately 3 times more GCaMP spikes in SomaGCaMP6f1 fish compared to GCaMP fish (**Figure S5** and **Supplemental Table 11** for full statistics).

### SomaGCaMP6f2 reduces crosstalk in brains of behaving mice

For in vivo mouse experiments, we expressed the two SomaGCaMP6f variants in the dorsal striatum of mice, which contains a homogenous population of densely packed medium spiny neurons, whose cell bodies are accessible to imaging. Recently, it has been suggested that medium spiny neurons form populations of clustered cells with highly correlated neural activity (Barbera et al., 2016), although the relative strength of this correlation remains controversial – in part due to questions about neuropil contamination (Klaus et al., 2017). We expressed SomaGCaMP6f1 and SomaGCaMP6f2 in the dorsal striatum of the living mouse brain, and found, as before, that the brightness of SomaGCaMP6f1 in-vivo was approximately 4.5 times lower compared than that of GCaMP6f (**Figure S6** and **Supplemental Table 12** for full statistics). SomaGCaMP6f2 had a similar brightness compared to GCaMP6f (**Figure S6** and **Supplemental Table 12** for full statistics). We compared imaged calcium activity patterns within the dorsal striatum between GCaMP6f- vs. SomaGCaMP6f2-expressing mice running on a spherical treadmill. In SomaGCaMP6f2-expressing mice we noted a substantial reduction in neuropil fluorescence as compared to GCaMP6f (**Figure 5A-B**). SomaGCaMP6f2 decay times were faster than GCaMP6f decay times (**Figure S6**). SomaGCaMP6f2 reported approximately 20% more calcium events then GCaMP6f (**Figure 5E**).

**Figure 5.**
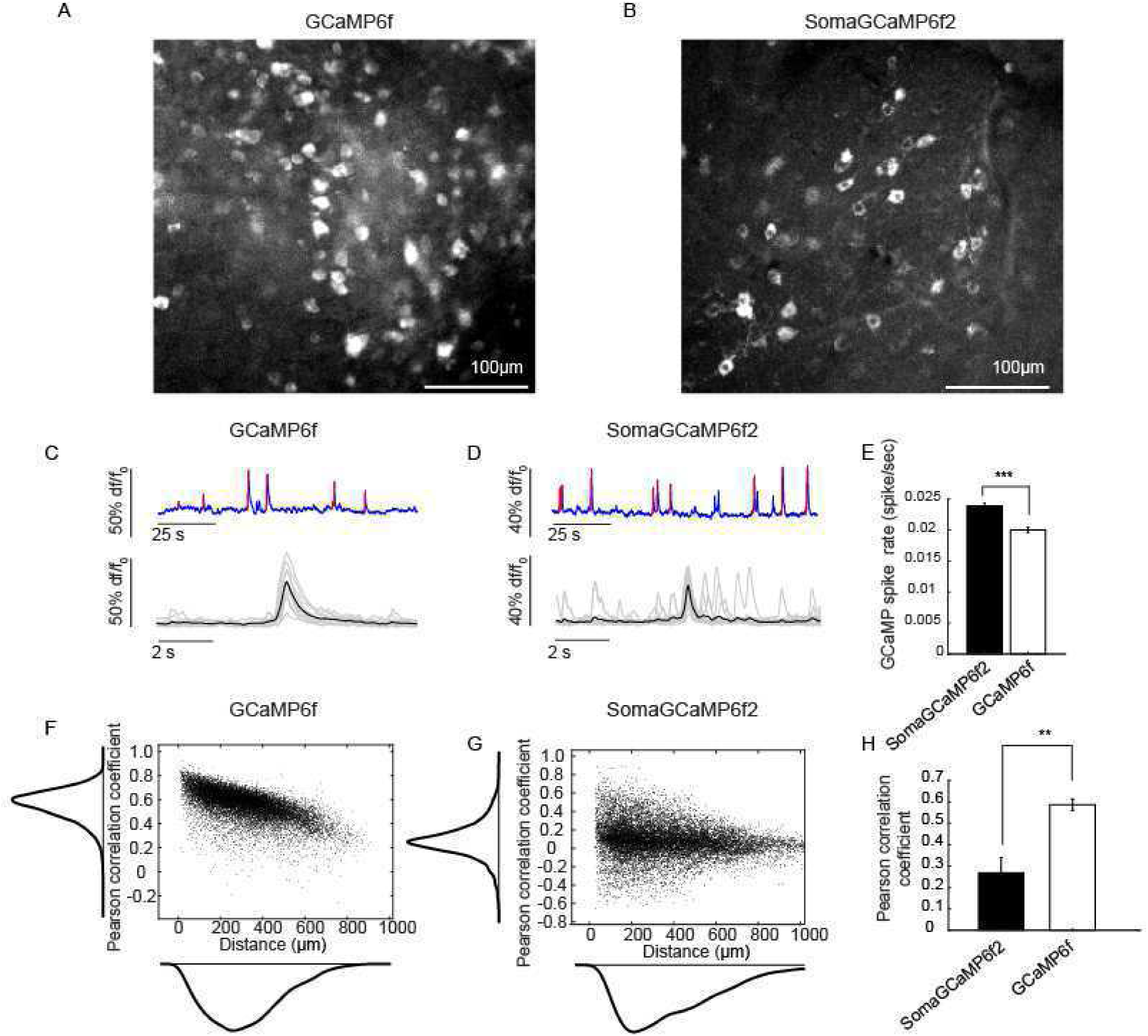
SomaGCaMP6f2 reduces neuropil contamination in the striatum of behaving mice. (**A, B**) Representative projection images showing the summed fluorescence, across all frames acquired in an imaging session (i.e., so that any neuron active at any time can be visualized), from the dorsal striatum in GCaMP6f- (**A**) or SomaGCaMP6f2- (**B**) expressing mice. Calcium imaging was performed using a 460 nm LED, with each imaging session lasting 5-12 minutes. (**C, D**) Representative calcium traces from two neurons shown in the images above that reflect GCaMP6f (**C**), or SomaGCaMP6f2 (**D**), fluorescence over a two minute (top) window. Normalized calcium traces are shown in blue as changes in df/f_0_. Calcium activation events were identified based on thresholding (see STAR Methods) and detected individual events are highlighted in red. Note that smaller events were not always detected using this methodology. Bottom: traces show calcium signals from the full session traces shown above, aligned to their peak amplitude. Individual events are shown in gray and their averaged response is shown in black. (**E**) Bar chart showing mean GCaMP-spike rates for neurons expressing either SomaGCaMP6f2 or GCaMP6f (n = 594 neurons from 4 mice expressing SomaGCaMP6f2, n = 930 neurons from 7 GCaMP6f mice). ***P <0.001, Wilcoxon rank sum test between the GCaMP-spike rates of SomaGCaMP6f2 and GCaMP6f expressing neurons; see **Supplemental Table 6** for full statistics for **Figure 5**. (**F, G**) Representative correlograms (from a single mouse each) denoting the relationship of distance to the strength of correlated fluorescence between ROIs from the sessions shown above for GCaMP6f (**F**) or SomaGCaMP6f2 (**G**). Distance distributions are shown on the x-axis and Pearson correlation coefficients are shown on the y-axis. (**H**) Bar chart showing the mean Pearson correlation coefficients from all SomaGCaMP6f2 or GCaMP6f mice (n = 176121 cell-pairs from 4 SomaGCaMP6f2 mice; n = 431985 cell-pairs from 7 GCaMP6f mice). **P < 0.01, Wilcoxon rank sum test between SomaGCaMP6f2 and GCaMP6f; see **Supplemental Table 6** for full statistics for **Figure 5**.

We calculated the Pearson correlation coefficients between all the possible neuron pairs within our imaging field. Representative correlograms from two mice expressing either GCaMP6f or SomaGCaMP6f2 are shown in **Figure 5F** and **Figure 5G** respectively. Within the striatum of GCaMP6f-expressing mice, we identified high correlations for nearby cells, that fell off with increasing distance. In contrast, SomaGCaMP6f2-expressing mice had far lower correlations across the board; we even found instances of strong negative correlations that were not present in GCaMP6f mice. Across the population, neurons were approximately 50% less correlated when expressing SomaGCaMP6f2 then with GCaMP6f (**Figure 5H**). Taken together our results reveal that the correlated activity of calcium signals is likely overrepresented, particularly for neurons close together in space.

## Discussion

We here report that it is possible to target genetically encoded calcium sensors to cell bodies in multiple species in vivo. The variants we focused on, SomaGCaMP6f1 and SomaGCaMP6f2, demonstrated satisfactory brightness, sensitivity, and kinetics in mouse and zebrafish brain. We observed decreased crosstalk, as reflected by lower numbers of artifactual (e.g., not patch pipette detectable) spikes, and reduced artifactual correlation between neurons that are nearby in both zebrafish and mouse brain. Although nuclear localized GCaMP can also achieve isolation between neurons, the slow speed has given pause to potential users; soma-targeting results in severalfold higher signal-to-noise ratio and severalfold faster kinetics, compared to nuclear GCaMP.

Having fewer artifactual spikes increases the accuracy of assessment of neural codes in the living brain. Examples of such experiments include: neurons expressing GCaMP6f in the CA1 area of the hippocampus of mice were imaged using single-photon microscopy during a trace-conditioning task (Mohammed et al., 2016); in the visual cortex of mice, the GCaMP6f spiking of neurons that responded to a flashing stimulus were recorded (Kim et al., 2016); in the dorsal horn of mice, different neural populations were shown to respond to different tactile stimuli (Sekiguchi et al., 2016); in zebrafish, neurons that control turning during swimming were identified using whole-brain light-sheet calcium imaging. SomaGCaMP6f variants could prove useful in the experiments above, since eliminating erroneous spikes could help experimenters better determine which neurons are contributing to a behavior, and how.

Reducing artifactual correlation may also help with studies of functional connectivity, where correlated neural activity has been used to infer functional connectivity in retina (Greschner et al., 2011), cortex (Alonso and Martinez, 1998), and many other systems. Single-photon calcium imaging has a speed advantage compared to two-photon imaging, and wide-field calcium imaging in mice (Modi et al., 2014) and fruit flies (Streit et al., 2016) is feasible and robust. The advantage of the SomaGCaMP6f variants in performing single-photon imaging in these model systems is that they enable separation of bona-fide physiological correlation from non-physiological correlation. Examples for such experiments include: in *Dropsphila melanogaster*, different compounds that decrease synchrony between cells were tested by calcium imaging in neurons (Streit et al., 2016); In experiments performed in zebrafish, pairwise correlations between neurons were used to cluster them (Sekiguchi et al., 2016).

## STAR Methods

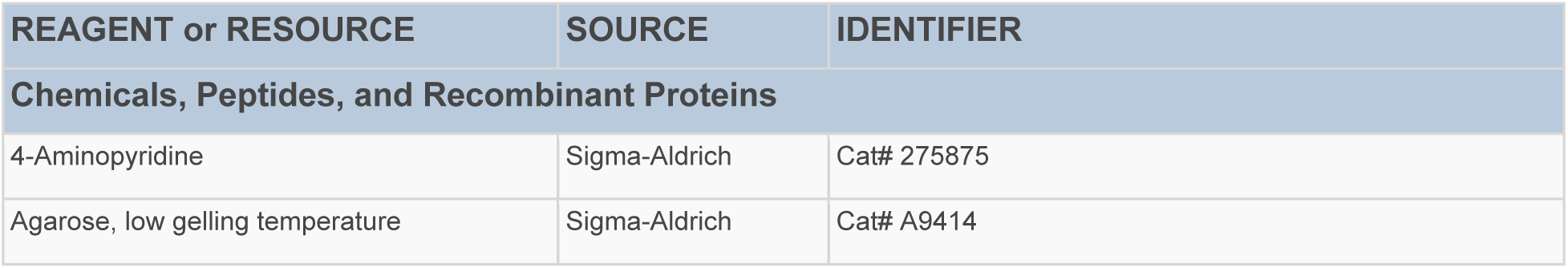
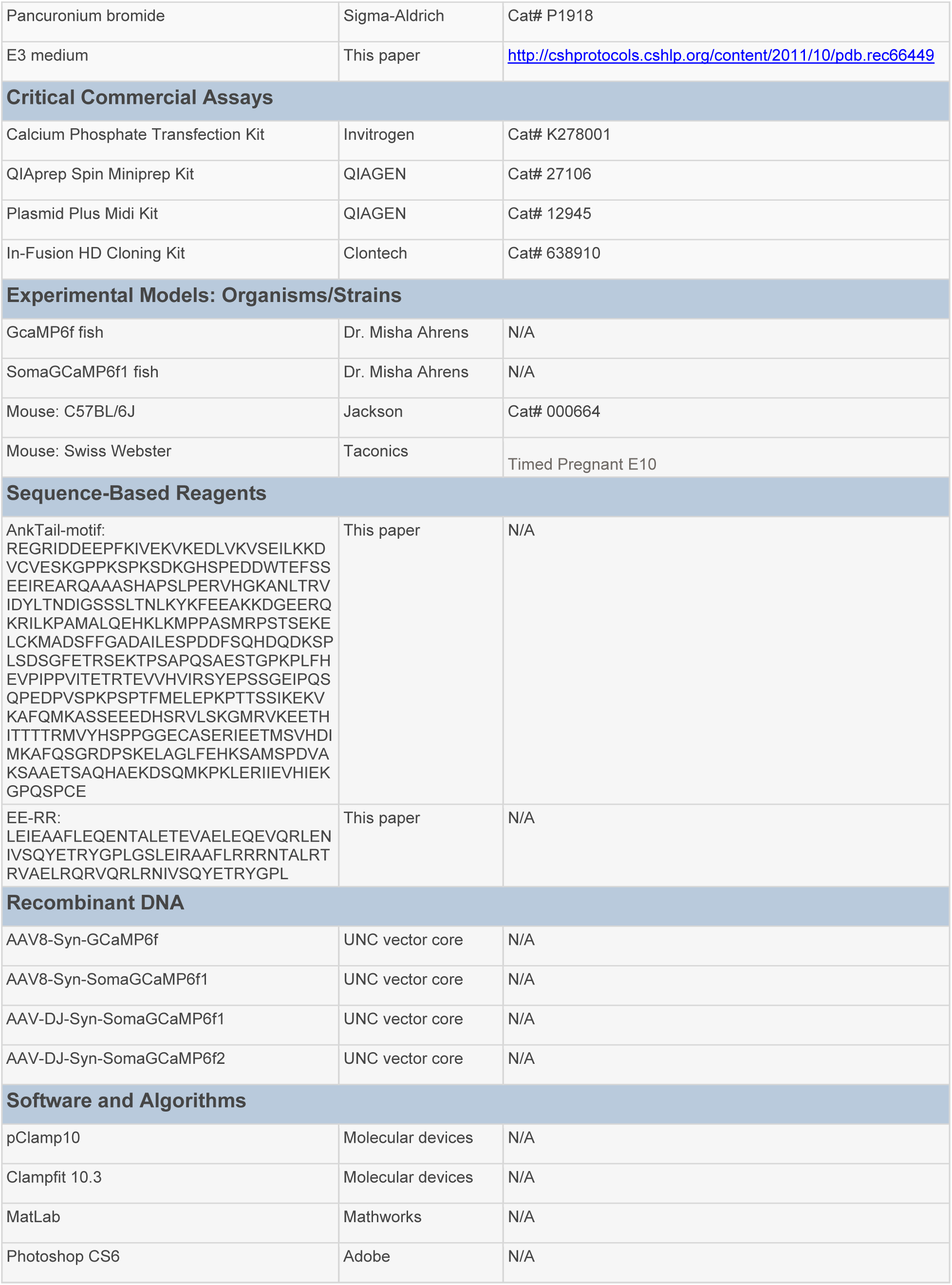
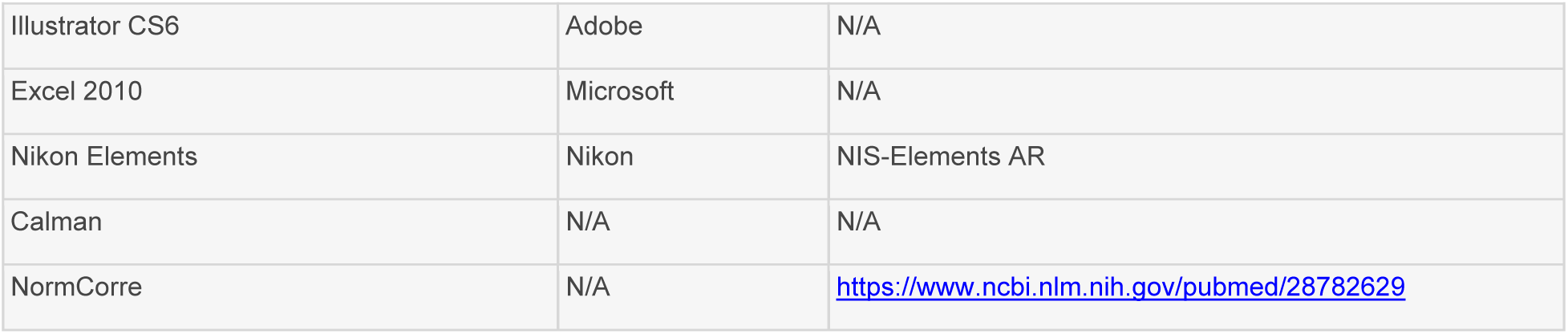

### Contact for Reagent and Resource Sharing

Further information and requests for reagents may be directed to, and will be fulfilled by, the Lead Contact and corresponding author, Dr. Ed Boyden.

### Experimental Model and Subject Details

#### Procedures

Procedures involving animals were in accordance with the National Institutes of Health Guide for the care and use of laboratory animals and approved by the Massachusetts Institute of Technology Animal Care and Use Committee. Zebrafish experiments at Janelia were conducted according to protocols approved by the Institutional Animal Care and Use Committee of the Howard Hughes Medical Institute, Janelia Research Campus. Zebrafish experiments at MIT were conducted according to protocols approved by the Institutional Animal Care and Use Committee of MIT. Hippocampal neuron culture was prepared from postnatal day 0 or day 1 Swiss Webster (Taconic) mice as previously described (Klapoetke et al., 2014). In-utero electroporation and subsequent slice work was performed on female Swiss Webster mice (Taconic).

### Zebrafish animals and trangenesis

For **Figure 4**, we used a previously published transgenic zebrafish line expressing GCaMP6f in the cytosol Tg(elavl3:GCaMP6f)jf1 (Freeman et al., 2014) The soma-localized GCaMP6f fish was generated as previously described (Freeman et al., 2014) using the Tol2 transposon system, in which indicators were subcloned into a Tol2 vector that contained the zebrafish elavl3 promoter. The transgene construct and transposase RNA were injected into 1–2-cell-stage embryos, and the transgenic lines were isolated by the high expression of bright green fluorescence in the central nervous system in the next generation. The larvae were reared in 14:10 light-dark cycles according to a standard protocol at 28.5° C, in a solution containing Instant Ocean salt from Carolina Biological Supply Company (65mg/L Instant Ocean, 30 mg/L Sodium bicarbonate). Experiments were performed on animals 5–7 days post fertilization (dpf) at room temperature. Fish lines and DNA constructs for elavl3:SomaGCaMP6f1 available upon request.

## Method Details

### Neural culture and transfection

For neuronal expression of GCaMP6f fusions with trafficking sequences during the screen for soma targeting sequences and for neuronal expression of mCardinal for **Figure 1** and **2**, we performed transfection at 4 days in vitro (DIV) with a commercial calcium phosphate kit (Invitrogen). We added an additional washing step with acidic MEM buffer (pH 6.8 – 6.9) after calcium phosphate precipitate incubation to completely re-suspend residual precipitates (Jiang and Chen, 2006). We used 1μg of DNA. Neurons were imaged 14–18 DIV (days in vitro; 10–14 days post-transfection).

### Gene synthesis

All genes were synthesized (by Epoch Life Science) with mammalian codon optimization and subcloned into pAAV backbone under CAG or Syn promoter, see **Supplemental Tables 1, 2 and 13** for descriptions and amino acid sequences. Briefly, for the final selected variants, 1200 bp from the tail region of the human AnkyrinG protein (Zhang and Bennett, 1998) (AnkTail-motif) were cloned followed by the ER2 (Hofherr et al., 2005) trafficking sequence from the potassium channel Kir2.1, with the resulting molecule being GCaMP6f-27-AnkTail-motif-ER2, named SomaGCaMP6f1, and 264 bp of a *de novo* designed coiled-coil peptide EE-RR fused to the C-terminus of GCaMP6f via a 27 amino acid flexible linker, named SomaGCaMP6f2. A nuclear localization sequence (NLS) was synthesized based on a sequence found in the literature (Kosugi et al., 2009).

### Image analysis

#### Analysis of GCaMP brightness along neurites, in cultured neurons, brain slices and zebrafish brains

Images for this analysis were taken for cultured neurons (**Figure 1**) at 14–18 DIV (10–14 days post-transfection), for live brain slices prepared as described below using mice at P12 – P24 (**Figure 3**), and for zebrafish larvae at 5-7 dpf (**Figure 4**). The image analysis was performed in ImageJ. For each neuron we first defined the boundaries of the soma. To that end, we drew a 20

µm diameter circle near the soma, inside which there was no apparent fluorescence from the soma or from neurites. We defined the average fluorescence in the circle as background fluorescence. We considered pixels with fluorescence intensity of at least 10% above background levels as part of the soma and processes, and we defined the boundary between soma and its processes by the apparent cell morphology. Then, we drew a polygon along the defined soma boundary and measured the average fluorescence inside of it, and subtracted the previously calculated background value. The resulting value was considered soma fluorescence. To measure fluorescence intensities along neurites, we defined 1µm^2^ rectangles along the neurite that were up to 100 µm away from soma at increments of 10 µm. The distance between each rectangle and the soma was measured along the neurites (not the minimal linear distance from the soma, since neurites were curved). We then defined the background value exactly as described above for the soma. We made sure that the pixel intensity values at the boundaries of the rectangle were at least 10% above background levels, to be considered inside the neurite. We averaged the fluorescence intensity in each rectangle, then subtracted the background, then divided it by the average soma fluorescence and plotted the resulting ratio with respect to distance along the neurite. The ratios for each distance were averaged across neurites and data was plotted (using Matlab) as average and standard error of the mean.

#### Analyzing brightness, df/f_0_, signal-to-noise ratio (SNR), fluorescent rise-time and fluorescence decay time following 1 action potential in-vitro

For **Figure 2**, hippocampal cells expressing the GCaMP6f trafficking variants were bathed with synaptic blockers (0.01 mM NBQX and 0.01 mM GABAzine) and patched (in current clamp), and at the same time images were acquired with a Hamamatsu Orca Flash 4.0 with an exposure of 20 ms. An action potential was elicited in the neuron using a 10 ms, 50-200 pA current injection, and the resulting fluorescence change was recorded for a period of 20 seconds, to allow the GCaMP6f fluorescence to return to baseline. To avoid sampling bias, we imaged and patched the first 2-3 cells detected according to the GCaMP fluorescence brightness in each plate. To calculate the GCaMP6f brightness at the soma of each cell, we defined the boundary of the soma by the apparent cell morphology in the image and subtracted the background fluorescence (as defined above) from the average fluorescence inside the soma boundary. To calculate df/f_0_ we first calculated baseline fluorescence. Baseline fluorescence was defined as the average fluorescence during the 1-second period right before the beginning of fluorescence response. df/f_0_ was calculated by dividing the maximum fluorescence change by baseline fluorescence. To calculate the signal to noise ratio (SNR) we divided the maximum fluorescence change by the standard deviation of baseline fluorescence during the 1-second period right before the onset of a GCaMP-spike. We calculated τ_on_ by extracting the time constant from the exponential fit of the rising segment of the fluorescence response. We calculated τ_off_ by extracting the time constant from the exponential fit of the falling segment of the fluorescence.

#### Measuring df/f_0_ and soma-to neuropil ratio in acute brain slices for SomaGCaMP variant screening

For **Figure S1**, regions of interest (ROIs) denoting cell bodies and neuropil were determined manually on a projection of the standard deviation of the fluorescence per pixel in the movies using ImageJ: twenty cells and one neuropil section were traced by hand using ImageJ’s freehand selection and ROI manager tools, from which 21 time histories of average fluorescence values F were extracted of length 2000 frames (40 seconds at 50Hz). The baseline fluorescence was defined as a 4-second time window with no apparent action potentials, from which we define *B* as the mean value in the baseline. For each neuron with, we define the df/f_0_ as 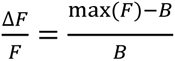. We next calculated soma to neuropil df/f_0_ ratio by dividing the soma df/f_0_ by the neuropil df/f_0._

#### Manually tracing cell bodies in slice patching and imaging crosstalk experiments in mouse brain slices

When tracing the cells (**Figure 3**, **Figure S4**), we chose a region of interest that was inside the cell body. We avoided choosing the ROI as the entire cell body, since that ROI may contain GCaMP6f filled processes originating from neighboring cells. We defined a cell body by the apparent cell morphology as was done as in the in vitro current clamp experiments. We then chose an ROI inside the cell body, approximately 1µm from the cell body’s apparent boundaries.

#### Analyzing brightness, df/f_0_ and signal-to-noise ratio (SNR) in acute slice patching experiments of GCaMP6f or SomaGCaMPf1

For **Figure 3** and **Figure S4**, we defined the boundary of the soma by the apparent cell morphology from the movies recorded in slice patching experiments, and measured the average fluorescence inside the soma boundary in each frame. To calculate df/f_0_ we first calculated baseline fluorescence. Baseline fluorescence was defined as the average fluorescence during the 100 to 500 ms period right before the beginning of fluorescence response. df/f_0_ was calculated by dividing the maximum fluorescence changes over baseline fluorescence in each cell body. To calculate the signal to noise ratio (SNR) we divided the maximum fluorescence change by the standard deviation of baseline fluorescence during the 100 to 500 ms period right before the onset of GCaMP-spikes.

#### Analyzing brightness, df/f_0_ and signal-to-noise ratio (SNR) in zebrafish larvae with either transient expression or stable pan-neuronal expression of GCaMP6f or SomaGCaMPf1

The movies recorded from zebrafish larvae with stable pan-neuronal expression using a lightsheet microscope (**Figure 4**) were first motion corrected using NormCorre (Pnevmatikakis and Giovanucci 2017). The movies recorded from zebrafish larvae with transient expression using a 2-photon microscope (**Figure 4**) were not motion corrected because little motion was observed. We defined the boundary of the soma by the apparent cell morphology from the movies, and measured the average fluorescence inside the soma boundary in each frame. To calculate df/f_0_ we first calculated baseline fluorescence. Baseline fluorescence was defined as the average fluorescence during the 1-second period right before the beginning of a fluorescence transient. df/f_0_ was calculated by dividing the maximum fluorescence change by baseline fluorescence in each cell body. To calculate the signal to noise ratio (SNR) we divided the maximum fluorescence change by the standard deviation of baseline fluorescence during the 1-second period right before the onset of a GCaMP-spike. To calculate correlation-coefficients between neuronal pairs in zebrafish larvae with stable pan-neuronal expression of GCaMP6f or SomaGCaMPf1, we processed the motion corrected movies with CaImAn (Pnevmatikakis et al 2016) to segment all putative neurons in the field of view, then denoised and deconvolved the fluorescence traces. An additional manual review was done for each candidate neuron from CaImAn to examine the spatial footprint and temporal characteristics to confirm it was a neuron. These filtered sets of neurons were then used for pairwise correlations of the denoised time signal and pairwise distance measurements using the centroid of the spatial footprints.

#### Analysis of in vivo calcium imaging data in live mice (for Figure 5)

##### a) Motion correction

Sessions varied between 5 and 12 minutes in length and imaging sessions were analyzed from four SomaGCaMP6f2 mice and seven GCaMP6f expressing mice. Motion correction was performed with a custom python script. For each imaging session, a reference image was generated by projecting the mean values of every pixel in the first file. The reference image and each frame of the video underwent a series of image processing steps to enhance the contrast and the character of the image. We first high-pass filtered the image with a Gaussian filter (python SciPy package, ndimage.gaussian_filter, sigma=50) to remove any potential non-uniform background. We then enhanced the edges of the high intensity areas by sharpening the image as described in http://www.scipy-lectures.org/advanced/image_processing/. In brief, we consecutively low-pass filtered the image with Gaussian filter at two levels (sigma = 2 and 1). The differences in two images, which represent the edges of high intensity areas, were multiplied by 100 and added back to the first low-pass filtered image, resulting in a sharpened image. Finally, to avoid potential bleaching that may affect the overall intensity of the whole image, we normalized the intensity of each image by shifting the mean intensity to zero and divided by the standard deviation of the intensity.

We then calculated the cross-correlations between the enhanced reference image and each frame to obtain the displacement between the location of max coefficient and the center of the image. The shift that countered the displacement was then applied to the original, unenhanced image to complete the motion correction.

##### b) Identification of regions of interest from mouse in-vivo experiments

To identify the regions of interest (ROIs) that represent neurons, we first generated time-collapsed images by subtracting the average intensity value of each pixel over all videos from its maximum intensity. We then applied ACSAT (Shen et al., 2018) to generate ROIs with the following parameters: iteration=2, minimum size=50 pixels, and maximum size=300 pixels. In brief, ACSAT is a threshold-based ROI segmentation algorithm that adaptively adjusts the threshold at both global and local levels to capture ROIs with various intensities. Due to the shifting process during motion correction, the time-collapsed image often contains high intensity strips at the edge, which cause false-positive ROIs in ACSAT. Therefore, we excluded any ROIs within 10 pixels of the edge. Also, ROIs that were identified which were exceedingly large or small in size (less than 50 pixels or greater than 500 pixels) were excluded. Centroids were then identified for each ROI using the MATLAB command “regionprops” with the “centroid” argument.

##### c) Trace interpolation for mouse in-vivo experiments

While SomaGCaMP6f2 sessions were recorded at a constant rate of 20Hz, the sampling frequency for GCaMP6f sessions was controlled by a MATLAB script which introduced slight variability within the sampling rate (21.31 +/- 0.02 Hz (+/- s.d)). Therefore, traces for GCaMP6 were interpolated between the first and last time point in each 4-video sequence given by the time stamps of the corresponding Tiff files. Interpolation was performed with a constant sampling interval of 50ms (20 Hz) using linear interpolation (“interp1” in MATLAB).

##### d) Computation of df/f_0_ and linear detrending for mouse in-vivo experiments

After interpolating the traces from GCaMP6f sessions, df/f_0_ values were computed for each trace by subtracting its mean and dividing by its initial fluorescence. Each trace was then subject to a linear detrending using the MATLAB command “detrend”. Following this step, traces were each manually inspected to ensure that they had a dynamic nature and represented actual neurons. Traces that didn’t meet these qualifications were excluded from further analysis (n=12 SomaGCaMP6f2, and n=15 GCaMP6f cells).

##### e) Identification of homologous subregions from GCaMP6f session for mouse in-vivo experiments

To equalize the number of neurons recorded from each session and to keep the range of distances between cells consistent from different imaging sessions, only a portion of the full field was analyzed from each recording session. To do so, we highlighted subregions from each GCaMP6f session for further analysis. First, we characterized the visible brain region in each GCaMP6f session by computing a bounding box around the area of cell labeling, and computed the total number of neurons in each bounding box. These computations were performed as follows:

First, an ROI mask was constructed for each session. Each mask was then morphologically closed using the MATLAB function imclose(*,strel), with “strel” a structuring element, in this case set to the shape of a disk with a radius of 30 pixels (strel(‘disk’,30)). Second, this image was morphologically eroded using the MATLAB command “imerode”, again using a “disk”-type structuring element but in this case with a radius of 10 pixels. Finally, the image was morphologically dilated using the MATLAB command “imdilate”, and a structuring element of a disk with radius 20 pixels. This produced an image with an opaque region encompassing the region of the image most densely laden with ROIs. Following these procedures, we computed a bounding box around this region using the command “regionprops” with a second argument of “boundingbox”. Finally, the number of ROIs with centroids in this bounding box was computed for each session. Limits of the bounding box used for calculating relative positions of the centroids were computed by rounding the coordinates of the x and y starting points of the bounding box, and taking those points between these values through the values (extent of x = round(x+width-1), extent of y = round(y+height-1)), where height and width are the properties of the bounding box returned by MATLAB. Centroids were rounded to their nearest whole pixel values for this analysis.

To compute the factors necessary to identify a bounding box across all other sessions, we computed summary statistics of these bounding boxes for each GCaMP6f session. To identify the height of our bounding box, we divided the height of each bounding box by the bounding box’s area, averaged these quantities, and then multiplied them by the average area across all bounding boxes. An analogous procedure was performed to find a suitable bounding box width. Lastly, the number of ROIs identified in each bounding box were averaged to find a target number of neurons. In summary, our target region had a height of approximately 396 µm, a width of approximately 804 µm, yielding an area of 3.1856e+05 µm^2^, with approximately 177 neurons in this region. Our SomaGCaMP6f2 data had an average bounding box height of approximately 373 µm, a width of approximately 715 µm, and an average area of 2.64e05 µm^2^. To locate an area that fulfilled these requirements, the height and width estimated were first both rounded to whole numbers. Then, first by vertical pixels and then by horizontal pixels, areas constituting the required widths and heights were searched and the number of neurons with (rounded) centroids within these areas were counted. After all rectangles with these characteristics were searched, the region identified that had a number of neurons closest to the average number of neurons in bounding boxes from all other sessions (∼177) was used as the region for analysis. If multiple regions had the same number of ROIs or were equally close in number, the first region that was identified was used. For the remainder of these analyses (peak characteristic comparison and pairwise-correlation analysis), only the identified ROIs within this region were used.

##### f) Event identification for mouse in-vivo experiments

Spectral frequency analysis has been shown to be a reliable tool for estimating calcium fluorescence events as it is less influenced by drifts in baseline activity (Deneux et al., 2016; Patel et al., 2015; Ruffinatti et al., 2013). Within our data we noticed that the onsets of Ca events could be detected using Fourier analysis where event onset coincided with increasing low frequency power (power_event_). To take advantage of this observation, we first calculated the spectrogram from traces (Matlab chronux, mtspecgramc with tapers=[2 3] and window=[1 0.05]), and averaged the power below 2 Hz. To detect any significant increase in power, we calculated the change in the power at each time point (power_diff_), and identified the outliers (3 median absolute deviations away from the median power) in power_diff_ (Matlab function isoutlier). For outliers that occurred at consecutive time points, we only kept the first outliner, which represented the start of the change. We further selected the outliers with positive power_diff_ as they were indicators for the increase in the power. After identifying the time points of the significant increase, we then determined the end of power_event_ by identifying the first time point where the power decreased.

To obtain the peaks and start points of Ca events, we first extended the end point of power_event_ to the second time point with decreased Ca signal. After extension, the peak was defined as the time point within power_event_ where the maximum Ca signal occurred, and the start point was defined as the time point with minimum Ca signal between the peak and the start of power_event_. To ensure the quality of Ca events, we excluded any Ca event with amplitude (the signal difference between the peak and the onset) less than 4 standard deviations of the trace in the 20 second time window prior to Ca event onset. At the end of this process, some Ca events were found to overlap. To address this issue, the final set of Ca events was set to be the union of all of the identified Ca events, and the peak amplitude of each new event was defined as the maximum of the event minus the minimum of the event.

##### g) Computation of peak characteristics for mouse in-vivo experiments

Once peaks were identified, we then determined their waveforms. Waveforms were defined as 10 seconds flanking (5 seconds before and 5 seconds following) an event peak. Once identified, we subtracted the minimum value off the waveform. Then, event rate, rise time and decay times were computed as follows. To compute the event rate for a particular session, the number of waveforms identified over the course of the session were totaled for each region of interest, and this number was then divided by the total length of the session. Next, rise times were computed using the mean post-minimum subtracted peak waveform taken across all waveforms for a given ROI. These waveforms are aligned naturally because each is centered around its peak. To obtain the rise and decay time for each ROI, we first calculated a threshold as following: all events were averaged together, centered around their peak maxima, and the following equation was used to determine a threshold value:

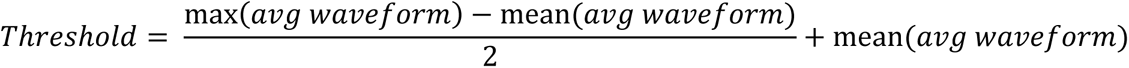

For rise time, the number of data points between the maximum of each identified event and the first point prior to the event where the trace fell to less than or equal to a significance threshold were computed. Falling times were computed by determining the number of data points between the maximum of an event and the first point following this maximum whose value dropped to a value less than or equal to the significance threshold. Any trace that lacked either an identified rise time or decay time, or both, was excluded from statistical analyses, and were also excluded from the computation of pairwise correlations. Event rates, fall times, and rise times computed ROI-wise from SomaGCaMP6f2 mice were compared with the respective values from ROIs in GCaMP6f mice via a Wilcoxon rank-sum test.

##### h) Pairwise-correlation analysis for mouse in-vivo experiments

Traces for each region of interest were truncated into 50 time point (2.5 second) segments in order to reduce the risk of non-stationarity of the df/f_0_ time traces, and correlation coefficients were computed pairwise over the course of each session. Pairwise correlation coefficients were then averaged over all of the segments of each session for each pair of ROIs. For statistical analysis, the average pairwise correlation coefficient across all ROI pairs for each recording session was computed, and results from GCaMP6f and SomaGCaMP6f2 animals were compared using a Wilcoxon rank-sum test.

### Electrophysiology

#### Current and voltage clamp recordings of cultured neurons

Whole cell patch clamp recordings in culture (for **Figure 2** and **Figure S2**) were made using Axopatch 200B or Multiclamp 700B amplifiers, a Digidata 1440 digitizer, and a PC running pClamp (Molecular Devices). For in vitro current-clamp recordings, neurons were patched 14–18 DIV (10–14 days post-transfection) to allow for sodium channel maturation. Neurons were bathed in room temperature Tyrode containing 125 mM NaCl, 2 mM KCl, 3 mM CaCl_2_, 1 mM MgCl_2_, 10 mM HEPES, 30 mM glucose and the synaptic blockers 0.01 mM NBQX and 0.01 mM GABAzine. The Tyrode pH was adjusted to 7.3 with NaOH and the osmolarity was adjusted to 300 mOsm with sucrose. For in vitro voltage-clamp recordings, neurons were patched 19-21 DIV (17-20 days post-transfection) and were done under similar conditions as current-clamp recordings, except the Tyrode also contained 1 μM tetrodotoxin (TTX, Tocris Bioscience). For recordings, borosilicate glass pipettes (Warner Instruments) with an outer diameter of 1.2 mm and a wall thickness of 0.255 mm were pulled to a resistance of 5–10 MΩ with a P-97 Flaming/Brown micropipette puller (Sutter Instruments) and filled with a solution containing 155 mM K-gluconate, 8 mM NaCl, 0.1 mM CaCl_2_, 0.6 mM MgCl_2_, 10 mM HEPES, 4 mM Mg-ATP, and 0.4 mM Na-GTP. The pipette solution pH was adjusted to 7.3 with KOH and the osmolarity was adjusted to 298 mOsm with sucrose.

#### Electrophysiology and calcium imaging in acute brain slice for cross talk analysis and assessment of sensitivity for spike number

Individual living slices (**Figure 3** and **Figure S4**) were transferred to a recording chamber mounted on an upright microscope (Olympus BX51WI) and continuously superfused (2–3 ml/min) with ACSF at room temperature. Cells were visualized through a 40x NA0.8 water-immersion objective to identify GCaMP6f-positive cells. Whole-cell current-clamp recordings were obtained from GCaMP6f-positive pyramidal neurons in layer 2/3 of motor cortex, using an Axopatch 700B amplifier (Molecular Devices) and Digidata 1440 digitizer (Molecular Devices). For recordings, borosilicate glass pipettes (Warner Instruments) with an outer diameter of 1.2 mm and a wall thickness of 0.255 mm were pulled to a resistance of 3–5 MΩ with a P-97 Flaming/Brown micropipette puller (Sutter Instruments) and filled with a solution containing 155 mM K-gluconate, 8 mM NaCl, 0.1 mM CaCl2, 0.6 mM MgCl2, 10 mM HEPES, 4 mM Mg-ATP, and 0.4 mM Na-GTP. The pipette solution pH was adjusted to 7.3 with KOH and the osmolarity was adjusted to 298 mOsm with sucrose. GCaMP fluorescence was excited by a SPECTRA X light engine (Lumencor) with 470/24 nm excitation filter (Semrock). To perform fair comparison of GCaMP6f1 and SomaGCaMP6f1 for **Figure 3 and S3**, excitation light power was adjusted on a cell-to-cell basis, in the range of 0.5 to 20 mW/mm^2^, to achieve similar intensity of fluorescence baseline between the two constructs. Fluorescence was collected through the same objective through a 525/50 nm emission filter and imaged onto an sCMOS camera (Andor Zyla5.5 or Hamamatsu Orca-Flash4.0 V2) at 50Hz acquisition frequency. For assessing the sensitivity of the GCaMP6f variants to action potential number using whole-cell patch clamp (**Figure S4A**) we performed 500 pA current injections (50 Hz current injections, 5 ms, in trains of 5, 10, or 20 pulses). For assessing crosstalk we performed the imaging as described above while stimulating cells in the slice with 0.1 mM 4-aminopyridine, aimed at producing low spike rates (as seen in **Figure 3I**).

### Imaging

#### Imaging GCaMP targeting variants in culture

GCaMP6f trafficking variants that were found to localize predominantly in the soma of cultured neurons (**Figure 1**, **Figure 2** and **Supplemental Table 1**) were imaged with an LED (X-Cite XLED1, Excelitas Tecnologies) mounted on a microscope for wide-field illumination (Leica 3000B), through a Leica HCX APO L 40x objective (air, NA=0.6). Imaging was performed with a Hamamatsu Orca Flash 4.0 camera using a 480 nm LED and GFP-3035D filter cube (Semrock) for GFP fluorescence (power, 34.84 mW/mm^2^).

#### Calcium imaging in acute brain slices for screening of somatic GCaMP6f variants

For **Figure S1**, individual slices were transferred to a recording chamber mounted on an inverted epifluorescnce microscope (Nikon Eclipse Ti inverted microscope equipped with 10x NA 0.3 objective lens, a SPECTRA X light engine (Lumencor) with 475/28 nm exciter (Semrock), and a 5.5 Zyla camera (Andor), controlled by NIS-Elements AR software) and continuously superfused (2–3 ml/min) with ACSF at room temperature. Cells were visualized through a 10x objective to identify GCaMP6f-positive cells under excitation light power in the range from 0.5 to 4 mW/mm^2^ adjusted to achieve comparable levels of baseline fluorescence for all screened constructs. 4-aminopyridine at a final concentration of 1 mM was added to induce neuronal activity.

#### Imaging GCaMP and SomaGCaMP6f1 in zebrafish

For **Figure 4A-G**, individual zebrafish larvae at 4-5 dpf expressing either GCaMP6f or SomaGCaMP6f1 were exposed to the paralytic agent alpha-bungarotoxin (Sigma Aldrich) for 30-45 seconds, at a concentration of 1 mg/ml. Then, the paralyzed fish were embedded in 1.5% ultralow-melting agarose (Sigma Aldrich) prepared in E3 medium, and imaged using a custom built 2-photon microscope. A forward moving grating was used as a stimulus as GCaMP6f or SomaGCaMP6f1 expressing cells were imaged at 15 Hz: for GCaMP6f experiments, 20s on / 20s off stimulus periods were used; for SomaGCaMP6f1, 10s on / 10s off (the difference in frequencies between GCaMP6f and SomaGCaMP6f1 was inadvertent).

For **Figure 4H-M** and **Figure S5**, individual zebrafish larvae at 4-5 dpf expressing either GCaMP6f or SomaGCaMP6f1 were exposed to the paralytic agent pancronium bromide (Sigma Aldrich) for 30-45 seconds, at a concentration of 0.20 mg/ml. The fish were under visual inspection until they stopped swimming. Then, the paralyzed fish were embedded in 1.5% ultralow-melting agarose (Sigma Aldrich) prepared in E3 medium. The embedded larva were mounted in an imaging chamber flooded with E3 medium, in a Lightsheet Z.1 microscope (Zeiss). For imaging, the fish were illuminated with an excitation laser line at 488 nm with maximum power of 50mW, through 10x/0.2NA illumination optics, and imaged through a 20x/1.0NA water dipping detection objective. Since the baseline fluorescence of SomaGCaMP6f1 was approximately 4.7 fold lower compared to GCaMP6f, the percentage of light power for GCaMP6f imaging was 5% while the light power for SomaGCaMP6f1 imaging was 22.5-25%. The fish were imaged at 25 Hz, downsampled to 1Hz, for periods of 10-20 minutes, while incubated with 1 mM 4-aminopyridine to induce spiking.

#### In vivo mouse imaging

For **Figure 5** and **Figure S6**, animals were positioned underneath a microscope, and imaged while freely locomoting on a spherical treadmill. For each animal, full session recordings (5-12 min) were performed while monitoring GCaMP fluorescence using the specifications noted below. Image acquisition occurred via a custom microscope equipped with a scientific CMOS (sCMOS) camera (ORCA-Flash4.0 LT Digital CMOS camera C11440-42U; Hamamatsu, Boston, MA). GCaMP was excited using a 5W LED (LZ1-00B200, 460 nm; LedEngin, San Jose CA). The custom microscope included a Leica N Plan 10X 0.25 PH1 microscope objective lens, a dual band excitation filter (FF01-468/553-25), a dichroic mirror (FF493/574-Di01-25×36), and a dual band emission filter (FF01-512/630-25; Semrock, Rochester, NY). Image acquisition was performed using HC Image Live (HC Image Live; Hamamatsu; Boston, MA). The exact sampling intervals varied based on demands of the Windows 7 operating system but was approximately 20Hz. For each image frame, exposure time was fixed at 20ms. Image data were stored as multi-page tagged image file format (mpTIFF’s).

### Animal surgery, training and behavior

#### Mouse surgery and virus injection (Figure 5 and Figure S6)

All animal procedures were approved by the Boston University Institutional Animal Care and Use Committee. Breeding pairs were obtained from Jackson Laboratory (Maine). A total of 11 mice (PV-cre mice; B6;129P2-Pvalb^tm1(cre)Arbr^/J), 8–12 weeks old at the start of the experiments, were used in these experiments. Both male and female mice were used in this study. Animals first underwent viral injection surgery targeting the left striatum under stereotaxic conditions (AP: +0.5, ML:-1.8 mm, DV: −1.6). Mice were injected with 500 nL of either (AAV9-Syn-GCaMP6f.WPRE.SV40; n=7; titer: 6.6 e12 GC/ml) or 500-800 nL AAVDJ-CAG-SomaGCaMP6f2; n=3; titer: 2.4e12 GC/ml, or 500nL AAVDJ-Syn-SomaGCaMP6f2; n=1; titer:5.6e12 GC/ml. AAV9-Syn-GCaMP6f was obtained from the University of Pennsylvania Vector Core and AAV9-Soma-GCaMP6f2 was obtained from the University of North Carolina Vector Core. All injections were made via pulled glass pipettes (diameter: 1.2 mm) pulled to a sharp point and then broken at the tip to a final inner diameter of ∼20 μm. Virus was delivered via slow pressure ejection (10-15 psi, 15-20 ms pulses delivered at 0.5 Hz). The pipette was lowered over 3 min and allowed to remain in place for 3 min before infusion began. The rate of the infusion was 100 nL/min. At the conclusion of the infusion, the pipette remained in place for 10 min before slowly being withdrawn over 2-3 minutes. Upon complete recovery (7+ days after virus injection, mice underwent a second procedure for the implantation of a sterilized custom imaging cannula (OD: 0.317 cm, ID: 0.236 cm, height, 2 mm diameter), fitted with a circular coverslip (size 0; OD: 3mm) adhered using a UV-curable optical adhesive (Norland Products). To access the dorsal striatum, the cortical tissue overlying the striatum was carefully aspirated away to expose the corpus callosum. The white matter was then thinned until the underlying striatal tissue could be visualized through the surgical microscope. The window was then placed and centered above the striatum. During the same surgery, a custom aluminum head-plate was attached to the skull, anterior to the imaging cannula.

#### Mouse Training (Figure 5 and Figure S6)

Following surgery for virus infusion and window implantation (typically about 21-28 days), mice were handled for several days before being headfixed to the treadmill/imaging apparatus. Mice then were habituated to running on the spherical treadmill while headfixed, 3-4 days per week, over the next two weeks at the same time of day as subsequent recordings. Each animal received at least 6 habituation sessions prior to the first recording day. Habituation was performed in the dark with the imaging LED illuminated to the same intensity as it would be for recording sessions.

#### Movement data acquisition (Figure 5 and Figure S6)

The spherical treadmill was constructed similar to that previously described by Dombeck et al.<u>^6^</u>. Briefly, the treadmill consisted of a 3D printed plastic housing and a Styrofoam ball supported with air. Movement was monitored using two computer USB mouse sensors affixed to the plastic housing at the midline of the Styrofoam ball. Each mouse sensor was mounted 3-4mm away from the surface of the ball to prevent interference with ball movement. The LED sensors projected on the ball surface 78 degrees apart. The x- and y-surface displacement measured by each mouse was acquired using a separate computer running a Linux OS (minimal CentOS 6), and a simple multi-threaded python script that asynchronously read and accumulated mouse motion events, and sent packaged <dx,dy> data at 100Hz to the image acquisition computer via a RS232 serial link. Packaged motion data were received on the imaging computer using a Matlab script that stored the accumulated motion between frame triggers synchronized to each acquired frame.

### In utero electroporation (Figure 3 and Figure S1)

Embryonic day (E) 15.5 timed-pregnant female Swiss Webster mice (Taconic) were deeply anesthetized with 2% isoflurane. Uterine horns were exposed and periodically rinsed with warm sterile phosphate buffered saline (PBS). A plasmid encoding GCaMP6f or SomaGCaMP6f variants under control of CAG promoter at final concentration 1-2 μg/μl diluted with PBS was injected into the lateral ventricle of the right cerebral hemisphere. Five voltage pulses (40 V, 50 ms duration, 1 Hz) were delivered using 5 mm round plate electrodes (ECM™ 830 Electroporation Generator, Harvard Apparatus). Injected embryos were placed back into the dam, and allowed to mature to delivery. All experimental manipulations were performed in accordance with protocols approved by the Massachusetts Institute of Technology Committee on Animal Care and were in accordance with the National Institutes of Health *Guide for the Care and Use of Laboratory Animals*.

### Acute brain slice preparation

Acute brain sections for cross talk analysis, and spike number sensitivity assessment (**Figure 3** and **Figure S4** respectively) were prepared using in utero electroporated mice at P12 – P24, as described above. Mice were used without regard for sex. Mice were anaesthetized by isoflurane inhalation, euthanized, and cerebral hemispheres were removed, placed in ice cold choline-based cutting solution consisting of (in mM): 110 choline chloride, 25 NaHCO_3_, 2.5 KCl, 7 MgCl_2_, 0.5 CaCl_2_, 1.25 NaH_2_PO_4_, 25 glucose, 11.6 ascorbic acid, and 3.1 pyruvic acid (339-341 mOsm/kg; pH 7.75 adjusted with NaOH), blocked and transferred into a slicing chamber containing ice-cold choline-based cutting solution. Coronal slices (300 μm thick) were cut with a Compresstome VF-300 slicing machine, transferred to a holding chamber with artificial cerebrospinal fluid (ACSF) containing (in mM) 125 NaCl, 2.5 KCl, 25 NaHCO3, 2 CaCl2, 1 MgCl2, 1.25 NaH2PO4 and 11 glucose (300-310 mOsm/kg; pH 7.35 adjusted with NaOH), and recovered for 10 min at 34 °C, followed by another 50 min at room temperature. Slices were subsequently maintained at room temperature until use. Both cutting solution and ACSF were constantly bubbled with 95% O_2_/5% CO_2_.

### Histological analysis of GCMP6f and SomaGCaMP6f1 in the mouse brain

Deeply anesthetized mice were perfused transcardially with 4% paraformaldehyde in 0.1 M phosphate buffer (pH 7.3) and brains were postfixed for 4 h at 4°C. 50 μm sections were cut with a Leica VT1000s vibratome and imaged using an inverted Nikon Eclipse Ti microscope equipped with a spinning disk sCSUW1 confocal scanner unit (Yokogawa, Tokyo, Japan), 642 nm solid state laser, a 40x, NA 1.15 objective (Nikon), and a 4.2 PLUS Zyla camera (Andor), controlled by NIS-Elements AR software.

## Acknowledgements

We thank Eftychios Pnevmatikakis and Andrea Giovannucci from the Flatiron Institute at the Simons Foundation for useful discussions about computational methods for neuropil contamination reduction, and for their active development of NormCorre and CaImAn. K.M.T. was a New York Stem Cell Foundation - Robertson Investigator and McKnight Scholar and this work was supported by funding from R01-MH102441 (NIMH) and Pioneer Award DP1-AT009925 (NCCIH). C.A.S. was supported by NIH grants F32 MH111216 (NIMH) and K99 DA045103 (NIDA), and a NARSAD Young Investigator Award (Brain and Behavior Research Foundation). E.S.B. was supported by John Doerr, NIH 1R01DA045549, NIH 1R01MH114031, the HHMI-Simons Faculty Scholars Program, Human Frontier Science Program RGP0015/2016, U. S. Army Research Laboratory and the U. S. Army Research Office under contract/grant number W911NF1510548, US-Israel Binational Science Foundation Grant 2014509, NIH 2R01DA029639, NSF CBET 1344219, the MIT Media Lab, the Open Philanthropy Project, NIH 1R24MH106075, NIH R44EB021054, NIH 1R01GM104948, and NIH Director’s Pioneer Award 1DP1NS087724.

## Supplemental figures

**Figure S1.**
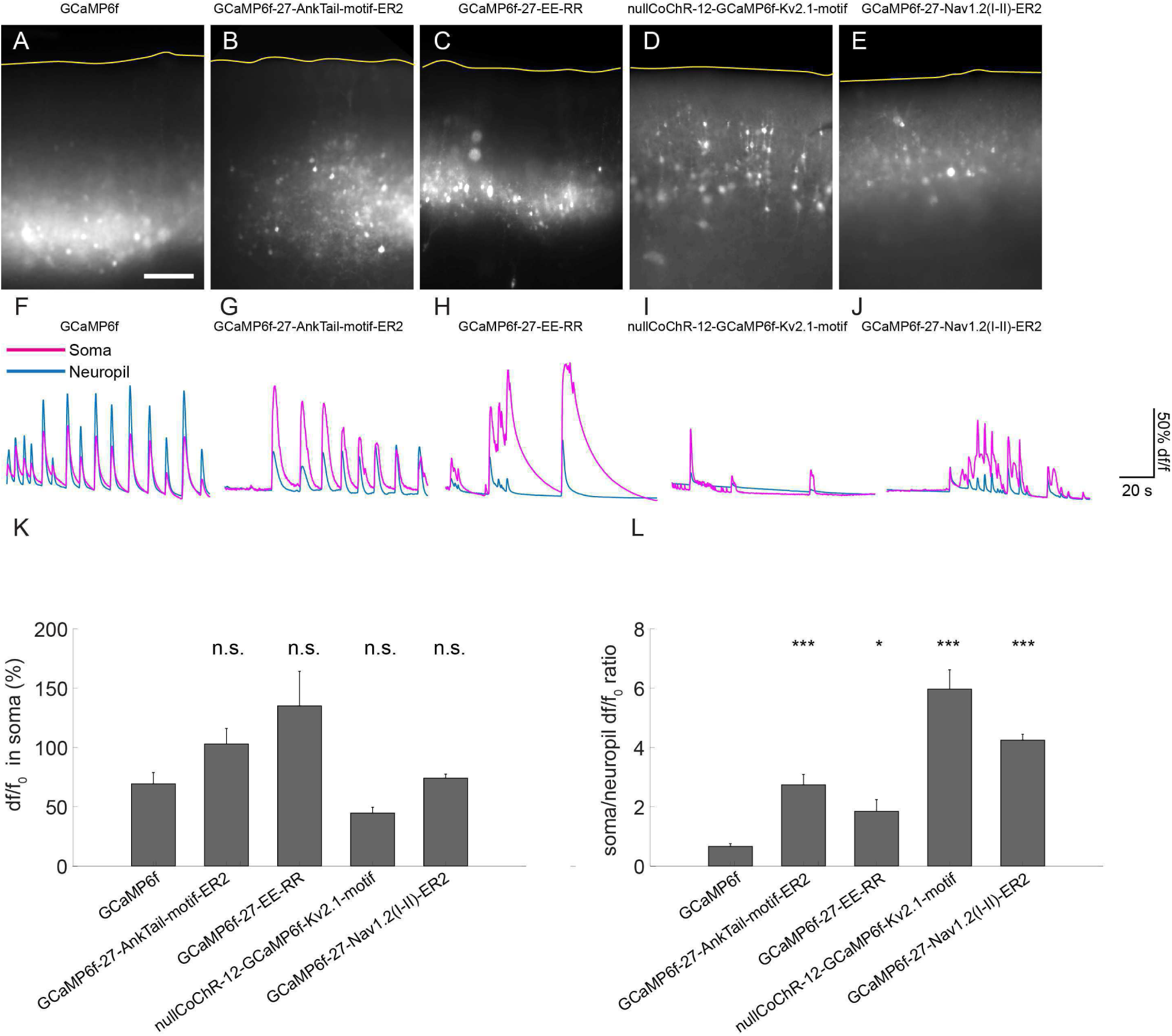
Slice imaging of soma-targeted GCaMP6f candidates during 4-Aminopyridine incubation. (A-E) Representative images of slices expressing GCaMP6f targeting variants. Scale bar: 200µm. Yellow line, edge of the brain. (A) GCaMP6f. (B) GCaMP6f-27-AnkTail-motif-ER2. (C) GCaMP6f-27-EE-RR. (D) nullCoChR-12-GCaMP6f-K_V_2.1-motif. (E) GCaMP6f-27-Na_V_1.2(I-II)-ER2. (F-J) Representative traces of the GCaMP signals from the soma (magenta) and the neuropil (blue). (F) GCaMP6f. (G) GCaMP6f-27-AnkTail-motif-ER2. (H) GCaMP6f-27-EE-RR. (I) nullCoChR-12-GCaMP6f-K_V_2.1-motif. (J) GCaMP6f-27-Na_V_1.2(I-II)-ER2. (K) A bar chart showing df/f_0_ in the somata of neurons expressing different GCaMP6f targeting variants (n = 20 cells from 2 slices from 2 mice for each variant). n.s., not significant, Kruskal-Wallis analysis of variance followed by post-hoc test via Steel’s test with GCaMP6f as control group; see **Supplemental Table 7** for full statistics for **Figure S1**. (L) A bar chart showing the ratio between df/f_0_ of the cell body and df/f_0_ of the neuropil for different GCaMP6f targeting variants (n = 20 cells from 2 slices from 2 mice for each variant). *P < 0.05, ***P < 0.001, Kruskal-Wallis analysis of variance followed by post-hoc test via Steel’s test with GCaMP6f as control group; see **Supplemental Table 7** for full statistics for **Figure S1**.

**Figure S2.**
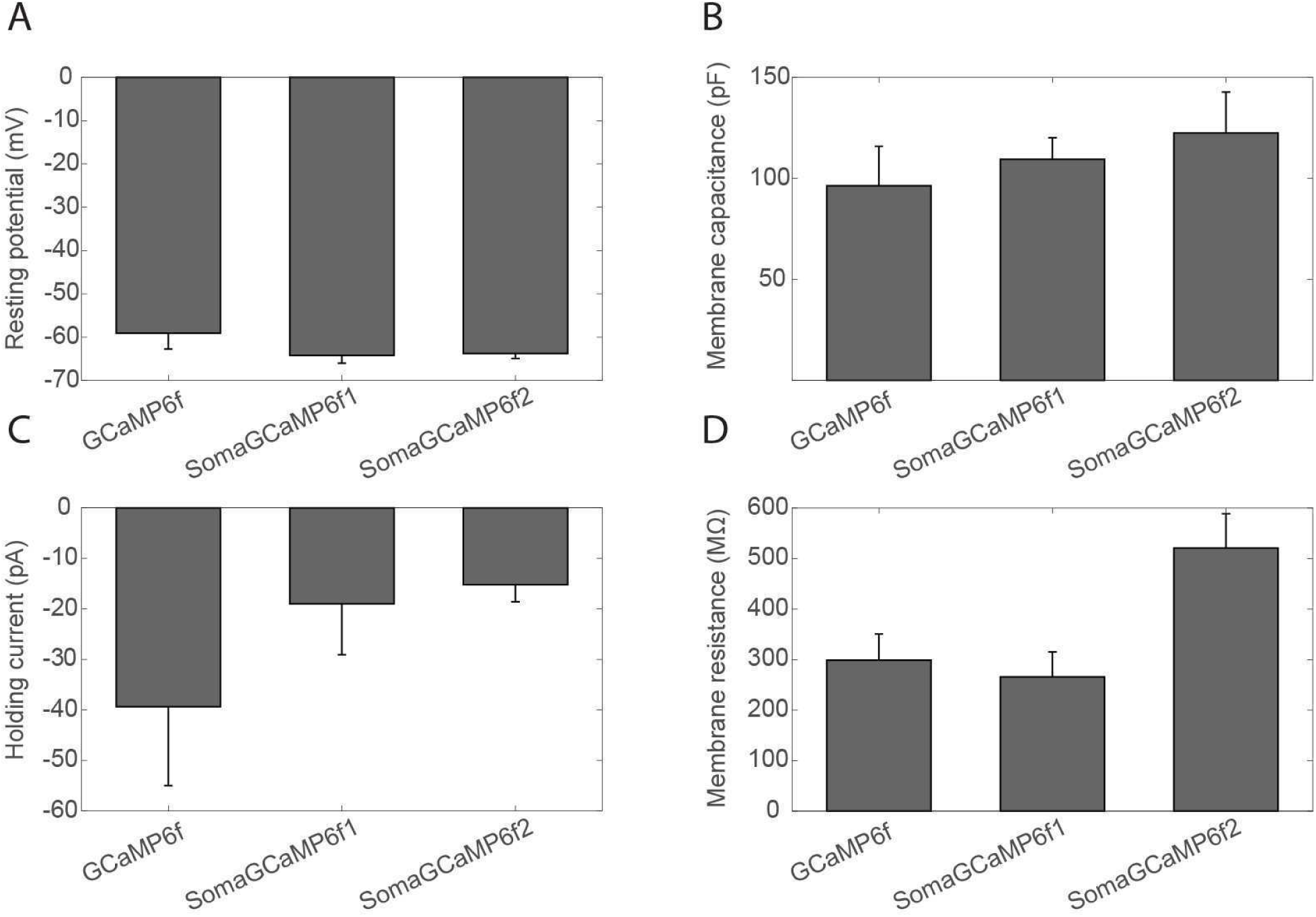
Properties of neurons expressing somatic vs. untargeted forms of GCaMP6f. Cultured hippocampal neurons expressing GCaMP6f, SomaGCaMP6f1 and SomaGCaMP6f2 were patched and membrane properties recorded. (**A**) Resting potential (n = 6 cells from 2 cultures for GCaMP6f; n = 7 cells from 2 cultures for SomaGCaMP6f1; n = 6 cells from 2 cultures for SomaGCaMP6f2.). Plotted is mean plus or minus standard error throughout the figure. Not significant, Kruskal-Wallis analysis of variance followed by post-hoc test via Steel’s test with GCaMP6f as control group. (**B**) Membrane capacitance (n = 5 cells from 2 cultures for GCaMP6f; n = 6 cells from 2 cultures for SomaGCaMP6f1; n = 6 cells from 2 cultures for SomaGCaMP6f2.). Not significant, Kruskal-Wallis analysis of variance followed by post-hoc test via Steel’s test with GCaMP6f as control group. (**C**) Holding current while held at −65 mV (n = 5 cells from 2 cultures for GCaMP6f; n = 6 cells from 2 cultures for SomaGCaMP6f1; n = 6 cells from 2 cultures for SomaGCaMP6f2.). Not significant, Kruskal-Wallis analysis of variance followed by post-hoc test via Steel’s test with GCaMP6f as control group. (**D**) Membrane resistance (n = 5 cells from 2 cultures for GCaMP6f; n = 6 cells from 2 cultures for SomaGCaMP6f1; n = 6 cells from 2 cultures for SomaGCaMP6f2.). Not significant, Kruskal-Wallis analysis of variance followed by post-hoc test via Steel’s test with GCaMP6f as control group. See **Supplemental Table 8** for full statistics for **Figure S2**.

**Figure S3.**
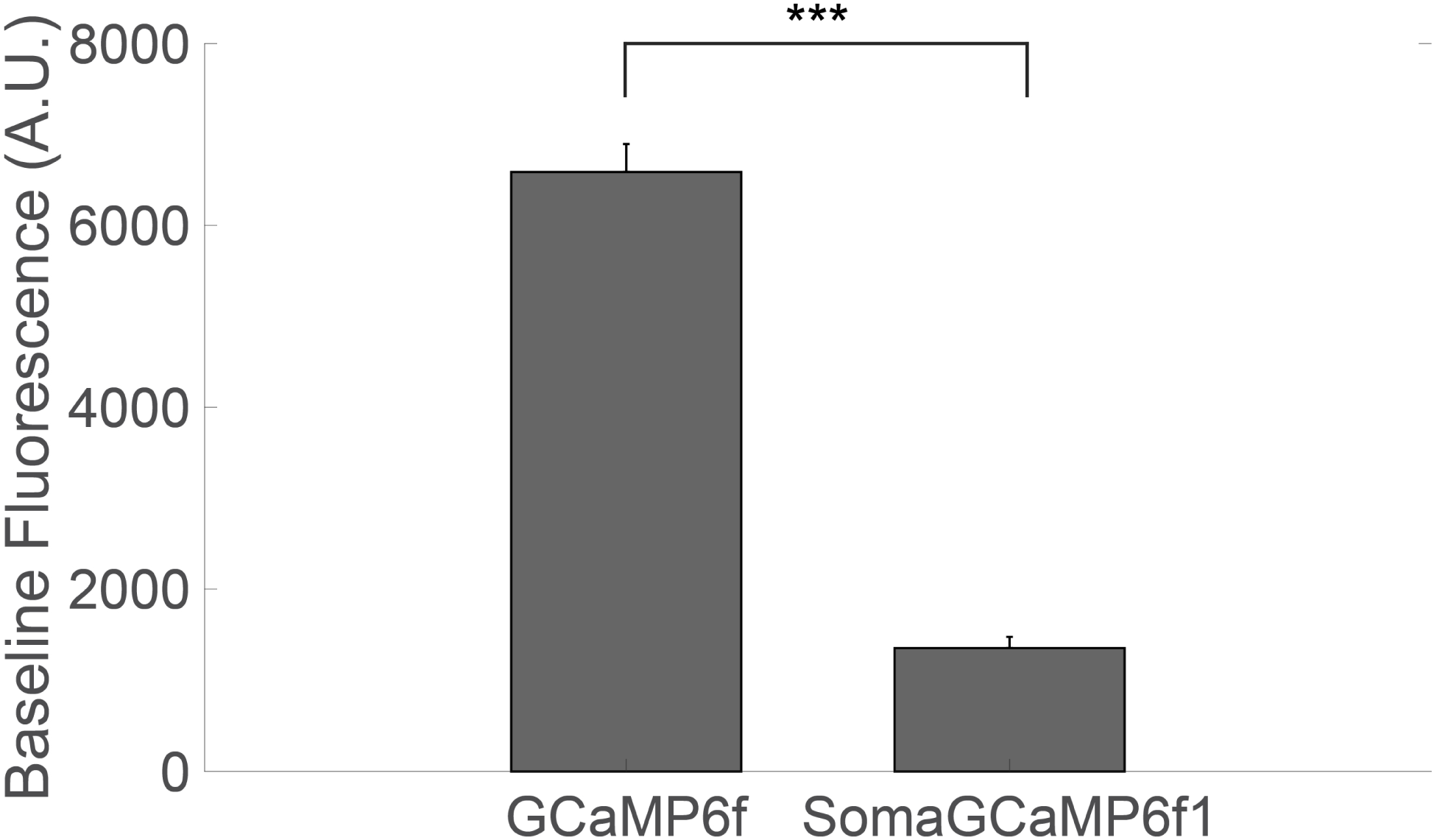
Baseline fluorescence brightness of GCaMP6f and SomaGCaMP6f1 in living brain slices. Bars show average baseline brightness values for cells expressing GCaMP6f or SomaGCaMP6f1 in slice (n = 42 neurons from 4 slices from 2 GCaMP6f mice; n = 43 neurons from 8 slices from 3 SomaGCaMP6f1 mice). Error bars indicate standard error of the mean. ***P < 0.001, Kolmogorov-Smirnov test of baseline fluorescence brightness between GCaMP6f and SomaGCaMP6f1; see **Supplemental Table 9** for full statistics for **Figure S3**.

**Figure S4.**
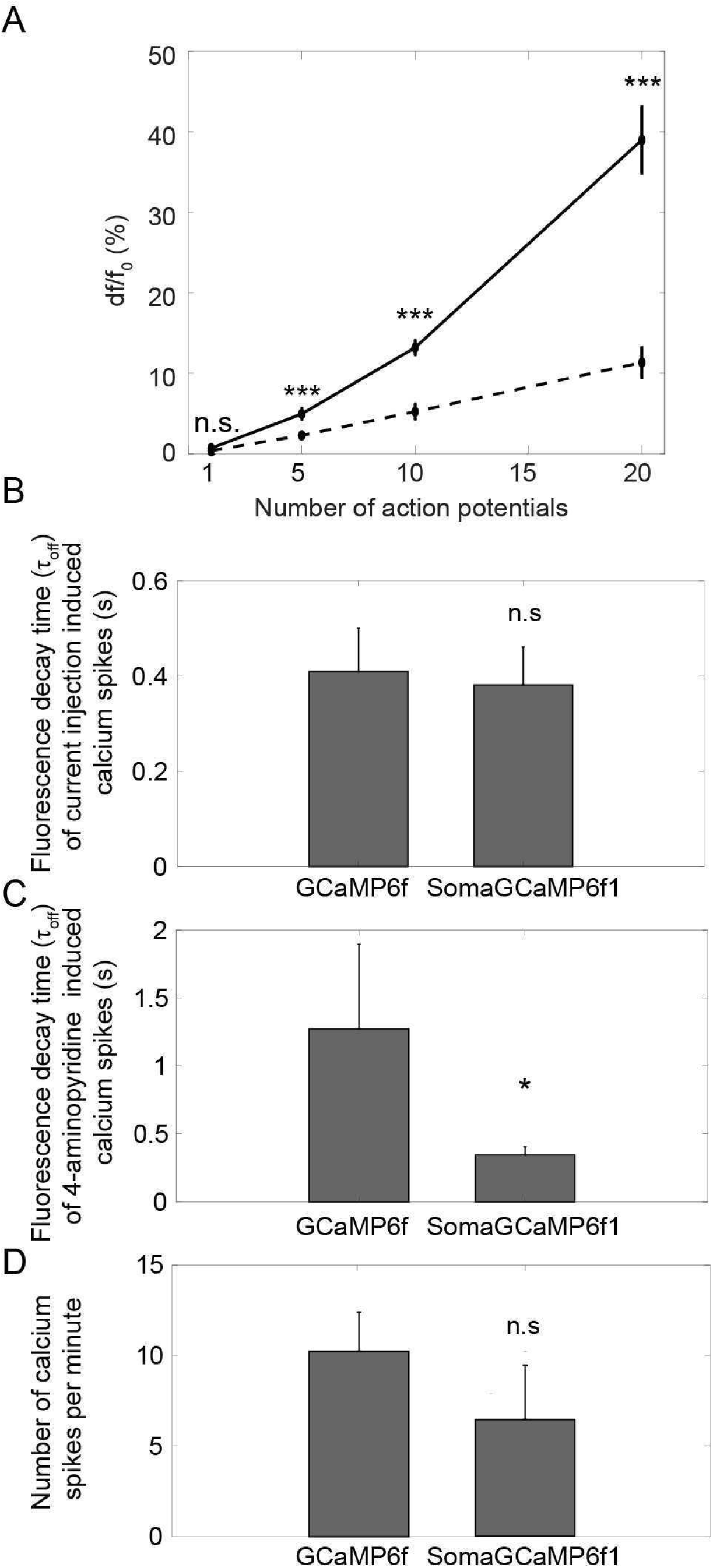
Sensitivity of multiple action potentials, temporal dynamics and event rate for GCaMP6f and SomaGCaMP6f1. **(A)** A graph showing the df/f_0_ of the calcium transient elicited after a train of 1, 5, 10 and 20 current pulses (500 pA, 5 ms duration, 50 Hz) for neurons expressing GCaMP6f (dotted line) or SomaGCaMP6f1 (unbroken line, n = 7 neurons from 5 slices from 2 mice for GCaMP6f; n = 5 neurons from 3 slices from 2 mice for SomaGCaMP6f1). n.s., not significant, ***P<0.001, Bonferroni-corrected Wilcoxon rank sum test of the df/f_0_ between GCaMP6f and SomaGCaMP6f1 expressing neurons; see **Supplemental Table 10** for full statistics for **Figure S4. (B)** Bar chart showing the mean τ_off_ of calcium spikes in slice, during electrophysiological inducement of single action potentials (n = 3 neurons from 3 slices from 3 mice for GCaMP6f; n = 3 neurons from 3 slices from 3 mice for SomaGCaMP6f1). n.s., not significant, Wilcoxon rank sum test between GCaMP6f and SomaGCaMP6f1; see **Supplemental Table 10** for full statistics for **Figure S4**. (**C**) Bar chart showing the mean τ_off_ of calcium spikes in slice, during 4-aminopyridine inducement of single action potentials (n = 5 neurons from 5 slices from 4 mice for GCaMP6f; n = 5 neurons from 4 slices from 3 mice for SomaGCaMP6f1). *P < 0.05, Wilcoxon rank sum test between GCaMP6f and SomaGCaMP6f1; see **Supplemental Table 10** for full statistics for **Figure S4**. **(D)** Bar chart showing the mean event rate of calcium spikes per minute in slice (n = 8 neurons from 8 slices from 4 mice for GCaMP6f; n = 6 neurons from 6 slices from 3 mice or SomaGCaMP6f1). n.s., not significant, Wilcoxon rank sum test between GCaMP6f and SomaGCaMP6f1; see **Supplemental Table 10** for full statistics for **Figure S4**.

**Figure S5.**
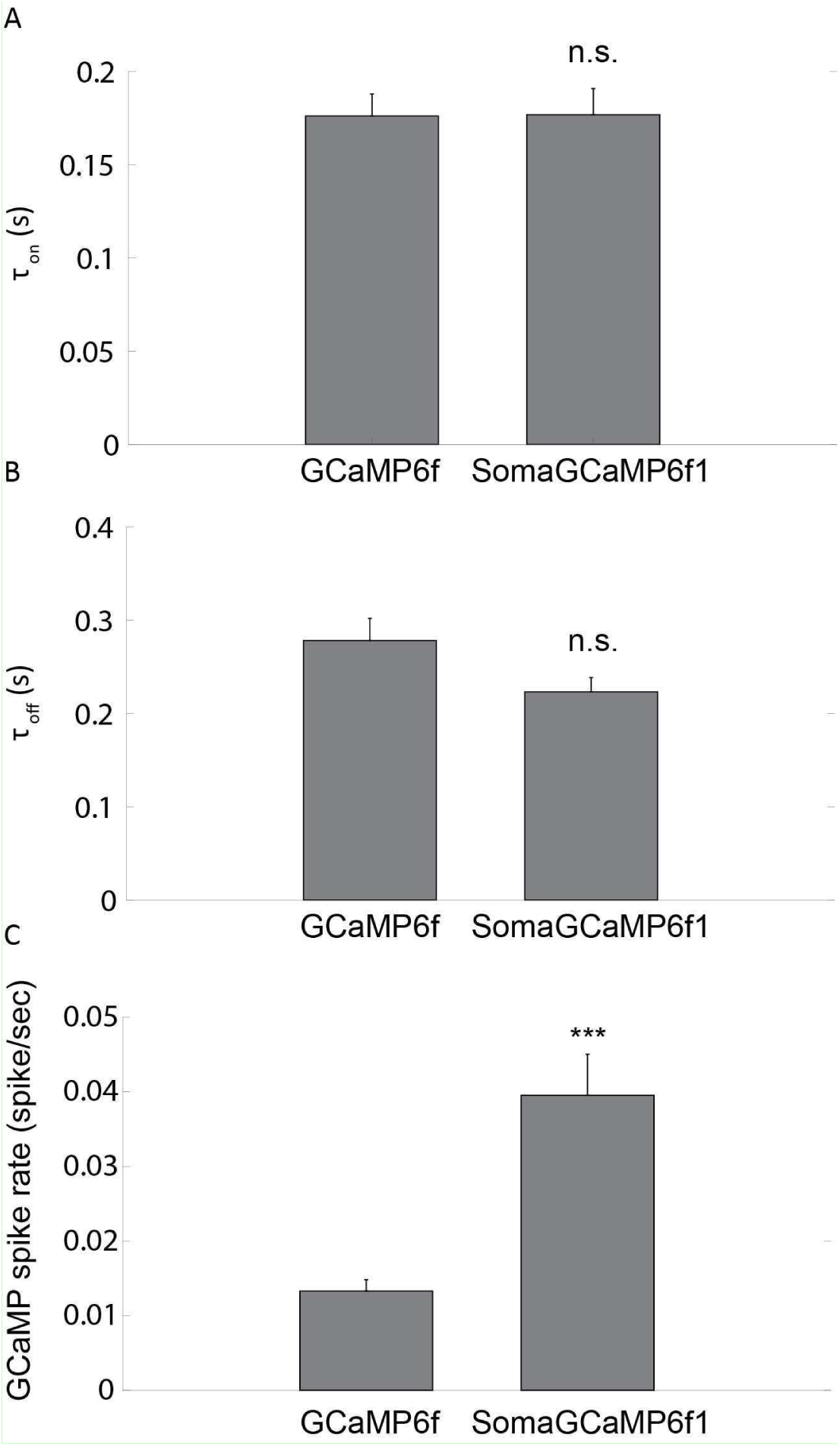
Temporal dynamics and calcium spike count for GCaMP6f and SomaGCaMP6f1 expressing neuron in zebrafish larva, driven by 4-AP. **(A)** Bar chart showing the mean τ_on_ of calcium spikes in zebrafish larva forebrain (n = 96 cells from 3 GCaMP6f fish; n = 146 cells from 4 SomaGCaMP6f1 fish). n.s., not significant, Wilcoxon rank sum test between GCaMP6f and SomaGCaMP6f1; see **Supplemental Table 11** for full statistics for **Figure S5**. **(B)** Bar chart showing the mean τ_off_ of calcium spikes in zebrafish larva forebrain (n = 96 cells from 3 GCaMP6f fish; n = 146 cells from 4 SomaGCaMP6f1 fish). n.s., not significant, Wilcoxon rank sum test between GCaMP6f and SomaGCaMP6f1; see **Supplemental Table 11** for full statistics for **Figure S5**. (**C**) A bar chart showing the mean GCaMP-spike rates for neurons in region of the larval zebrafish forebrain expressing either GCaMP6f or SomaGCaMP6f1 (n = 24 cells from 2 GCaMP6f fish; n = 24 cells from 2 SomaGCaMP6f1 fish). ***P<0.001, Wilcoxon rank sum test between GCaMP6f and SomaGCaMP6f1; see **Supplemental Table 11** for full statistics for **Figure S5**.

**Figure S6.**
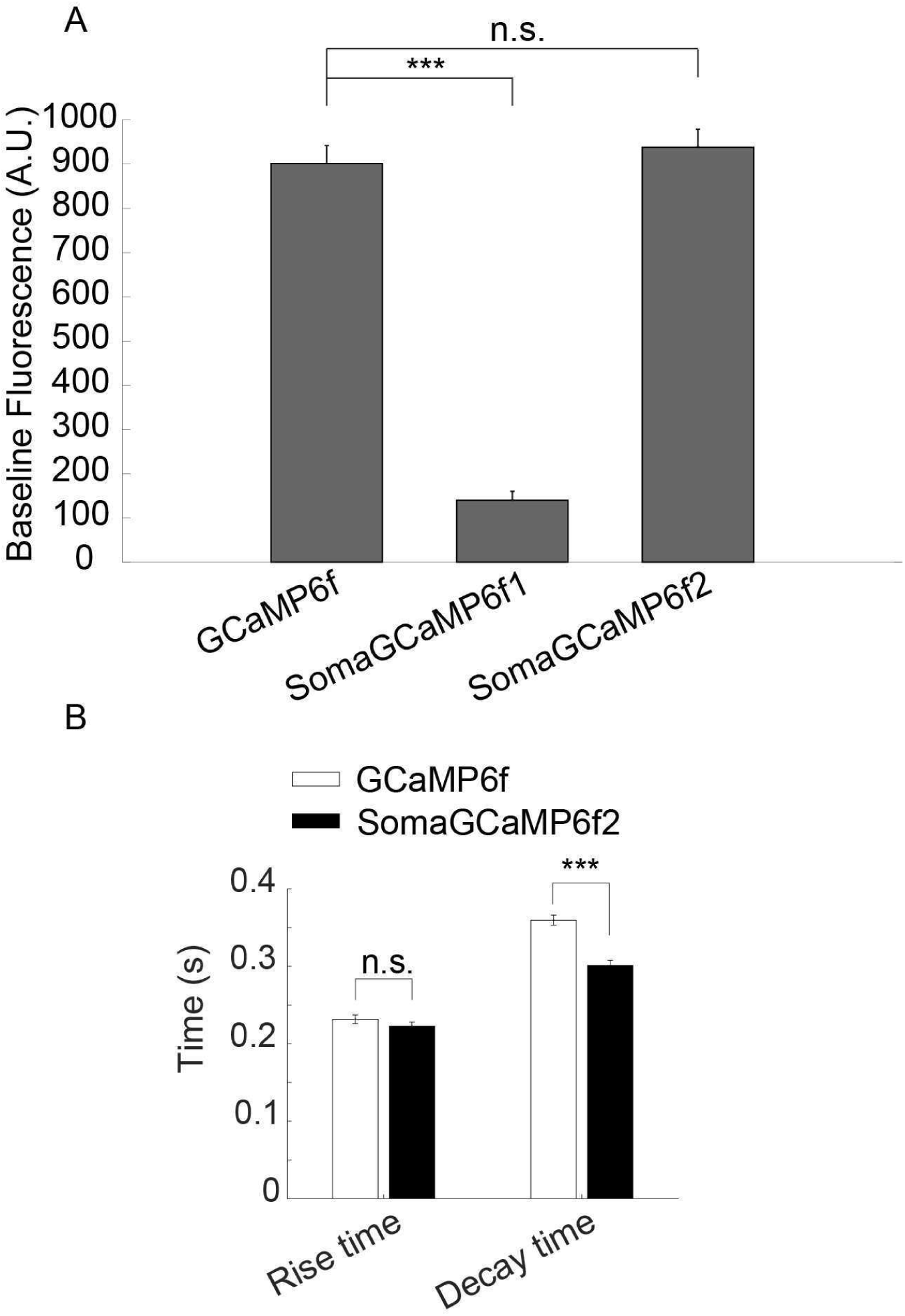
Baseline fluorescence brightness and kinetics of GCaMP6f and SomaGCaMP6f variants in mouse striatum in vivo. Bar chart showing the baseline fluorescence in vivo in the dorsal striatum for GCaMP6f, SomaGCaMP6f1 and SomaGCaMP6f2 (n = 75 neurons from 5 mice for GCaMP6f; n = 50 neurons from 2 mice for SomaGCaMP6f1; n = 80 neurons from 4 mice for SomaGCaMP6f2). ***P < 0.001, Kruskal-Wallis analysis of variance followed by post-hoc test via Steel’s test with GCaMP6f as control group; see **Supplemental Table 12** for full statistics for **Figure S6**. n.s., not significant, Kruskal-Wallis analysis of variance followed by post-hoc test via Steel’s test with GCaMP6f as control group; see **Supplemental Table 12** for full statistics for **Figure S6**. (**B**) Bar chart showing the average rise time (τ_on_) and the average decay time (τ_off_) for neurons expressing either SomaGCaMP6f2 or GCaMP6f (n = 594 neurons from 4 mice expressing SomaGCaMP6f2, n = 930 neurons from 7 GCaMP6f mice). n.s., not significant, Wilcoxon rank sum test between the rise times of SomaGCaMP6f2 and GCaMP6f expressing neurons; *** P <0.001, Wilcoxon rank sum test between the decay times of SomaGCaMP6f2 and GCaMP6f expressing neurons; see **Supplemental Table 12** for full statistics for **Figure S6**.

## Supplemental Information

**Supplemental Table 1:**

Proteins that were considered in this study as potential soma targeting fragments. For amino acid sequences corresponding to the acronyms used, see **Supplemental Table 13**.

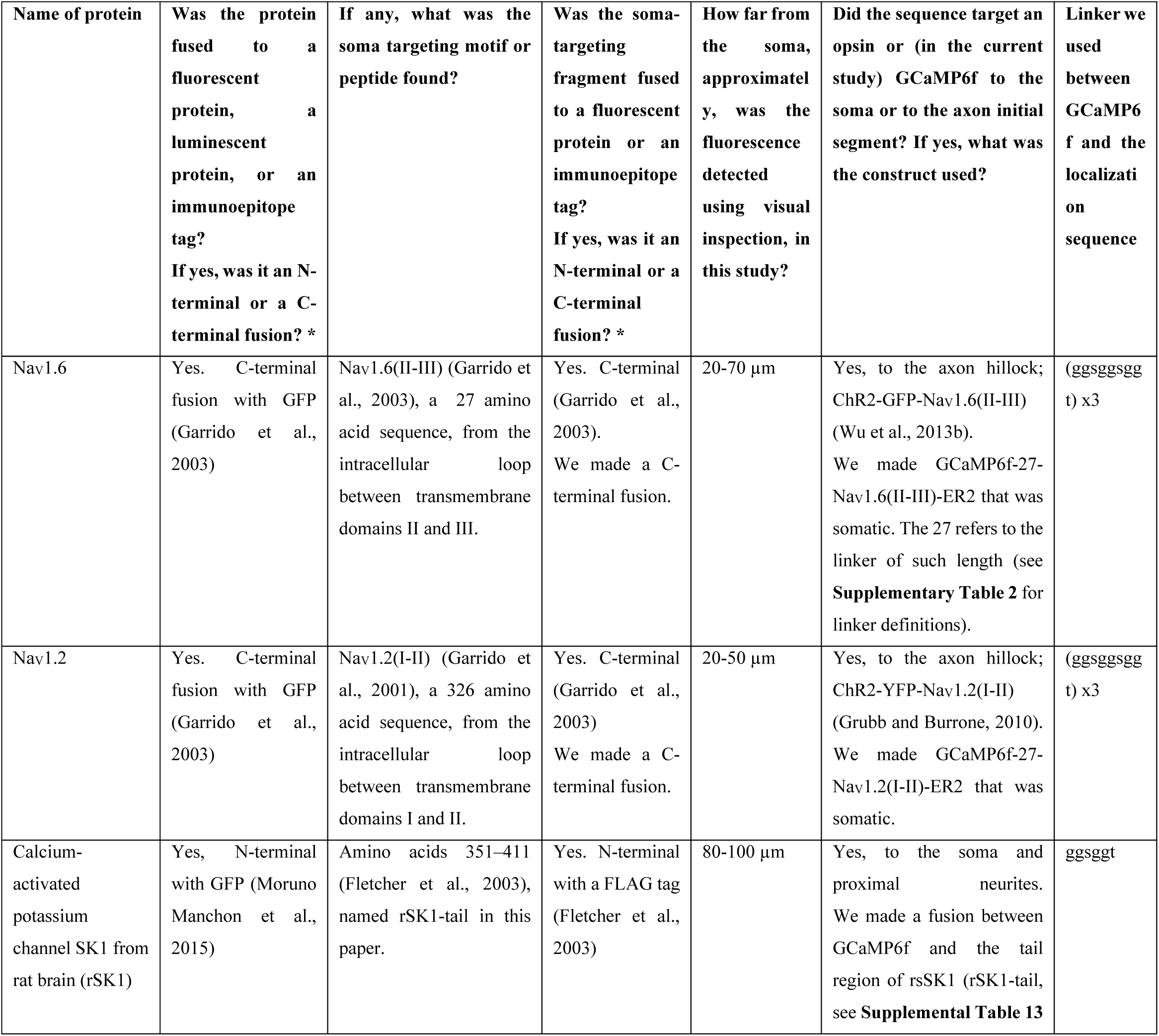
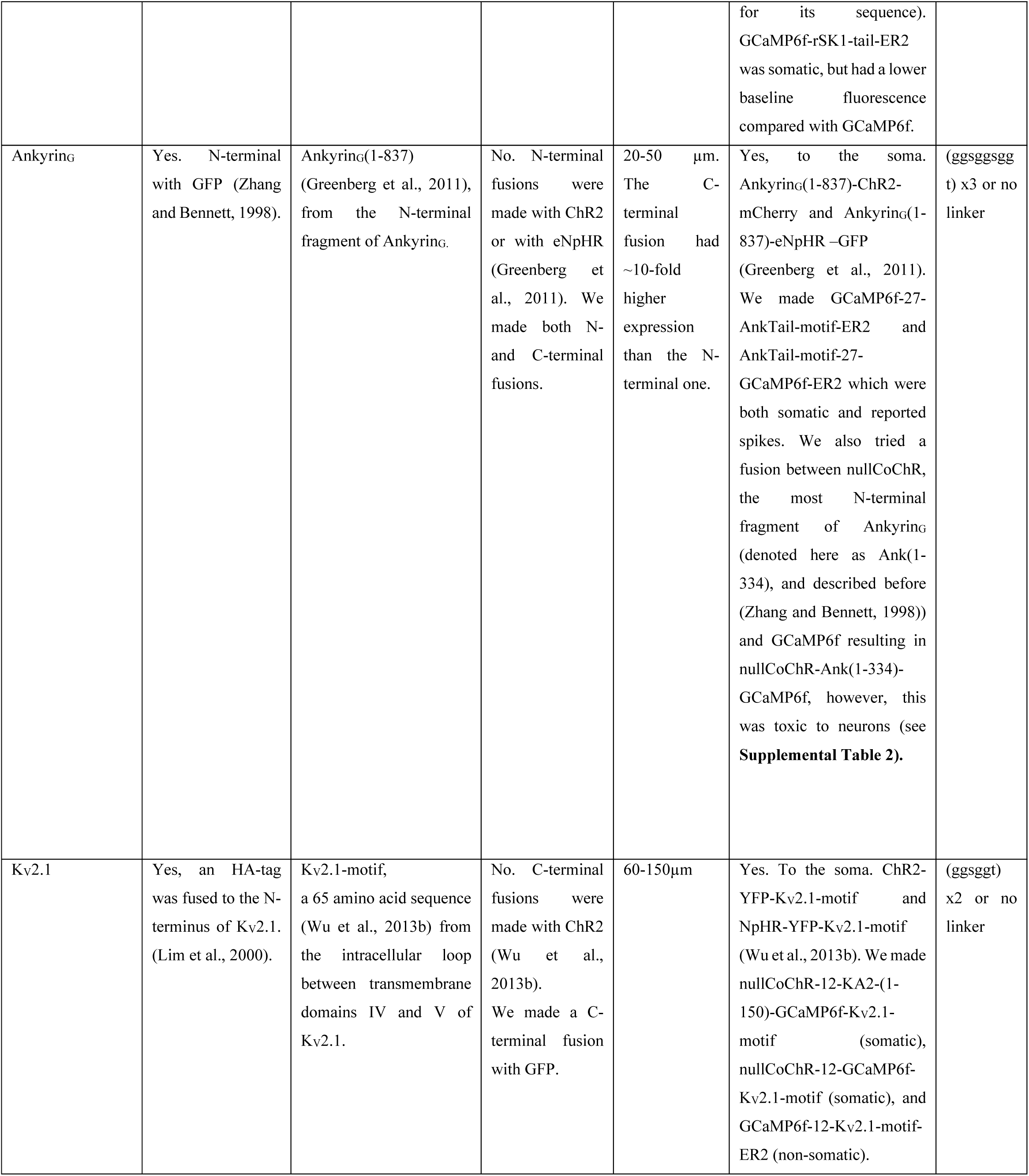
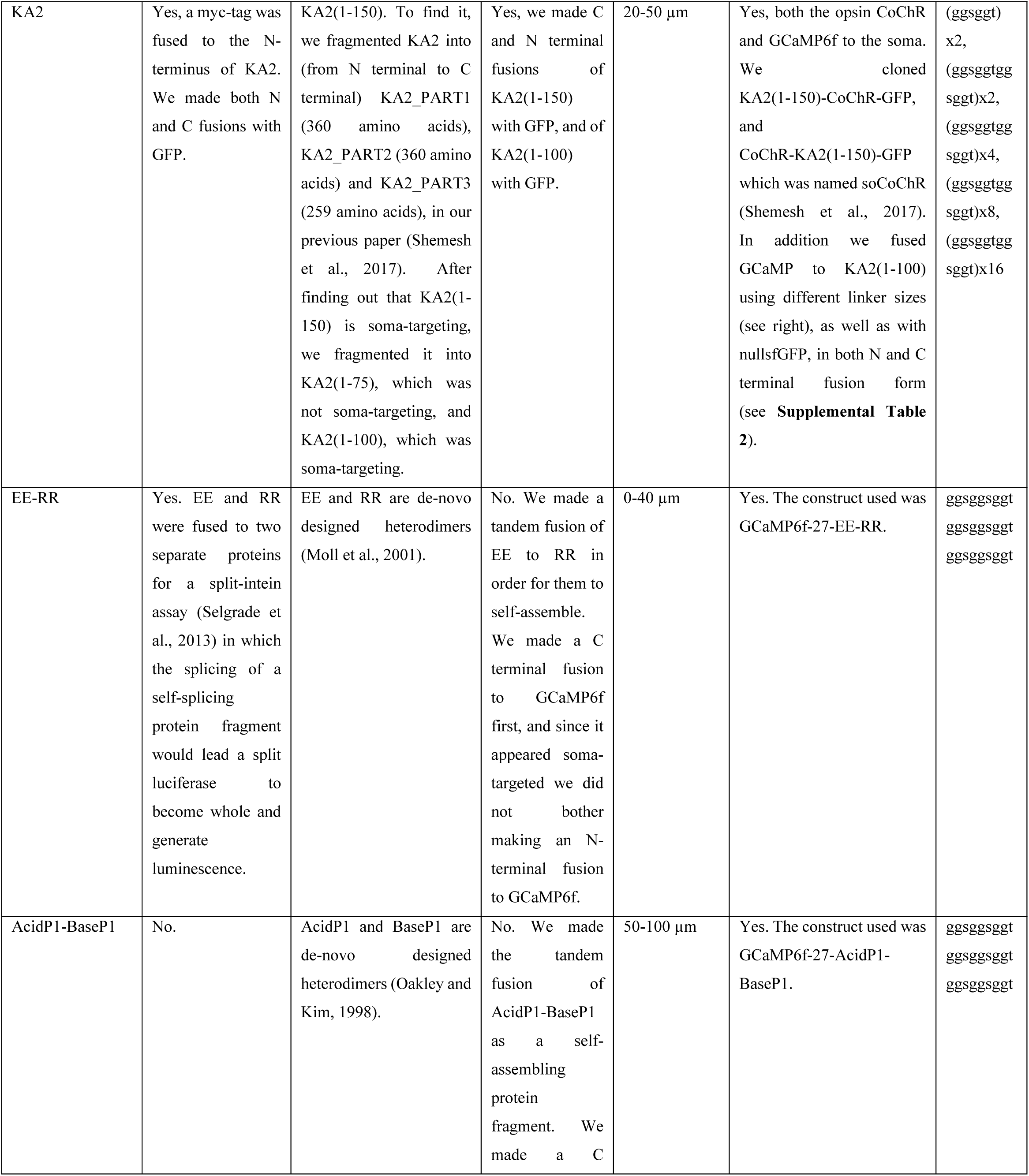
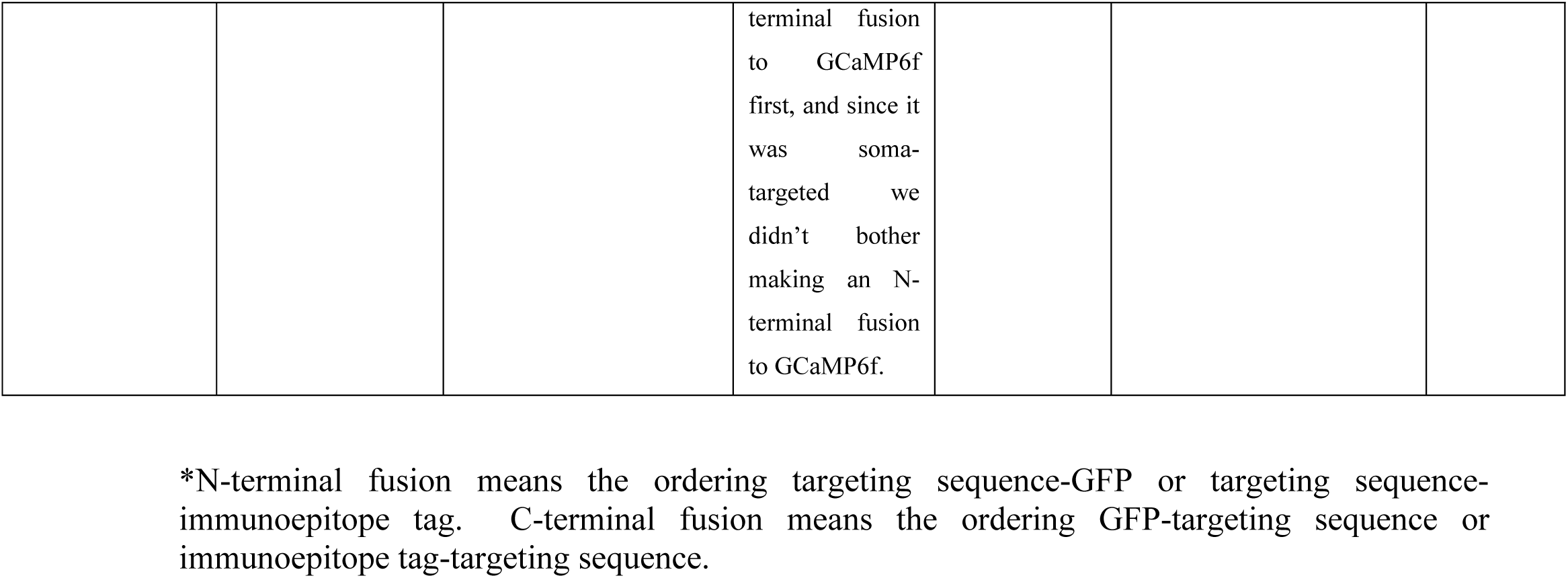

**Supplemental Table 2: GCaMP6f fusion proteins that were screened in cultured hippocampal neurons in this project**

In this table, the number inside the construct name is an abbreviation for the linker size:

12 = ggsggtggsggt

24 = ggsggtggsggtggsggtggsggt

27 = ggsggsggtggsggsggtggsggsggt

48 = ggsggtggsggtggsggtggsggtggsggtggsggtggsggtggsggt

96 = ggsggtggsggtggsggtggsggtggsggtggsggtggsggtggsggtggsggtggsggtggsggtggsggtggsggtggsggtggs

ggtggsggt

192 = ggsggtggsggtggsggtggsggtggsggtggsggtggsggtggsggtggsggtggsggtggsggtggsggtggsggtggsggtggs

ggtggsggtggsggtggsggtggsggtggsggtggsggtggsggtggsggtggsggtggsggtggsggtggsggtggsggtggsggt

ggsggtggsggtggsggt

KGC (Ma et al., 2001) and ER2 (Hofherr et al., 2005) are trafficking sequences from the potassium channel Kir2.1.

KA2(150)-Y76A is a mutant of KA2(1-150), in which the amino acid known for dimerization of KA2 (Kumar et al., 2011) was mutated to alanine (see **Supplemental Table 13** for sequences).

All the results in the following table are from cultured mouse hippocampal neurons (see STAR Methods).

For amino acid sequences corresponding to the acronyms used, see **Supplemental Table 13**.

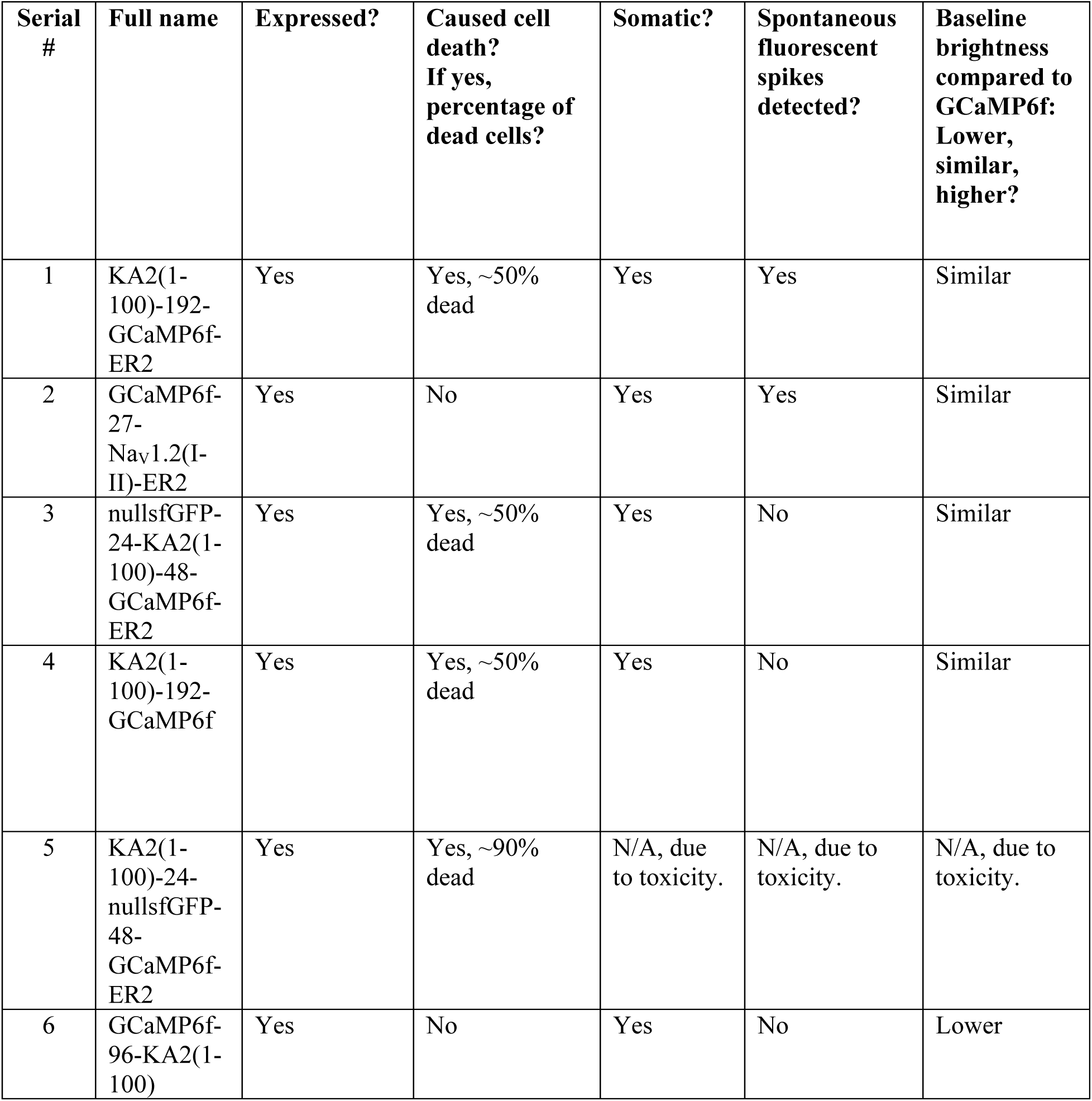
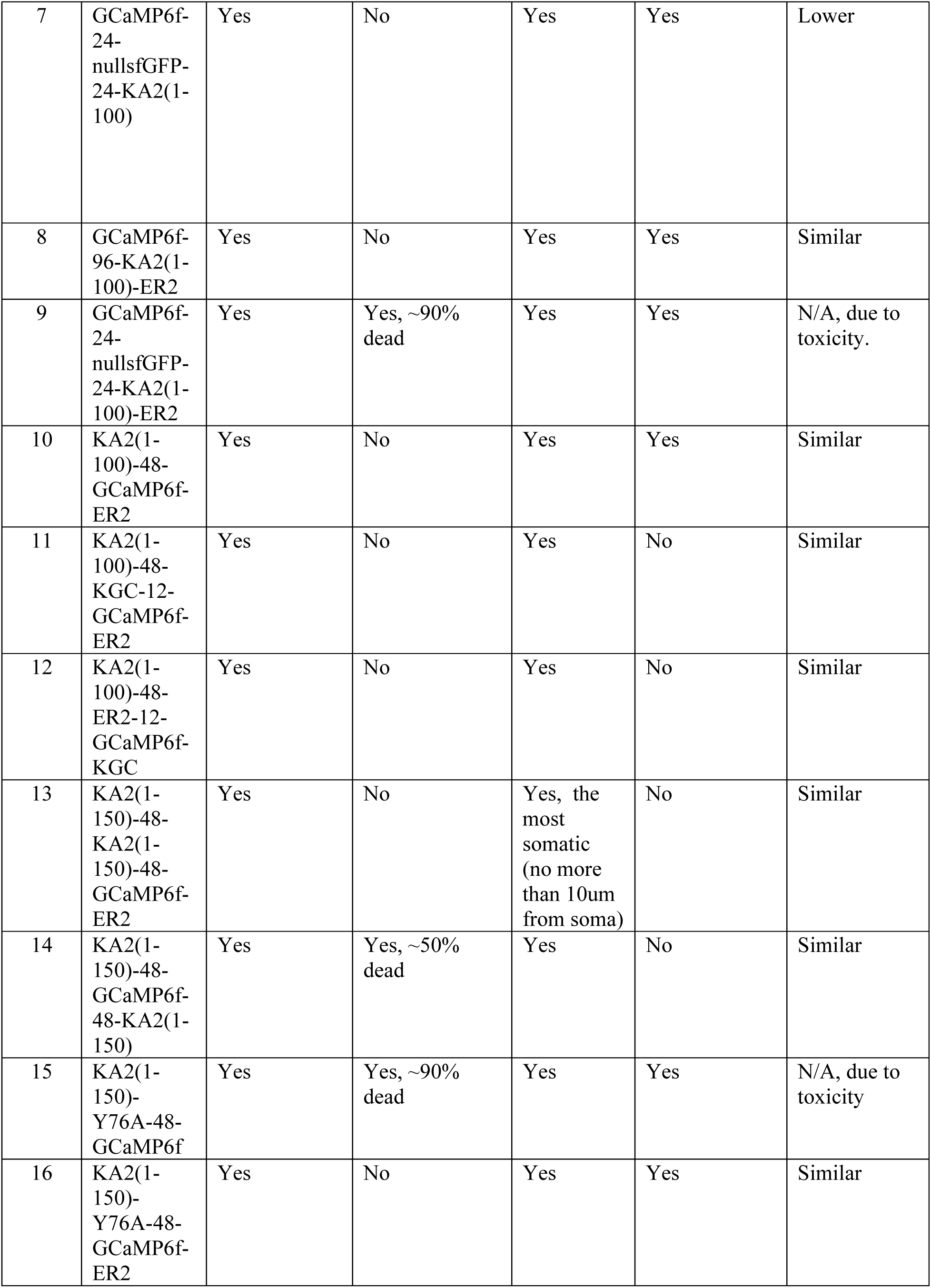
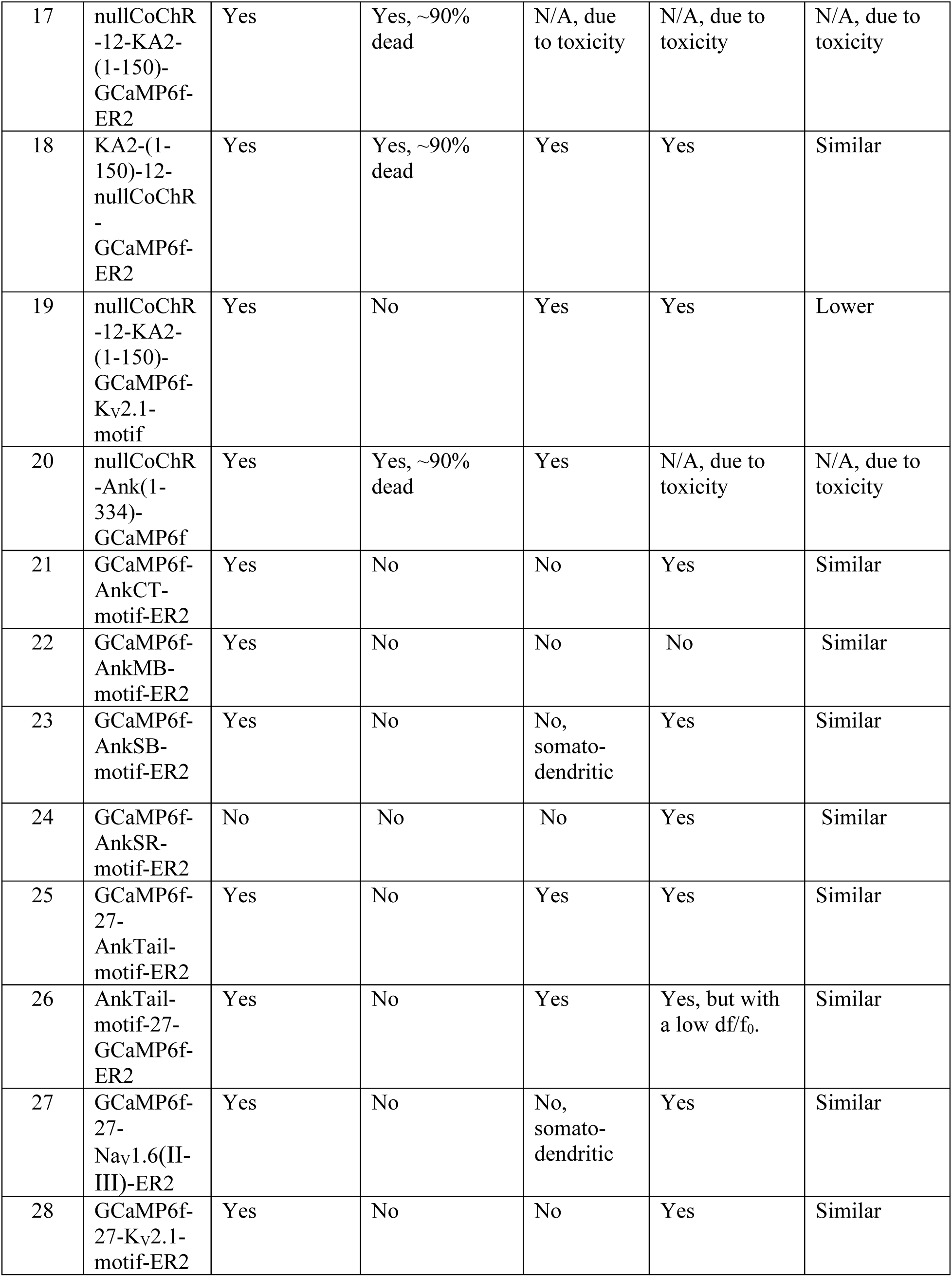
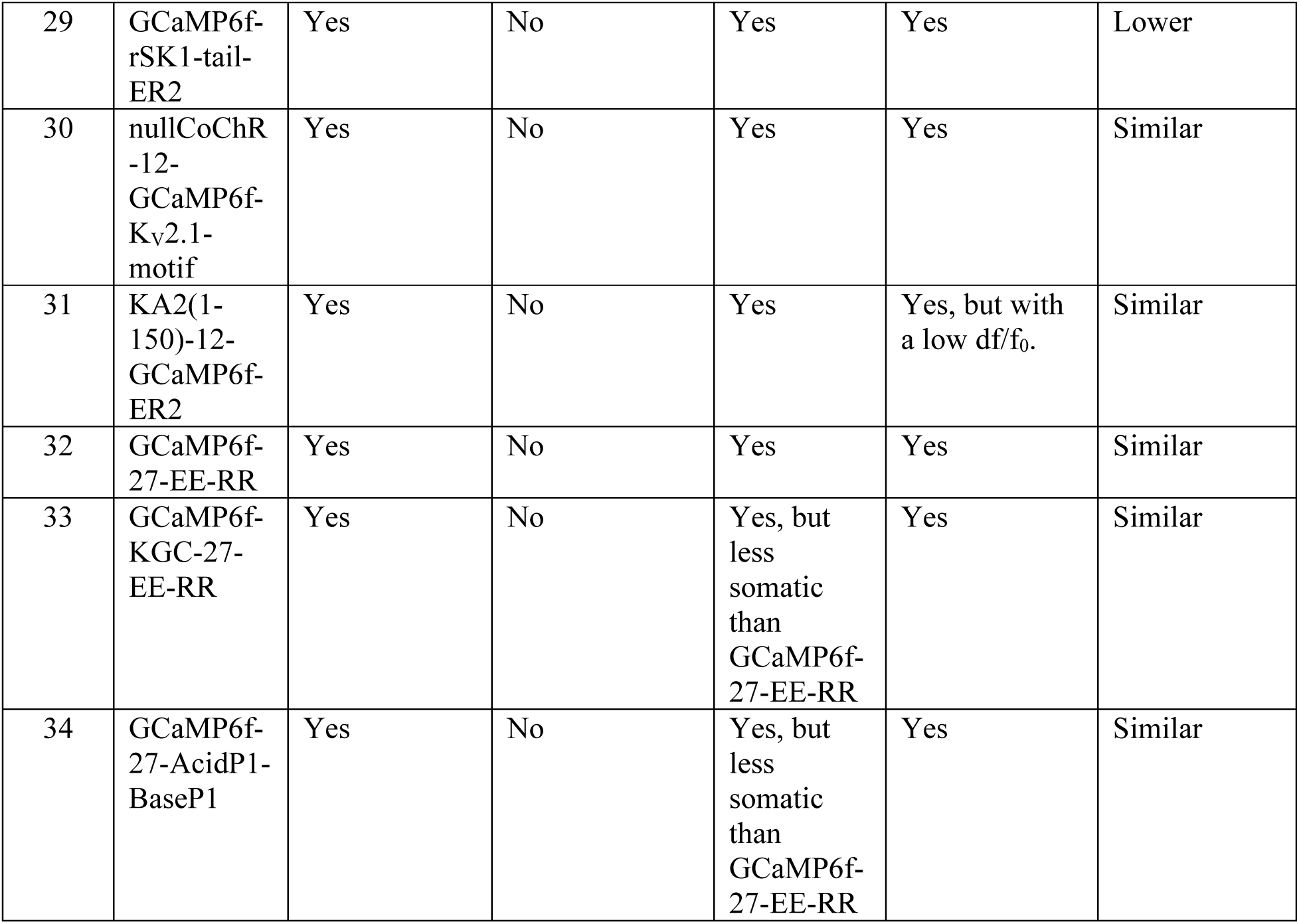

**Supplemental Table 3**: Statistical analysis for Figure 1.

Figure 1L, M, N - Brightness versus position along a neurite of GCaMP variants, normalized to GCaMP brightness at the soma

Kruskal-Wallis analysis of variance of neurite brightness followed by post-hoc test via Steel’s test with GCaMP6f as a control group. For GCaMP6f, n = 8 neurites from 8 cells from 3 cultures. For SomaGCaMP6f1, n = 5 neurites from 5 cells from 2 cultures. For SomaGCaMP6f2, n = 5 neurites from 5 cells from 3 cultures.

Wilcoxon / Kruskal-Wallis Tests (Rank Sums)

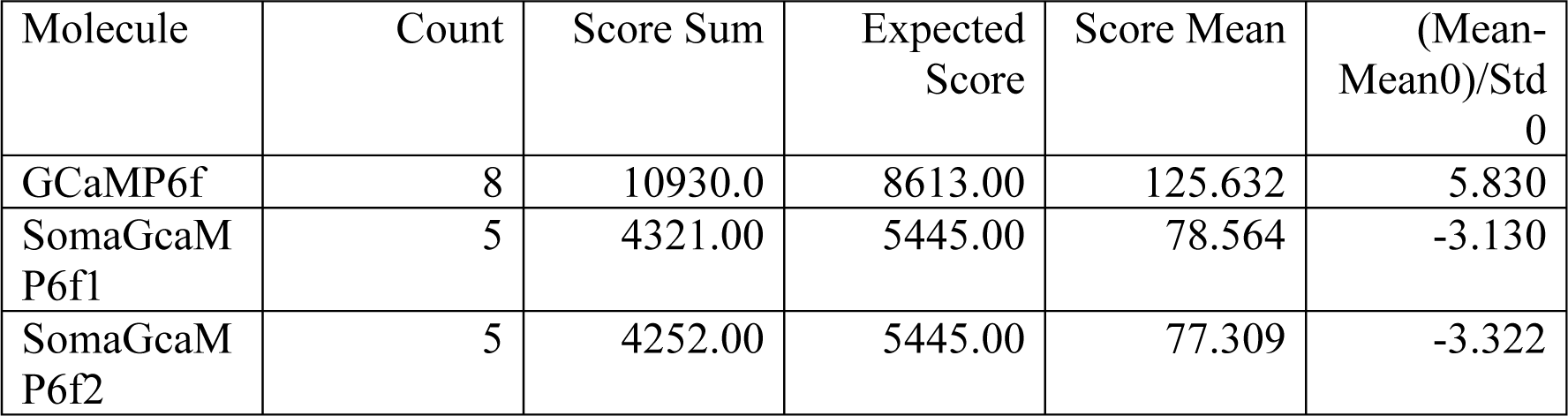

1-Way Test, ChiSquare Approximation

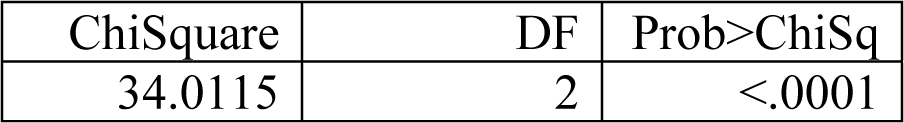

Nonparametric Comparisons With Control Using Steel Method

Control Group = GCaMP6f

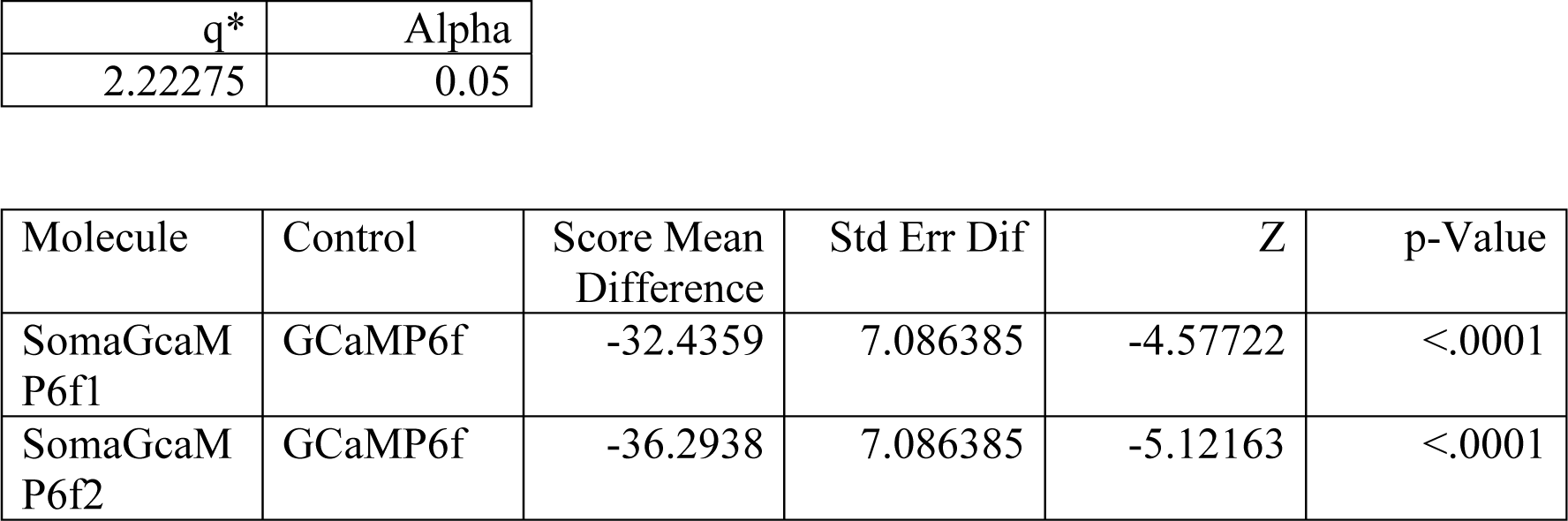

**Supplemental Table 4**: Statistical analysis for Figure 2.

Figure 2A - brightness

Brightness values for GCaMP6f, GCaMP6f-NLS, SomaGCaMP6f1 and SomaGCaMP6f2 (n = 8 cells from 2 cultures for GCaMP6f; n = 7 cells from 2 cultures for SomaGCaMP6f1; n = 5 cells from 2 cultures for SomaGCaMP6f2; n = 7 cells from 2 cultures for GCaMP6f-NLS).

Wilcoxon / Kruskal-Wallis Tests (Rank Sums)

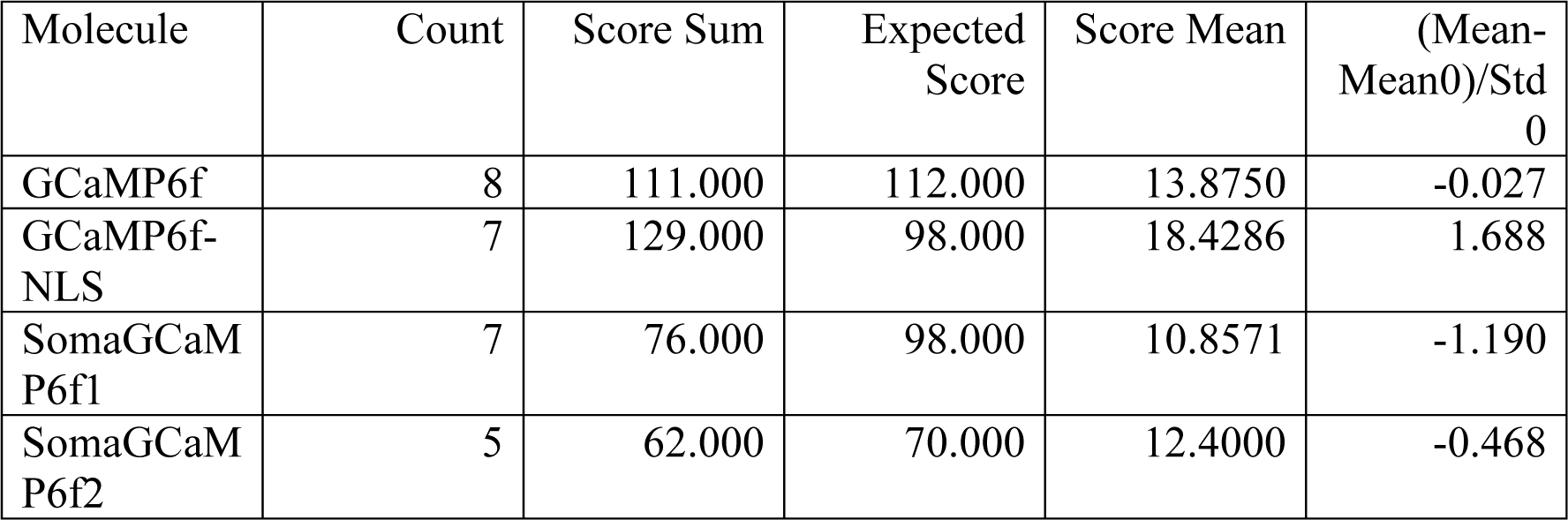

1-Way Test, ChiSquare Approximation

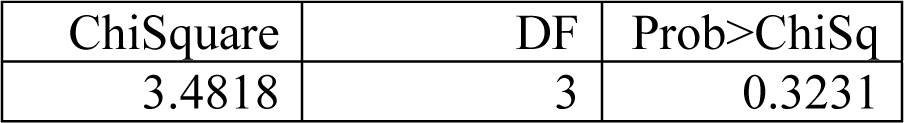

Nonparametric Comparisons With Control Using Steel Method

Control Group = GCaMP6f

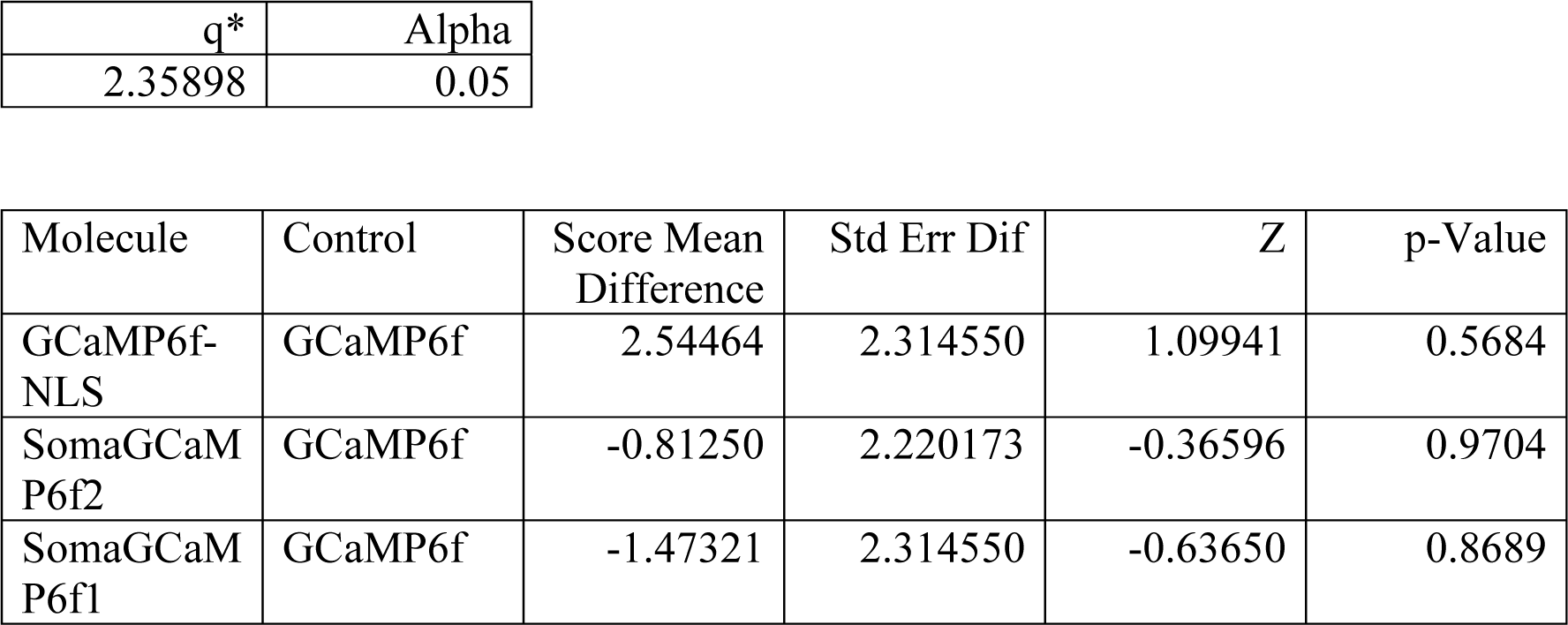

Figure 2C - df/f_0_

df/f_0_ for GCaMP6f, GCaMP6f-NLS, SomaGCaMP6f1 and SomaGCaMP6f2 (n = 8 cells from 2 cultures for GCaMP6f; n = 5 cells from 2 cultures for SomaGCaMP6f1; n = 7 cells from 2 cultures for SomaGCaMP6f2; n = 8 cells from 2 cultures for GCaMP6f-NLS).

Wilcoxon / Kruskal-Wallis Tests (Rank Sums)

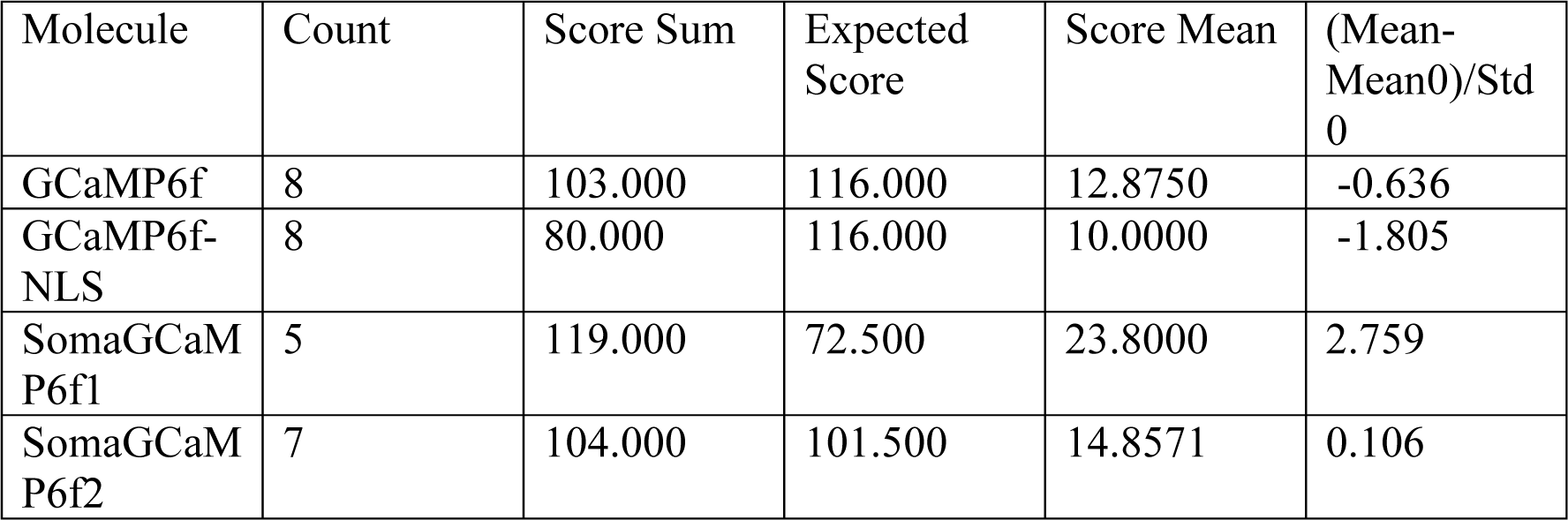

1-Way Test, ChiSquare Approximation

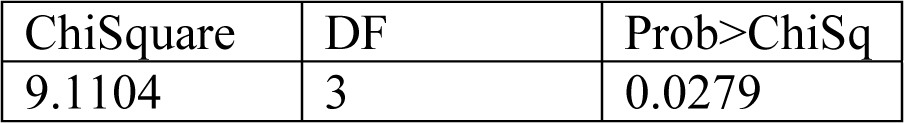

Nonparametric Comparisons With Control Using Steel Method

Control Group = GCaMP6f

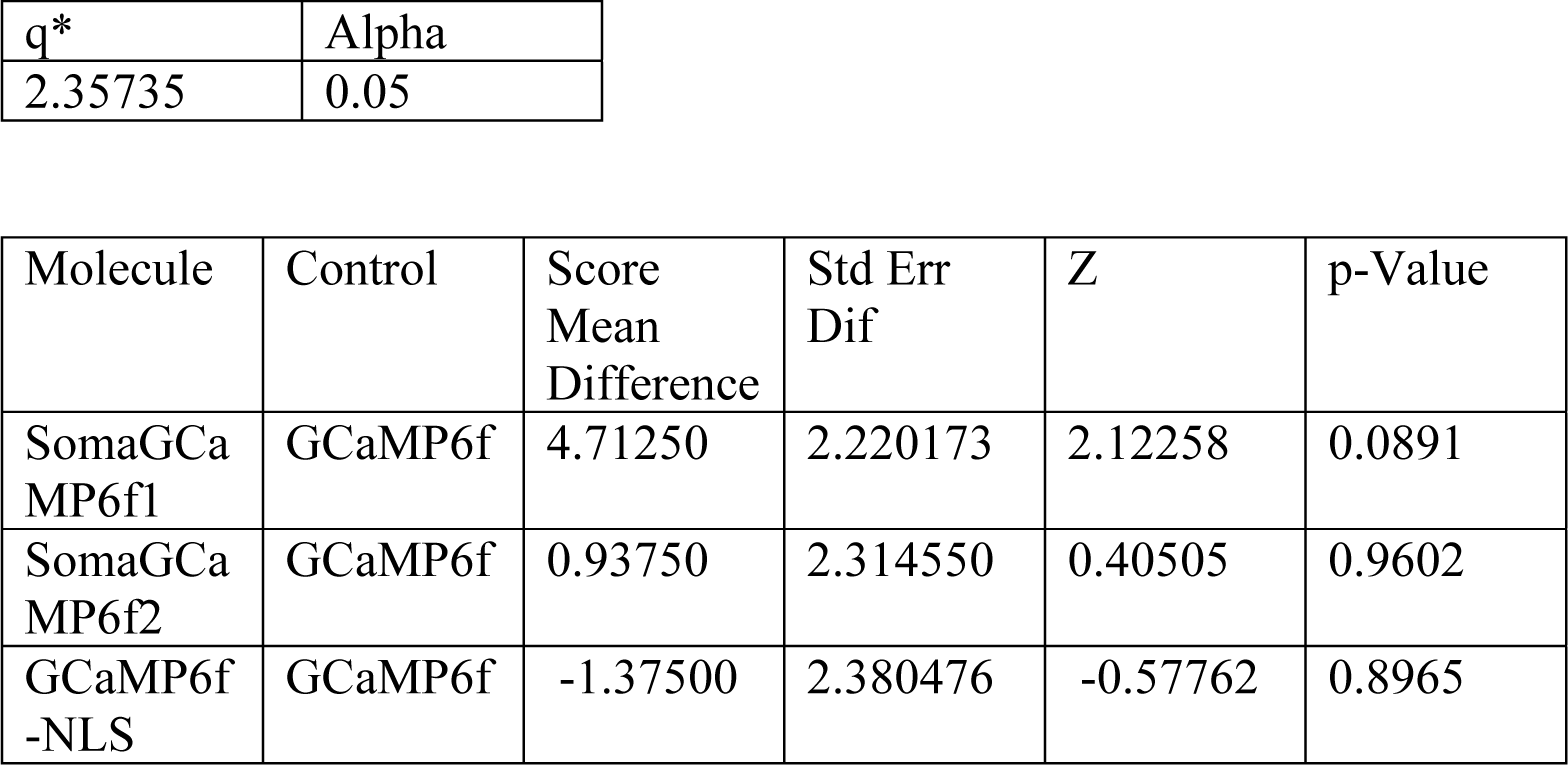

Figure 2D - SNR

Signal to noise ratio (SNR) for GCaMP6f, GCaMP6f-NLS, SomaGCaMP6f1 and SomaGCaMP6f2 (n = 8 cells from 2 cultures for GCaMP6f; n = 5 cells from 2 cultures for SomaGCaMP6f1; n = 7 cells from 2 cultures for SomaGCaMP6f2; n = 8 cells from 2 cultures for GCaMP6f-NLS).

Wilcoxon / Kruskal-Wallis Tests (Rank Sums)

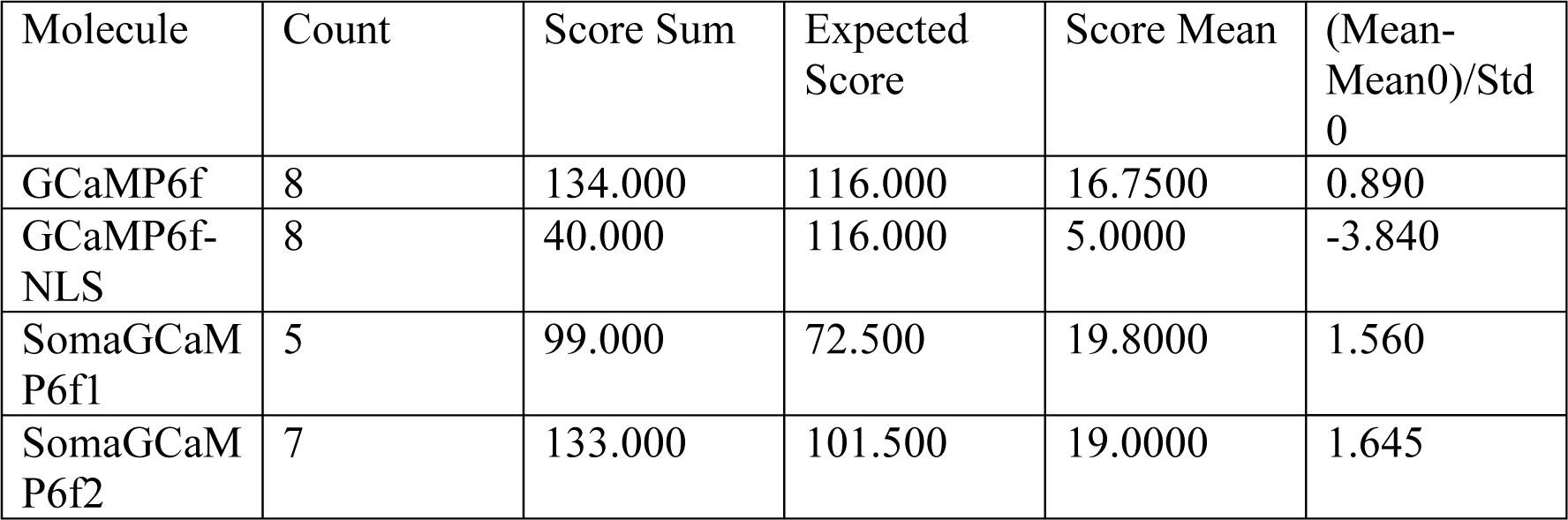

1-Way Test, ChiSquare Approximation

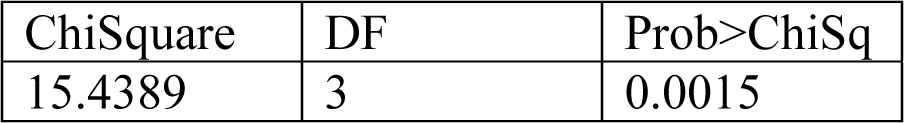

Nonparametric Comparisons With Control Using Steel Method

Control Group = GCaMP6f

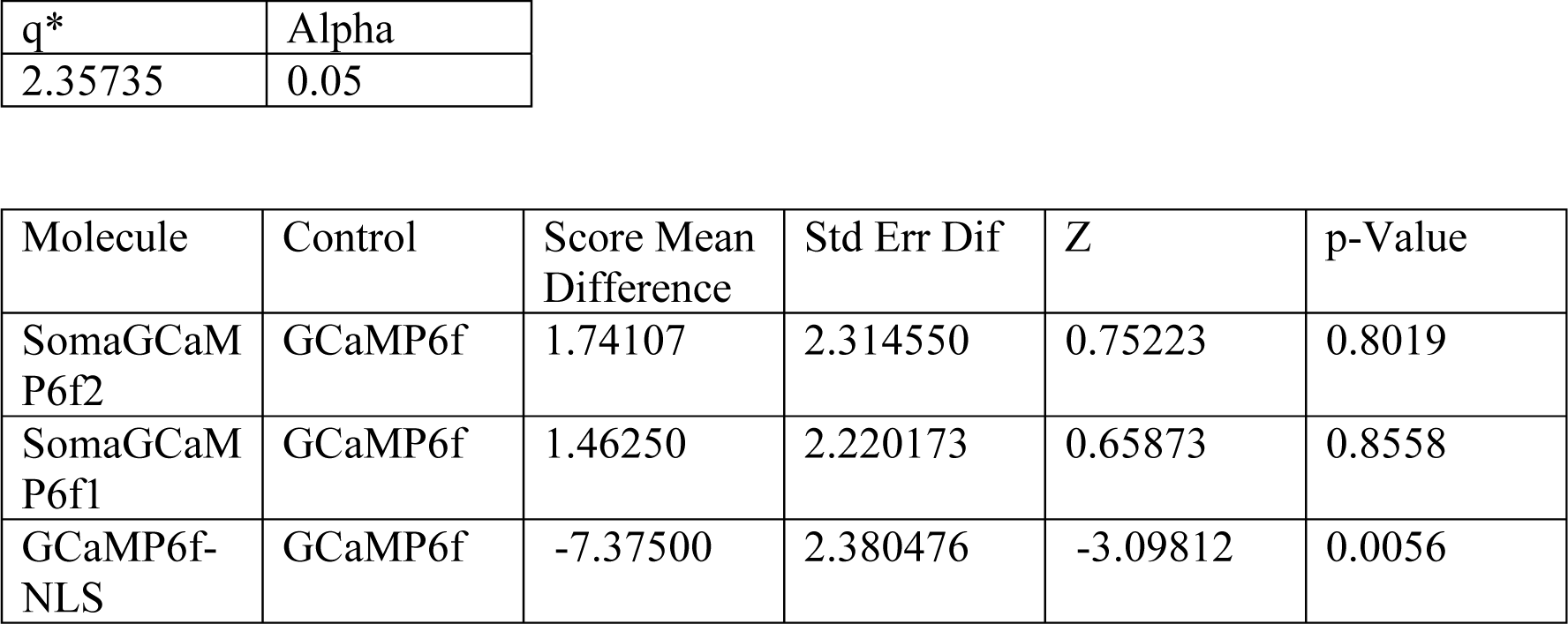

Figure 2E - Ton

Time constant for signal rise (Ton) for GCaMP6f, GCaMP6f-NLS, SomaGCaMP6f1 and SomaGCaMP6f2 (n = 8 cells from 2 cultures for GCaMP6f; n = 5 cells from 2 cultures for SomaGCaMP6f1; n = 6 cells from 2 cultures for SomaGCaMP6f2; n = 8 cells from 2 cultures for GCaMP6f-NLS).

Wilcoxon / Kruskal-Wallis Tests (Rank Sums)

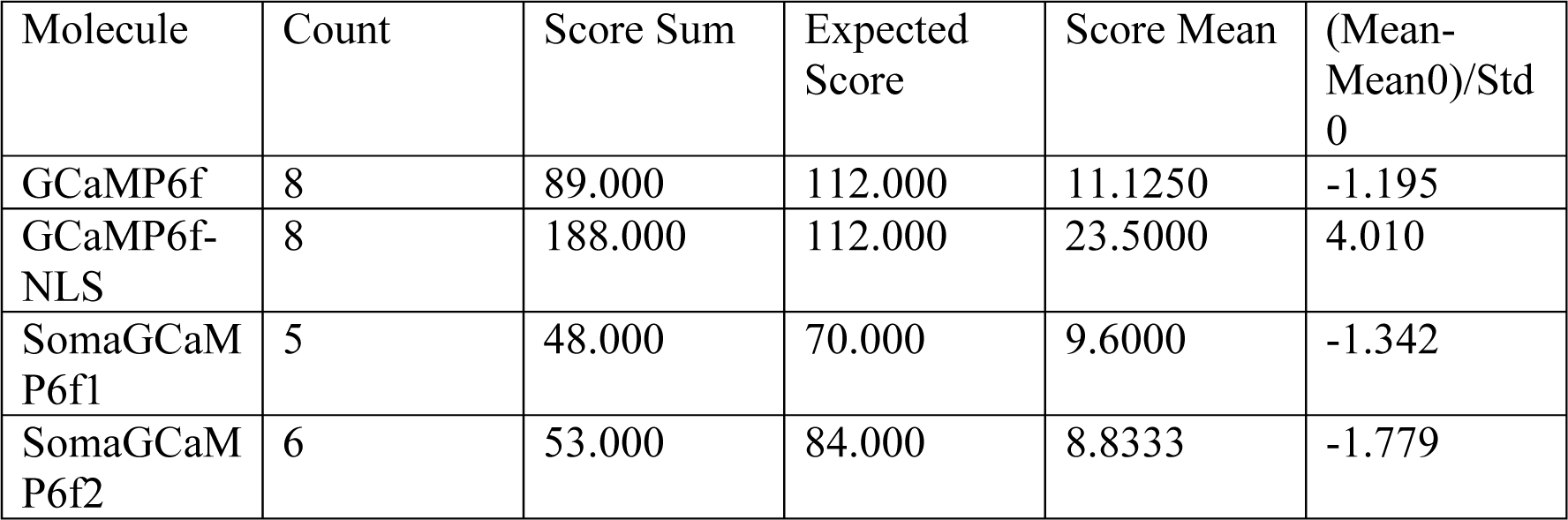

1-Way Test, ChiSquare Approximation

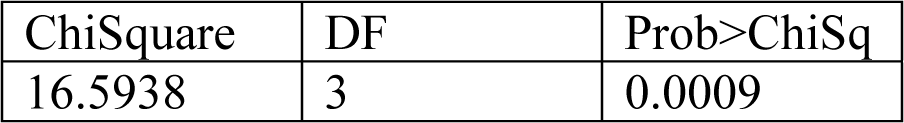

Nonparametric Comparisons With Control Using Steel Method

Control Group = GCaMP6f

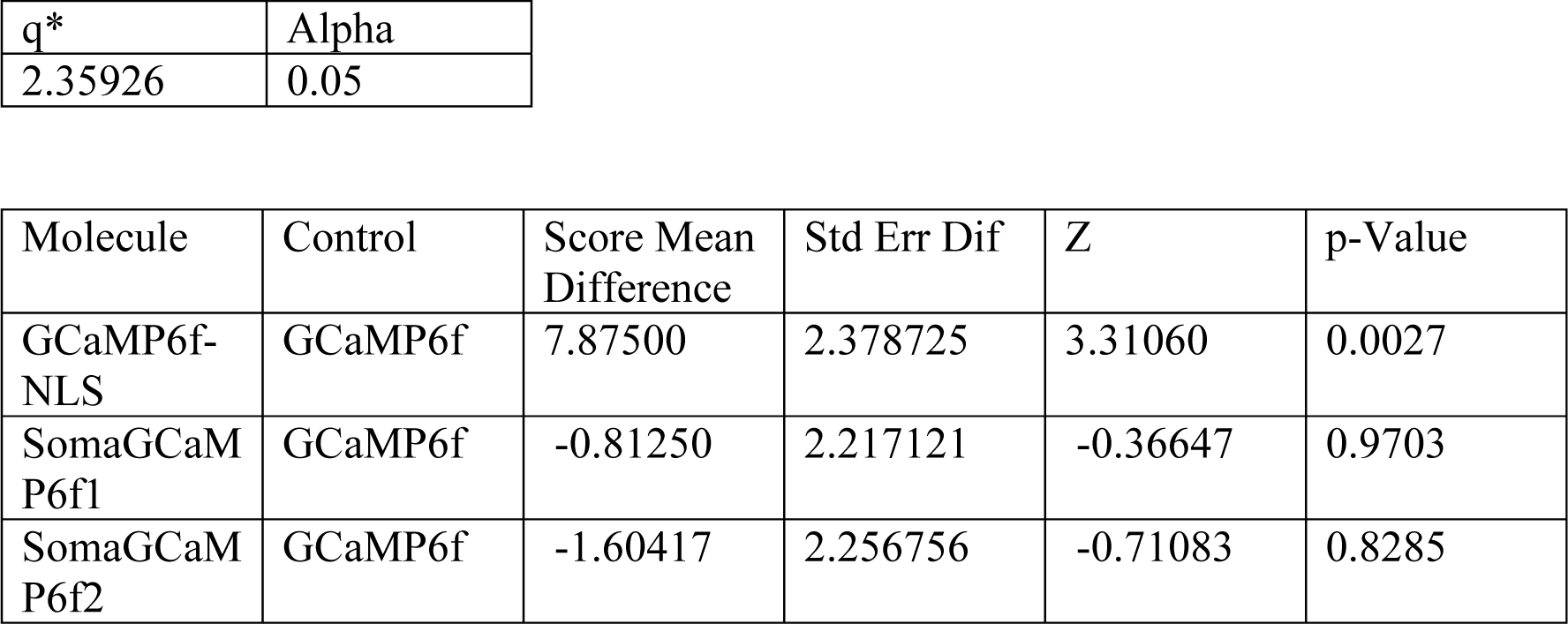

Figure 2F - Toff

Time constant for signal decay (Toff) for GCaMP6f, GCaMP6f-NLS, SomaGCaMP6f1 and SomaGCaMP6f2 (n = 7 cells from 2 cultures for GCaMP6f; n = 5 cells from 2 cultures for SomaGCaMP6f1; n = 7 cells from 2 cultures for SomaGCaMP6f2; n = 8 cells from 2 cultures for GCaMP6f-NLS).

Wilcoxon / Kruskal-Wallis Tests (Rank Sums)

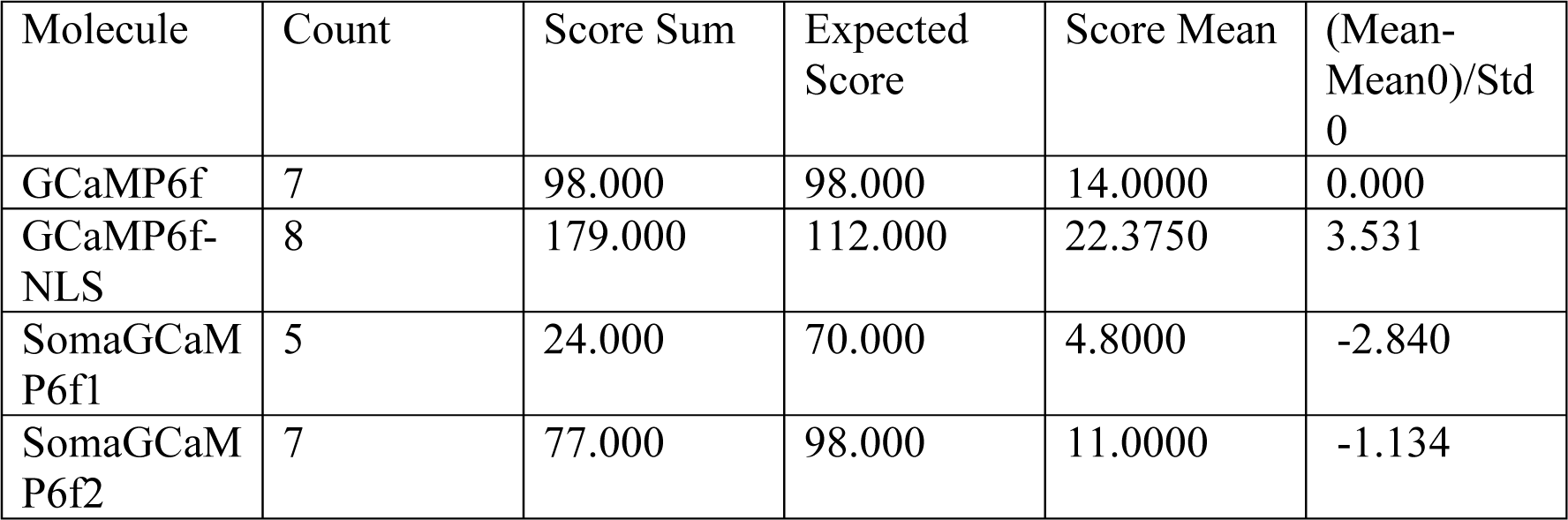

1-Way Test, ChiSquare Approximation

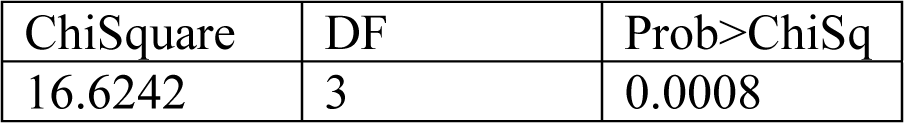

Nonparametric Comparisons With Control Using Steel Method

Control Group = GCaMP6f

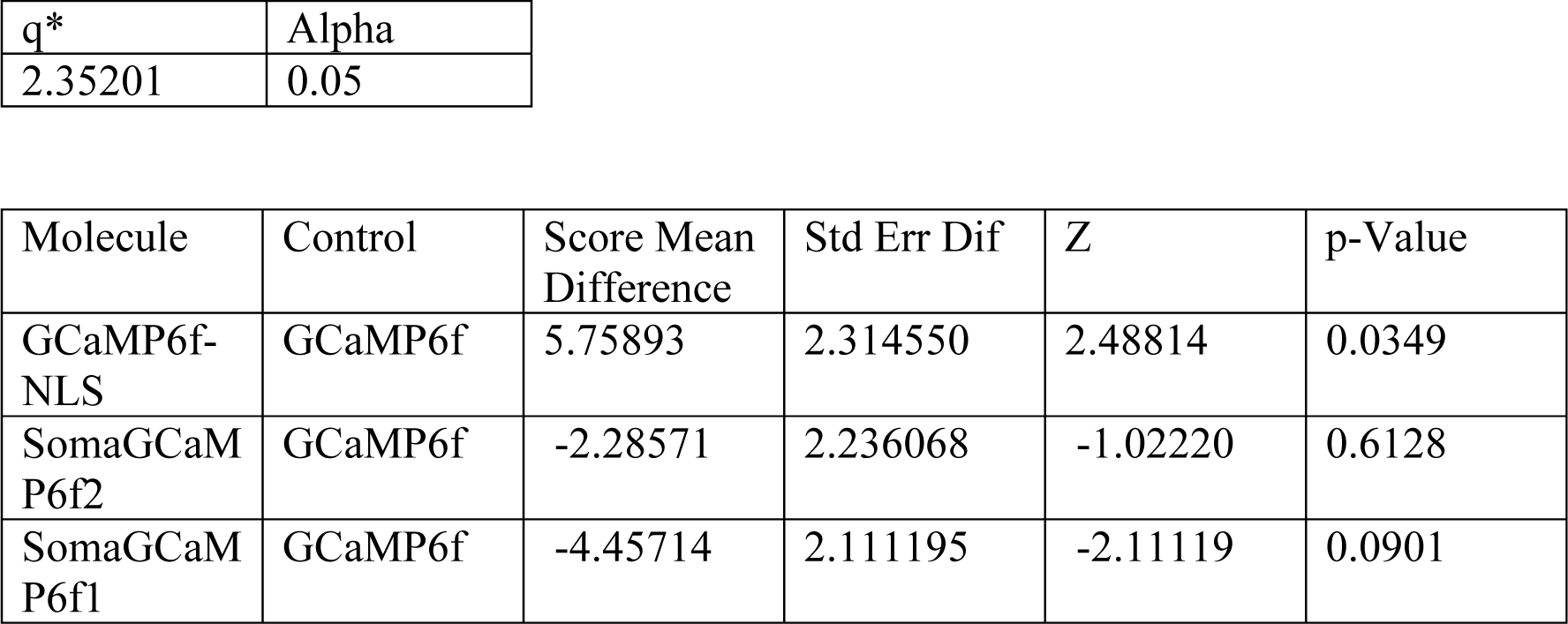

**Supplemental Table 5**: Statistical analysis for Figure 3.

<u>Figure 3A, B</u> Wilcoxon rank sum test for the expression density (number of expressing cells per 10^6^ um^3^) in the visual cortex of the slices from *in utero* electroporation between GCaMP6f and SomaGCaMP6f1 (n = 3 slices from 3 mice for GCaMP6f; n = 3 slices from 3 mice for SomaGCaMP6f1).

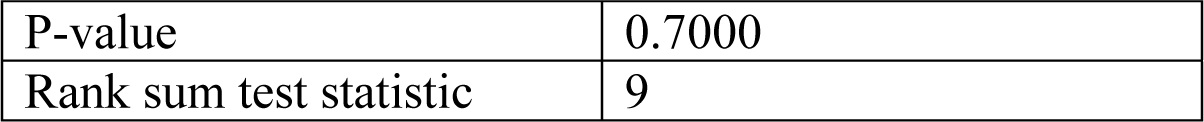

<u>Figure 3C</u> Wilcoxon rank sum test for the brightness of neurons in slices between GCaMP6f and SomaGCaMP6f1 (n = 7 neurons from 2 slices from 2 mice for GCaMP6f; n = 22 neurons from 6 slices from 3 mice for SomaGCaMP6f1), with light power adjusted to make them equal.

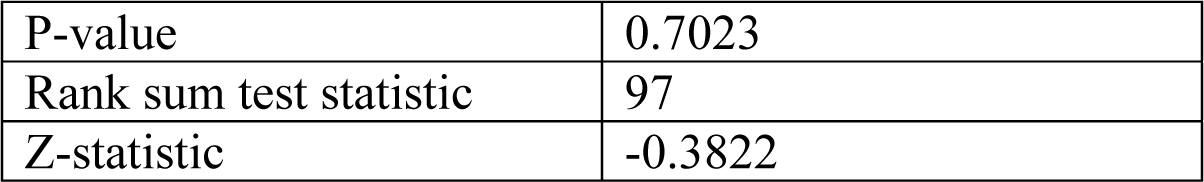

<u>Figure 3D</u> Two-sample Kolmogorov-Smirnov (K-S) test of normalized neurite brightness between GCaMP6f and SomaGCaMP6f1 (n = 11 neurites from 6 neurons from 2 slices from 2 mice for GCaMP6f; n = 11 neurites from 6 neurons from 2 slices from 2 mice for SomaGCaMP6f1).

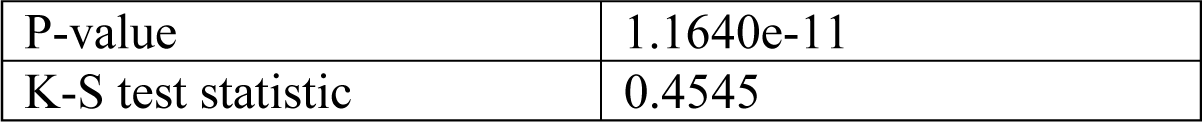

<u>Figure 3E</u> Wilcoxon rank sum test for the df/f_0_ of somata of neurons in slices between GCaMP6f and SomaGCaMP6f1 following an action potential (n = 14 APs from 3 neurons from 3 slices from 2 mice for GCaMP6f; n = 6 APs from 3 neurons from 3 slices from 3 mice for SomaGCaMP6f1).

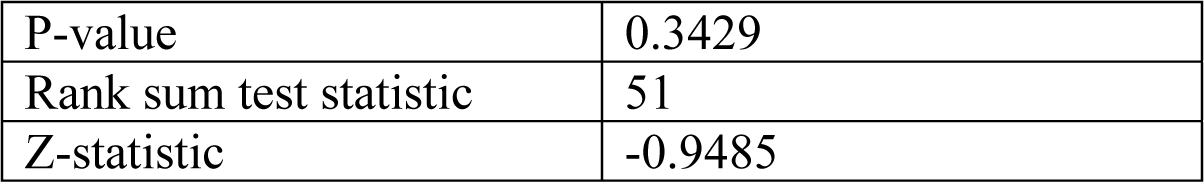

<u>Figure 3F</u> Wilcoxon rank sum test for the signal to noise ratio (SNR) of somata of neurons in slices between GCaMP6f and SomaGCaMP6f1 following an action potential (n = 14 APs from 3 neurons from 3 slices from 2 mice for GCaMP6f; n = 6 APs from 3 neurons from 3 slices from 3 mice for SomaGCaMP6f1).

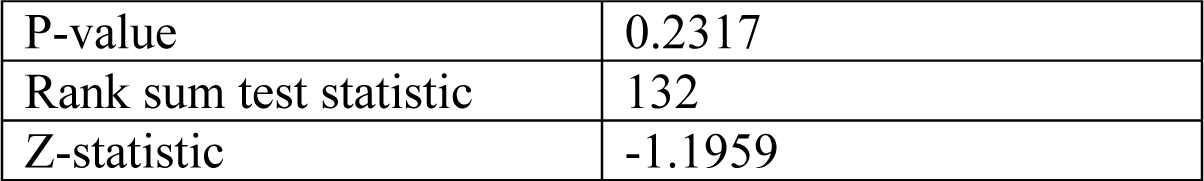

<u>Figure 3I</u> Wilcoxon rank sum test for the number of action potentials per minute in neurons in slices between GCaMP6f and SomaGCaMP6f1 following an action potential (n = 8 neurons from 8 slices for GCaMP6f from 4 mice; n = 6 neurons from 6 slices for SomaGCaMP6f1 from 3 mice).

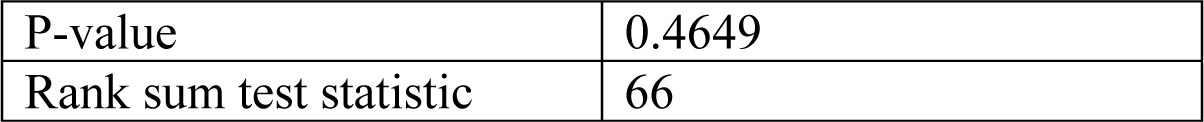

<u>Figure 3J</u> Wilcoxon rank sum test for the number of erroneous GCaMP-spikes per minute in neurons expressing either GCaMP6f or SomaGCaMP6f1 in slice (n = 8 neurons from 8 slices from 4 mice for GCaMP6f; n = 6 neurons from 6 slices from 3 mice for SomaGCaMP6f1).

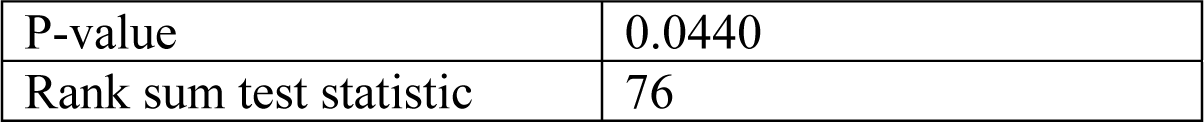

**Supplemental Table 6**: Statistical analysis for fish and mouse in vivo experiments (which include Figure 4 and 5).

Baseline brightness in zebrafish neurons in vivo, expressing GCaMP6f or SomaGCaMP6f1

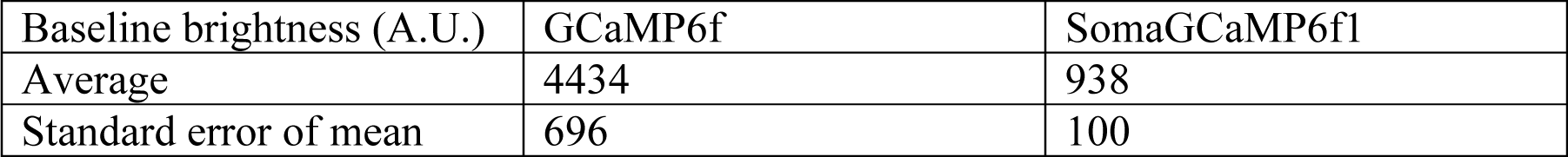

Wilcoxon rank sum test of baseline brightness between GCaMP6f (n = 25 neurons from 5 fish) and SomaGCaMP6f1 (n = 25 neurons from 5 fish):

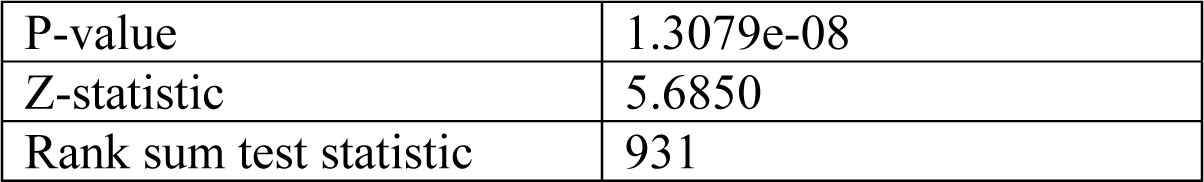

Figure 4D – Normalized brightness versus position along neurites of zebrafish neurons expressing GCaMP6f and SomaGCaMP6f1

Wilcoxon rank sum test of neurite brightness between GCaMP6f (5 neurites in 4 cells in 2 fish) and SomaGCaMP6f1 (8 neurites in 8 cells in 4 fish):

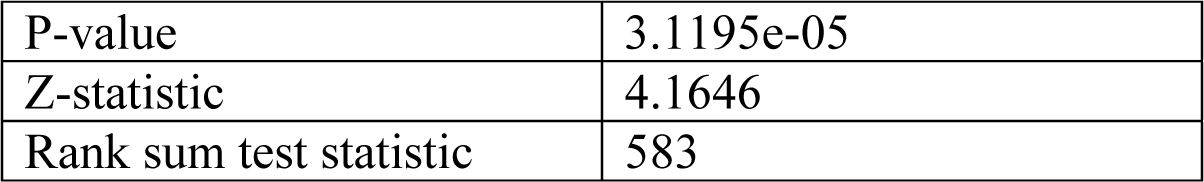

Figure 4F - df/f_0_ of somata of neurons in the visual area of zebrafish expressing GCaMP6f or SomaGCaMP6f1 in response to the moving grating

Wilcoxon rank sum test of df/f_0_ between GCaMP6f (n = 6 neurons from 3 fish) and SomaGCaMP6f1 (n = 5 neurons from 3 fish):

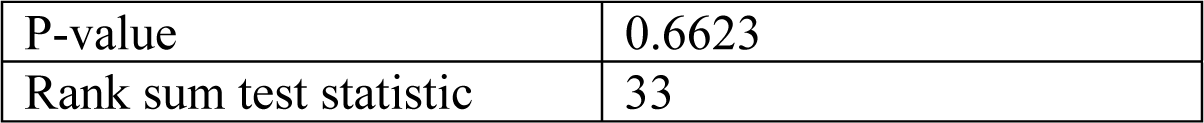

Figure 4G - Signal to noise ratio (SNR) of somata of neurons in the visual area of zebrafish expressing GCaMP6f or SomaGCaMP6f1 in response to the moving grating

Wilcoxon rank sum test of SNR between GCaMP6f (n = 6 neurons from 3 fish) and SomaGCaMP6f1 (n = 5 neurons from 3 fish):

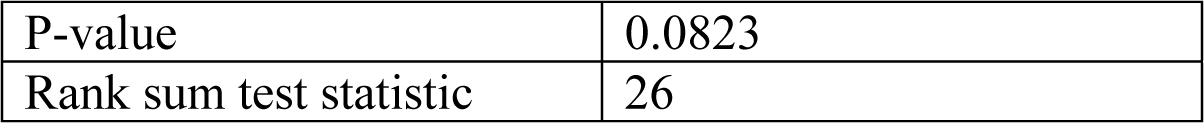

Figure 4J – df/f_0_ of somata of zebrafish neurons expressing GCaMP6f or SomaGCaMP6f1 and stimulated with 4AP

Wilcoxon rank sum test of df/f_0_ between GCaMP6f (n = 5 neurons from 2 fish) and SomaGCaMP6f1 (n = 5 neurons from 2 fish):

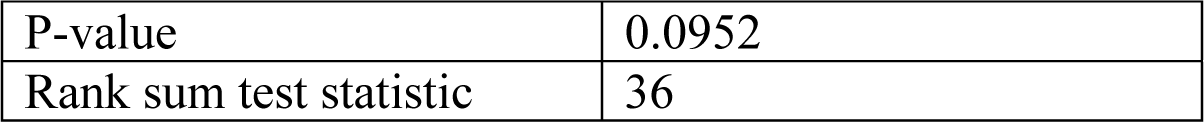

Figure 4K – signal to noise ratio (SNR) of somata of zebrafish neurons expressing GCaMP6f or SomaGCaMP6f1 and stimulated with 4AP

Wilcoxon rank sum test of SNR between GCaMP6f (n = 5 neurons from 2 fish) and SomaGCaMP6f1 (n = 5 neurons from 2 fish):

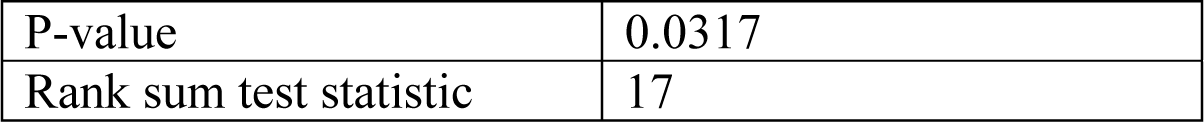

Figure 4M – correlations between cell pairs as a function of distance, between zebrafish neurons expressing GCaMP6f or SomaGCaMP6f1 and stimulated with 4AP

Two-dimensional Kolmogorov-Smirnov test between GCaMP6f and SomaGCaMP6f1 fish (n = 426 cells from 5 fish) and SomaGCaMP6f1 (n = 340 cells from 4 fish).

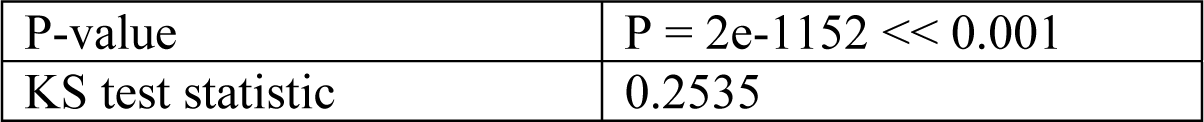

Figure 5E Wilcoxon rank sum test comparing session event rates between GCaMP6f and SomaGCaMP6f2 (n = 930 neurons from 7 GCaMP6f mice; n = 594 neurons from 4 SomaGCaMP6f2 mice).

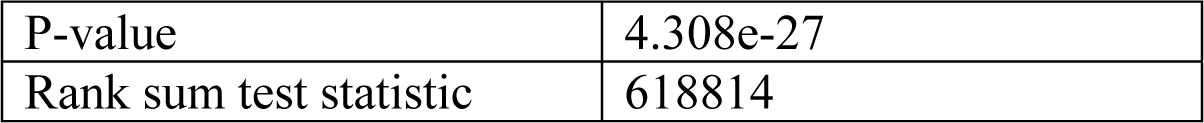

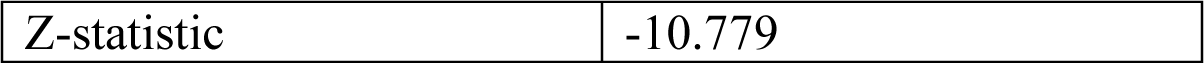

<u>Figure 5H</u> Wilcoxon rank sum test for the mean cell pairwise correlation strength between GCaMP6f and SomaGCaMP6f2 imaging sessions (n = 431985 cell-pairs from 7 GCaMP6f mice; n = 176121 cell-pairs from 4 SomaGCaMP6f2 mice.

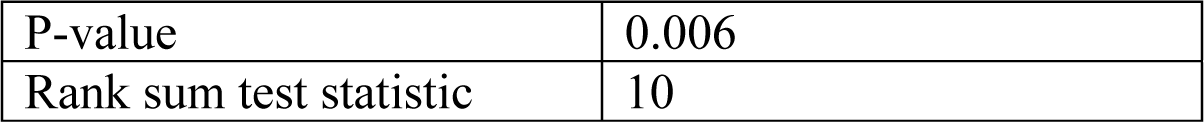

**Supplemental Table 7**: Statistical analysis for **Figure S1**.

Figure S1 K - df/f_0_ of different GCaMP6f targeting variants.

(n = 20 cells from 2 slices from 2 mice for each variant).

Wilcoxon / Kruskal-Wallis Tests (Rank Sums)

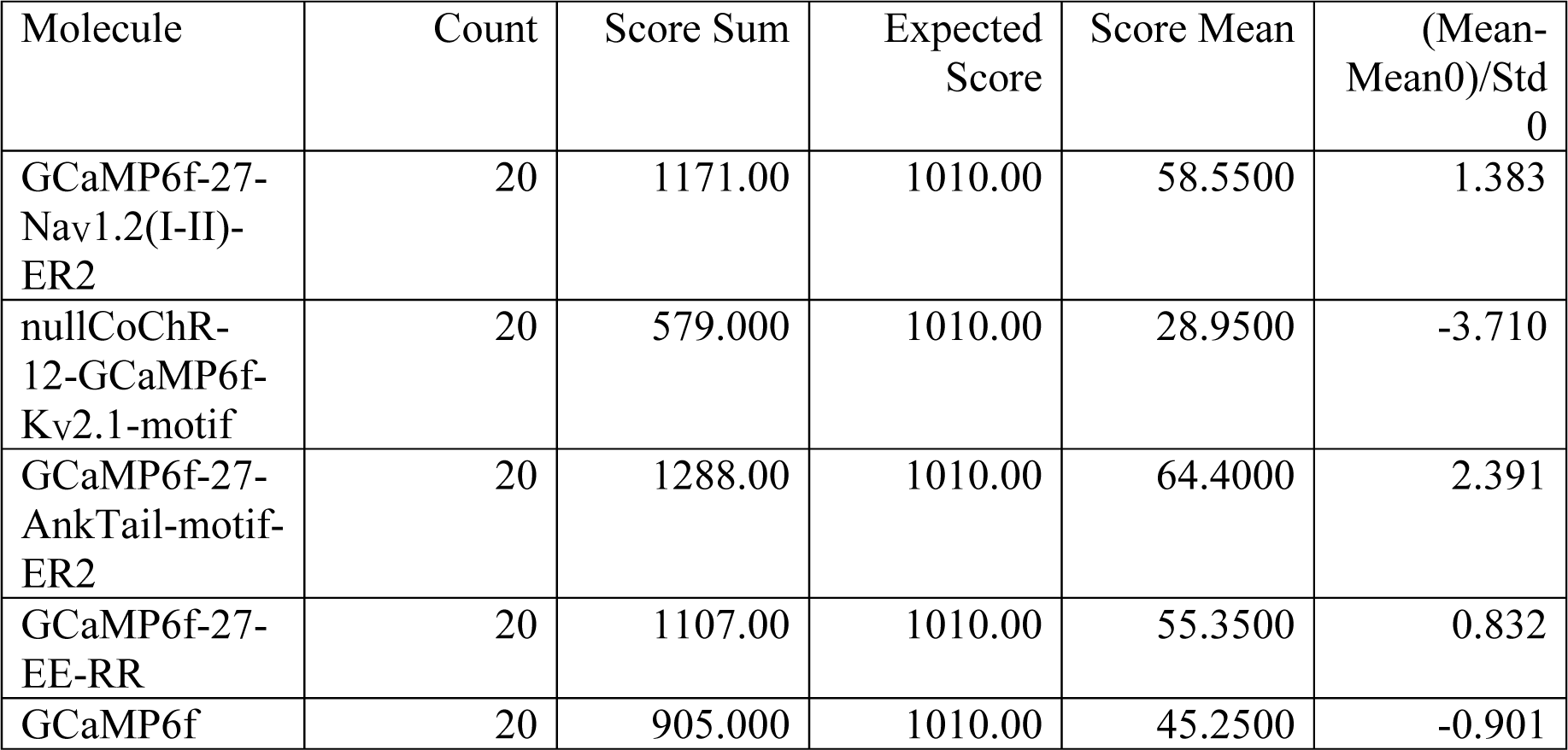

1-Way Test, ChiSquare Approximation

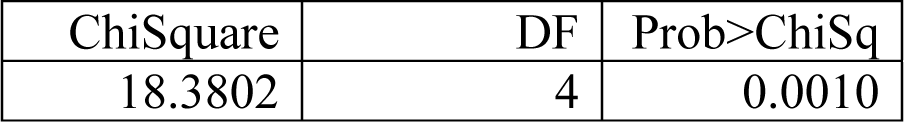

Nonparametric Comparisons with Control Using Steel Method

Control Group = GCaMP6f

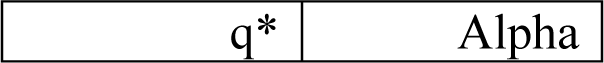

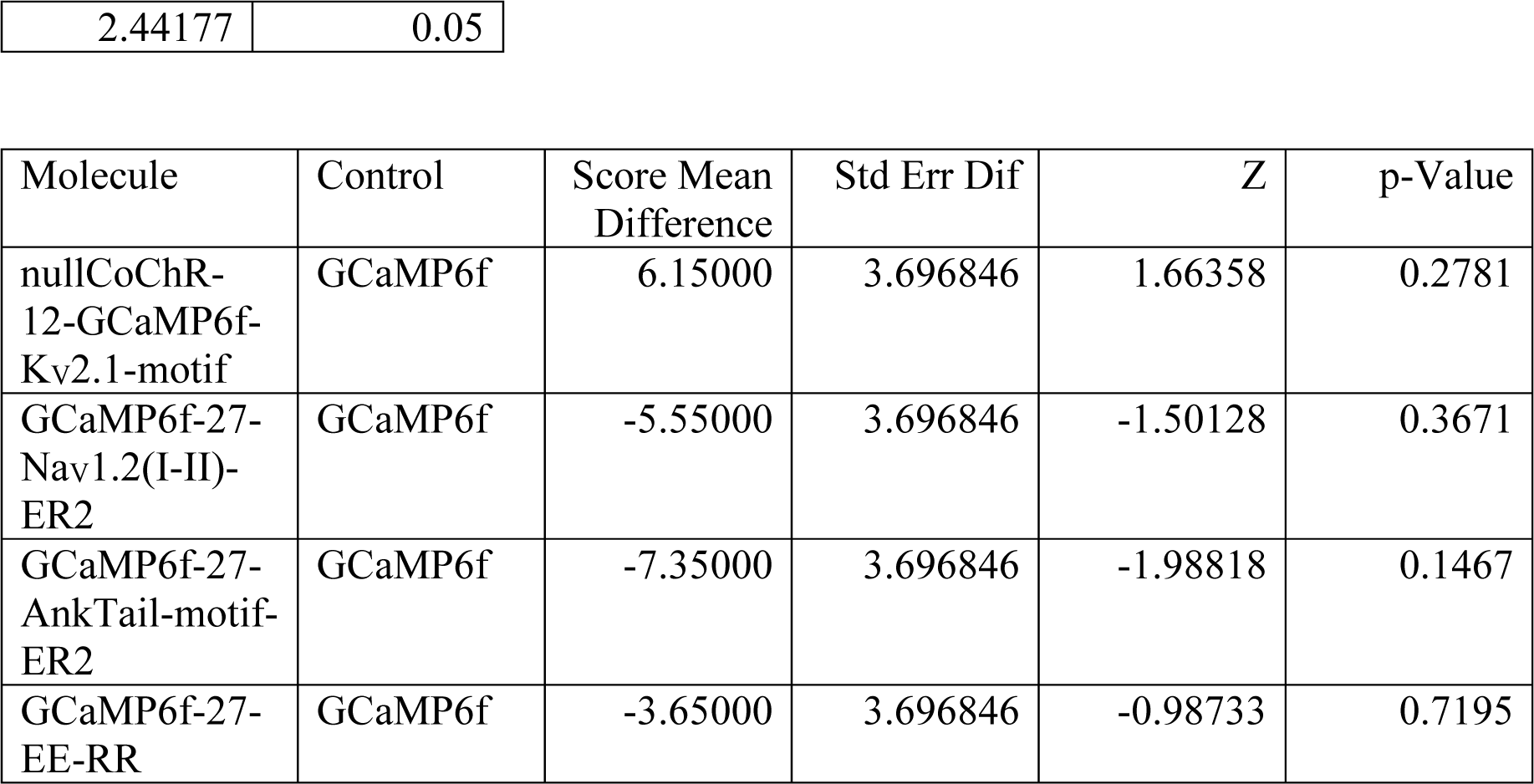

Figure S1 L - the ratio between df/f_0_ of the cell body and df/f_0_ of the neuropil for different GCaMP6f targeting variants.

(n = 20 cells from 2 slices from 2 mice for each variant).

Wilcoxon / Kruskal-Wallis Tests (Rank Sums)

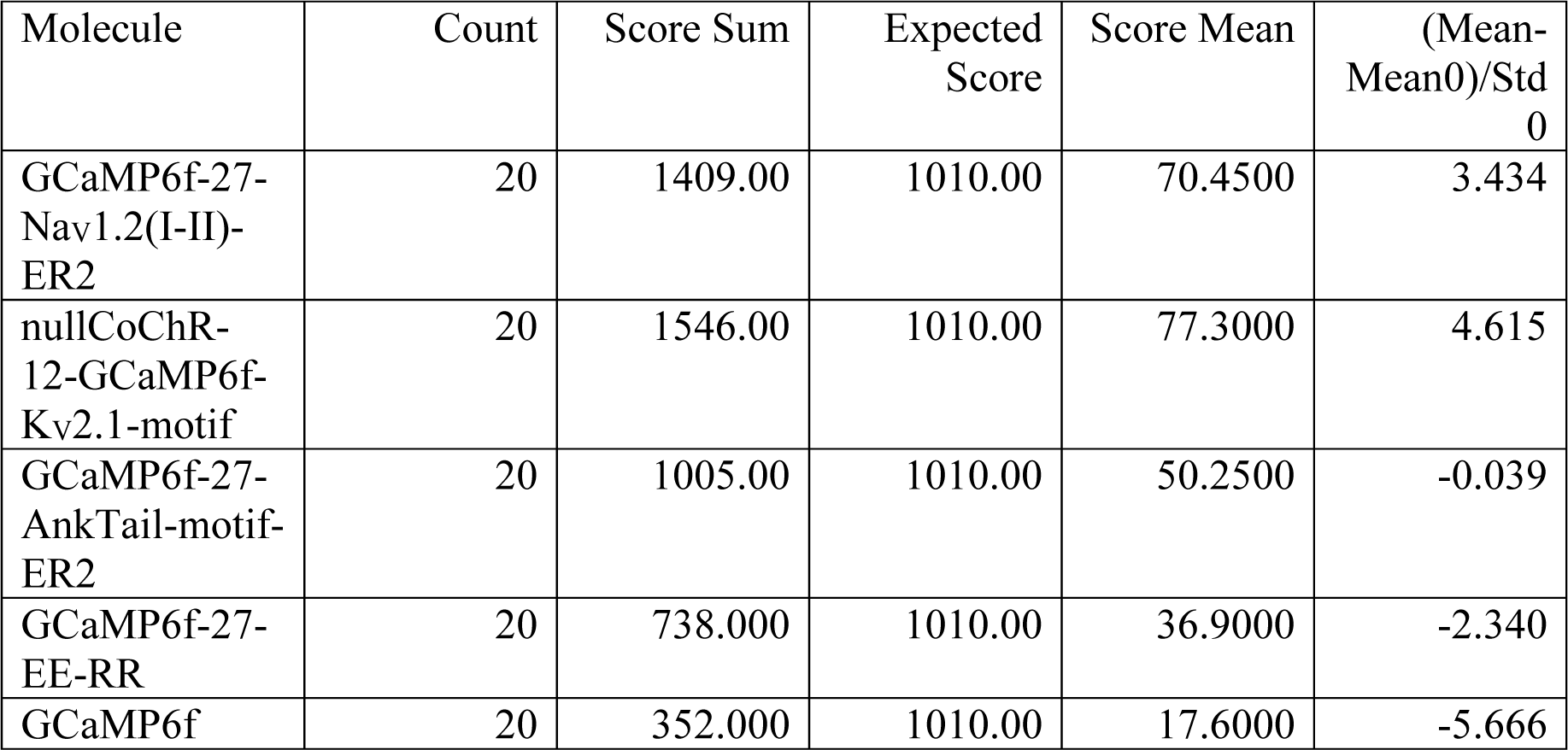

1-Way Test, ChiSquare Approximation

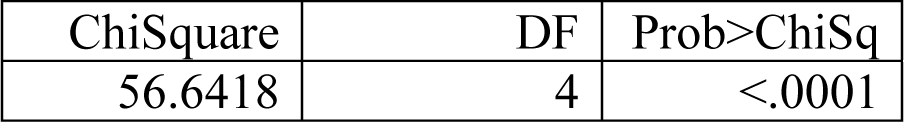

Nonparametric Comparisons With Control Using Steel Method

Control Group = GCaMP6f

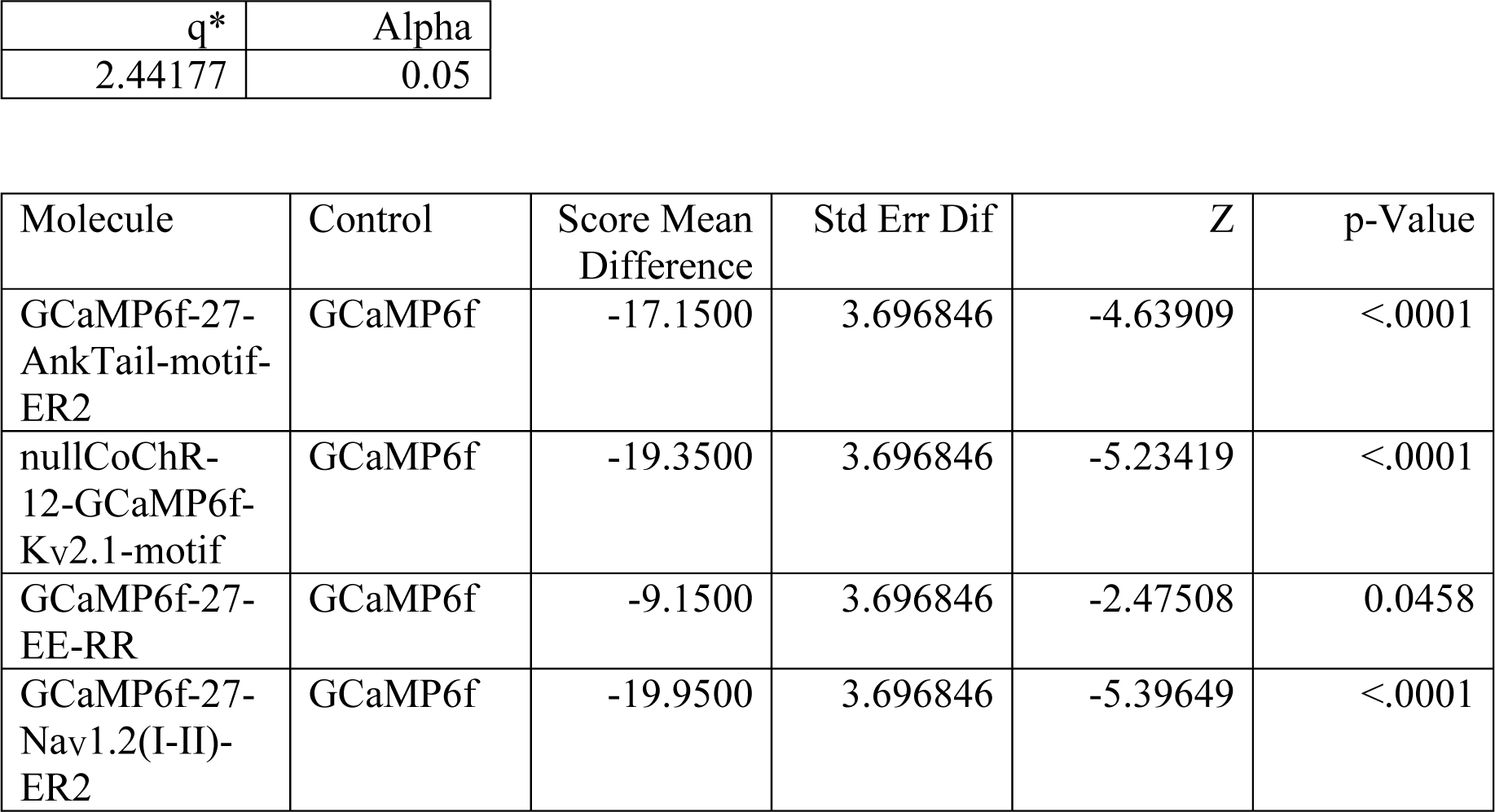

**Supplemental Table 8**: Statistical analysis for **Figure S2** – membrane properties

<u>Figure S2A</u>

Wilcoxon / Kruskal-Wallis Tests (Rank Sums)

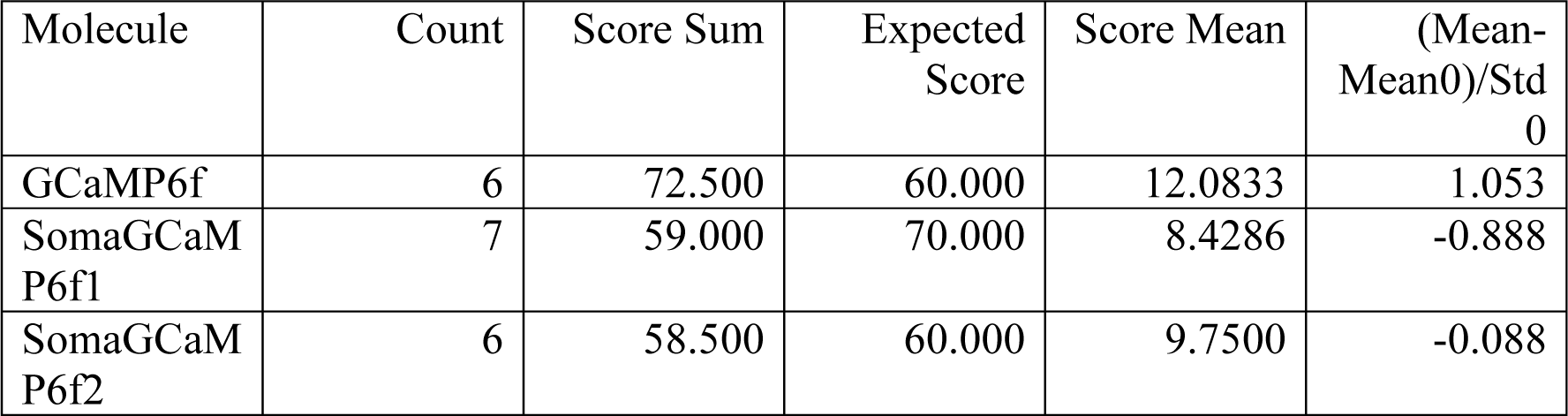

1-Way Test, ChiSquare Approximation

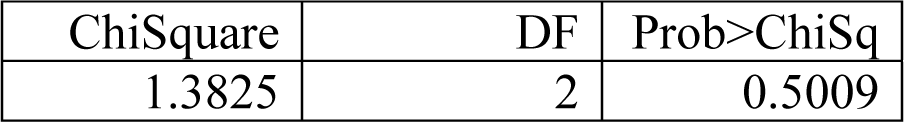

Nonparametric Comparisons With Control Using Steel Method

Control Group = GCaMP6f

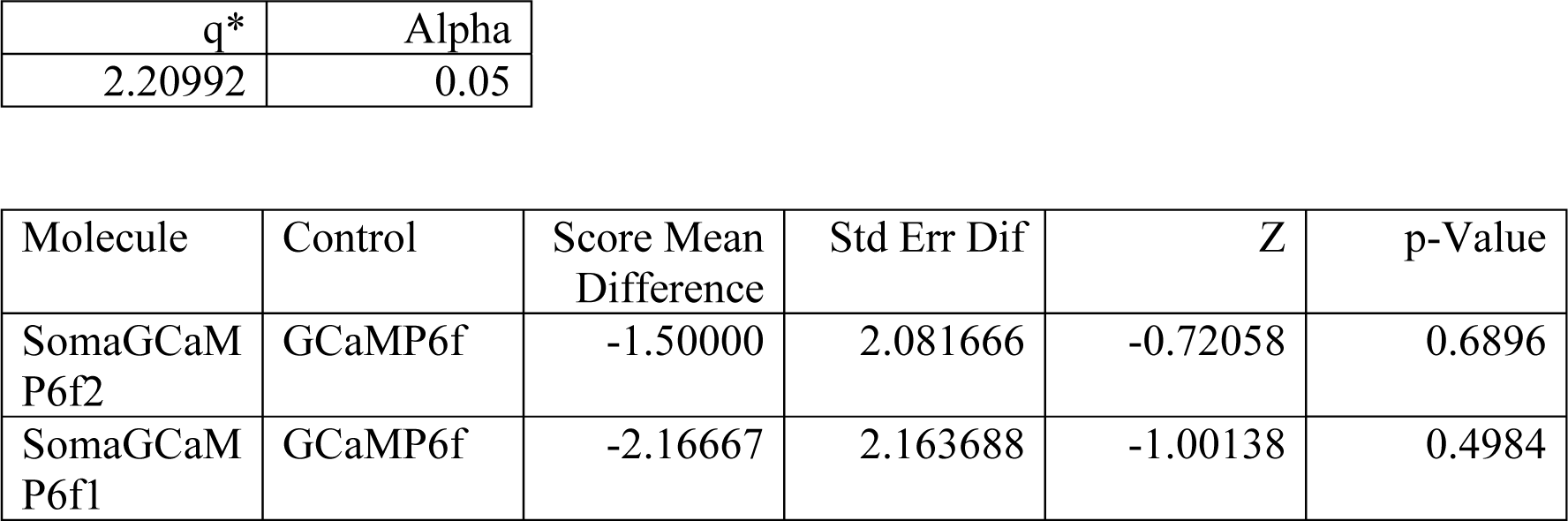

<u>Figure S2B</u>

Wilcoxon / Kruskal-Wallis Tests (Rank Sums)

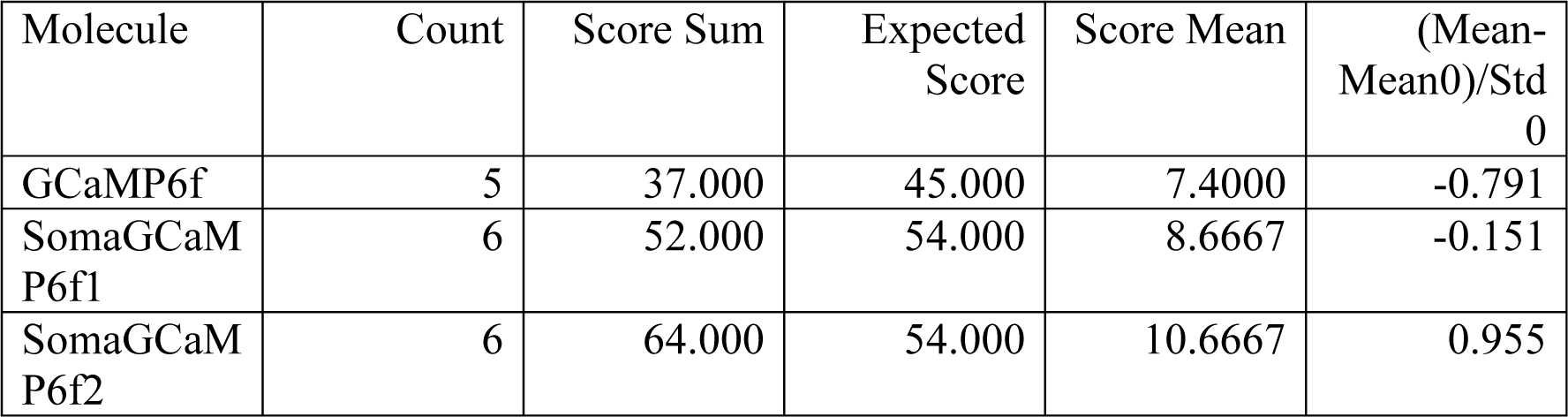

1-Way Test, ChiSquare Approximation

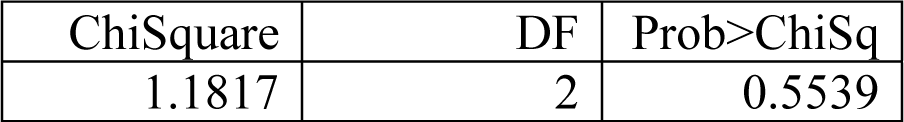

Small sample sizes. Refer to statistical tables for tests, rather than large-sample approximations. Nonparametric Comparisons With Control Using Steel Method

Control Group = GCaMP6f

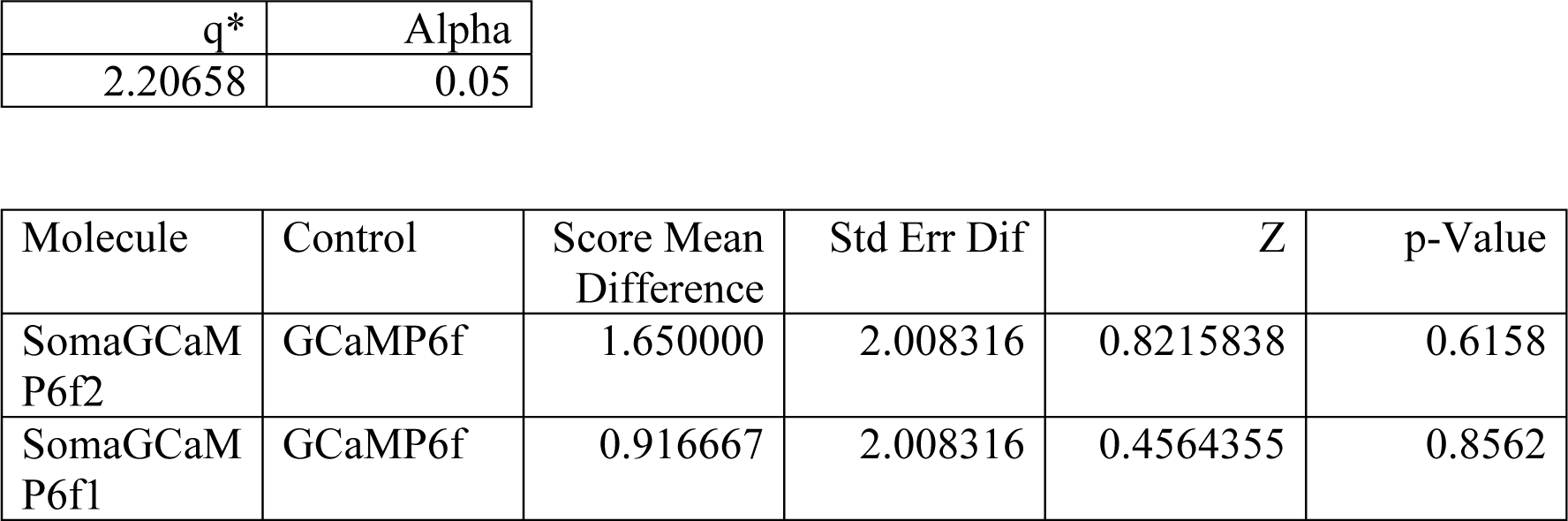

<u>Figure S2C</u>

Wilcoxon / Kruskal-Wallis Tests (Rank Sums)

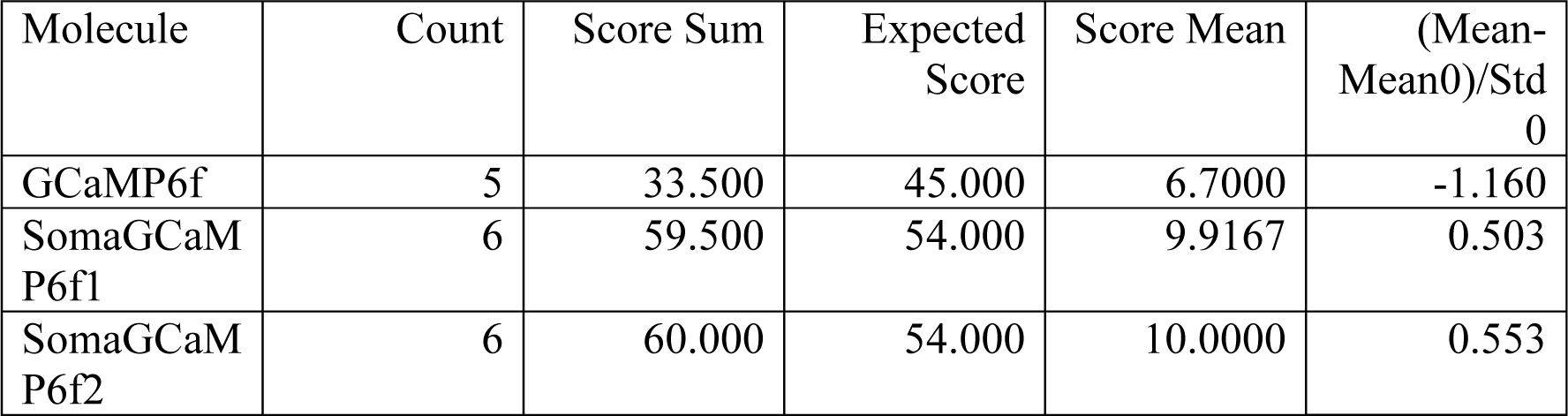

1-Way Test, ChiSquare Approximation

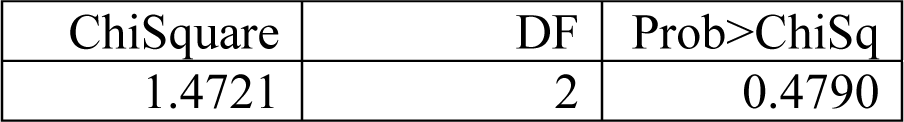

Small sample sizes. Refer to statistical tables for tests, rather than large-sample approximations. Nonparametric Comparisons With Control Using Steel Method

Control Group = GCaMP6f

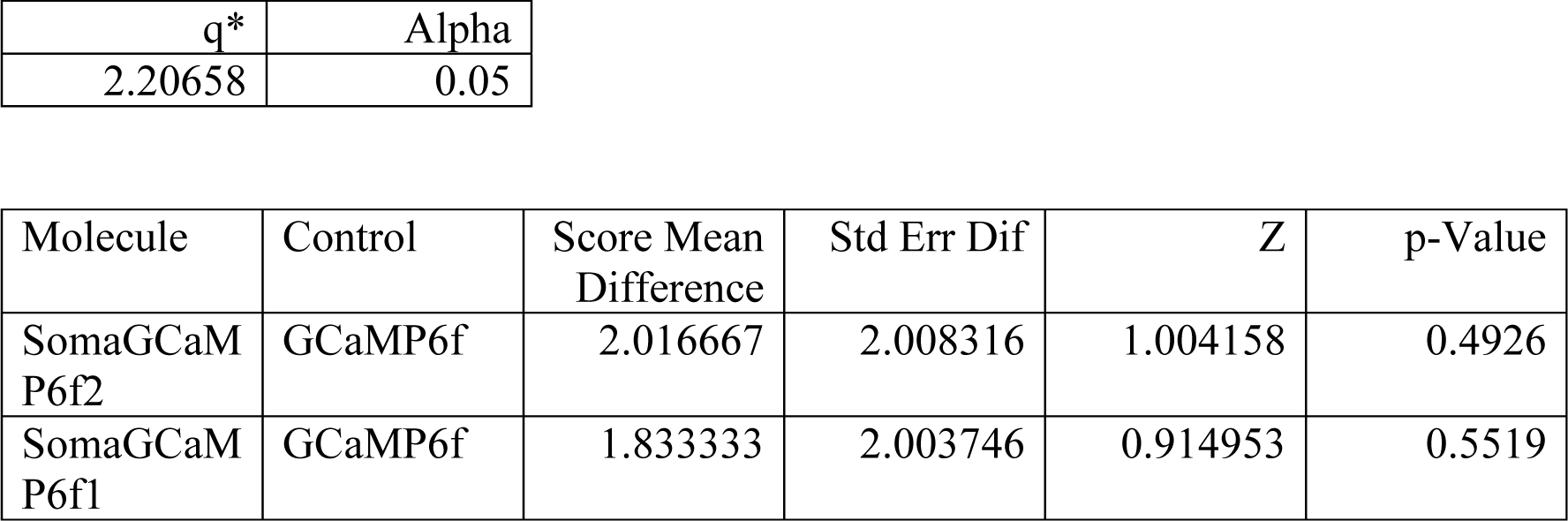

<u>Figure S2D</u>

Wilcoxon / Kruskal-Wallis Tests (Rank Sums)

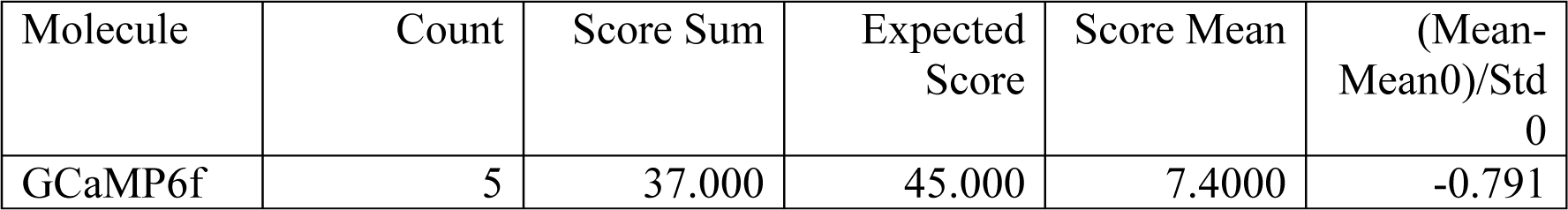

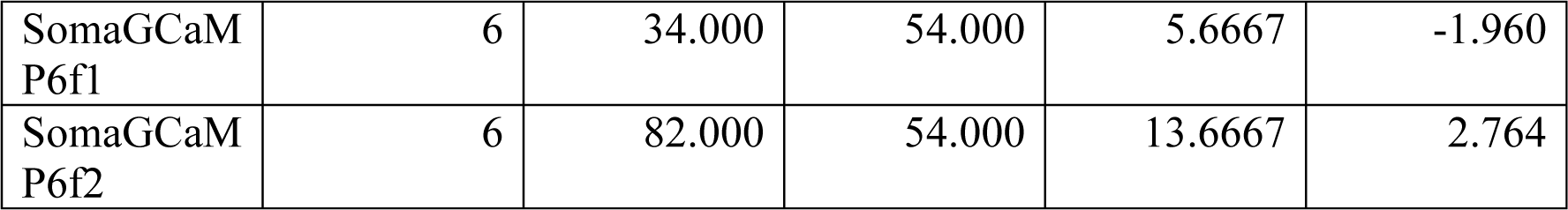

1-Way Test, ChiSquare Approximation

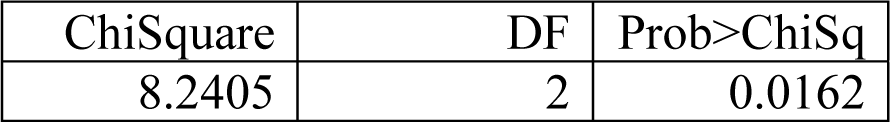

Small sample sizes. Refer to statistical tables for tests, rather than large-sample approximations. Nonparametric Comparisons With Control Using Steel Method

Control Group = GCaMP6f

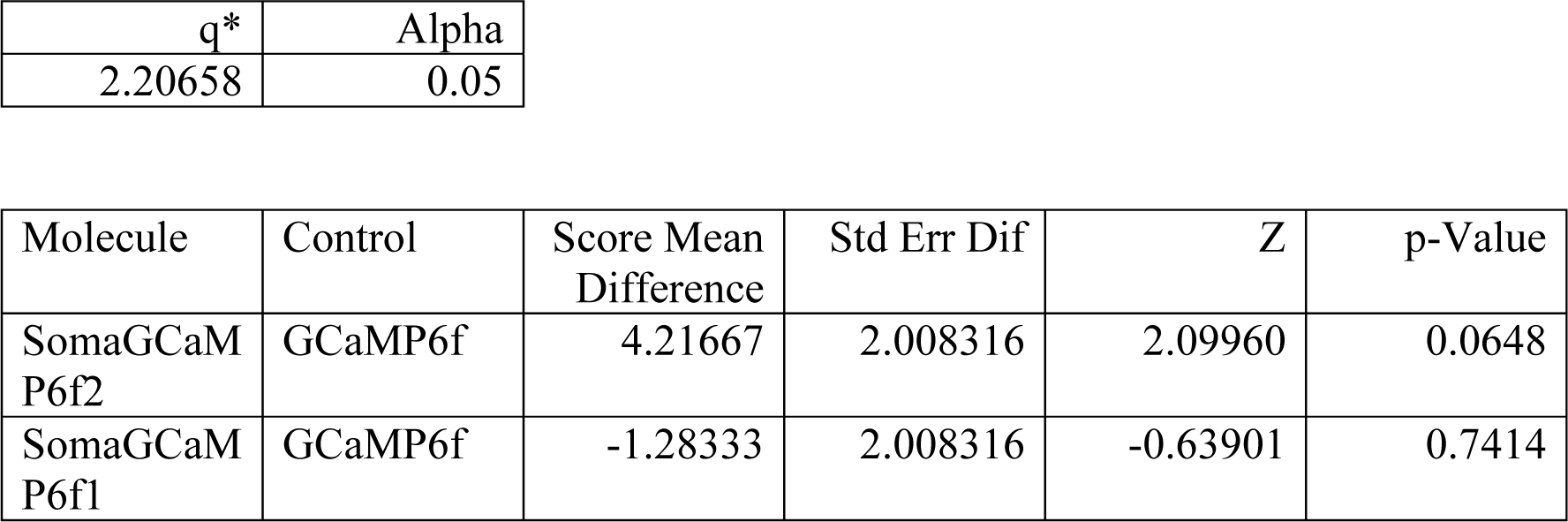

**Supplemental Table 9**: Statistical analysis for figures S3 – baseline brightness in mouse brain slice

Wilcoxon rank sum test of baseline brightness in slice between GCaMP6f and SomaGCaMP6f1 (n = 42 neurons from 4 slices from 2 GCaMP6f mice; n = 43 neurons from 8 slices from 3 SomaGCaMP6f1 mice).

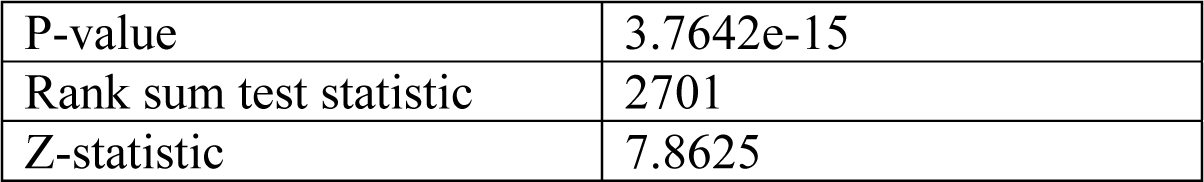

**Supplemental Table 10**: Statistical analysis for **Figure S4** – sensitivity for multiple action potentials, temporal dynamics and event rate for GCaMP6f and SomaGCaMP6f1.

<u>Figure S4A</u>

Bonferroni-corrected Wilcoxon rank sum test of the df/f_0_ between GCaMP6f (n = 7 neurons from 5 slices from 2 mice) and SomaGCaMP6f1 (n = 5 neurons from 3 slices from 2 mice) expressing neurons. The overall significance level α was set to 0.05, and the significance level of each individual Wilcoxon rank sum test was α/4 = 0.0125. P values less than 0.0125 are highlighted in bold.

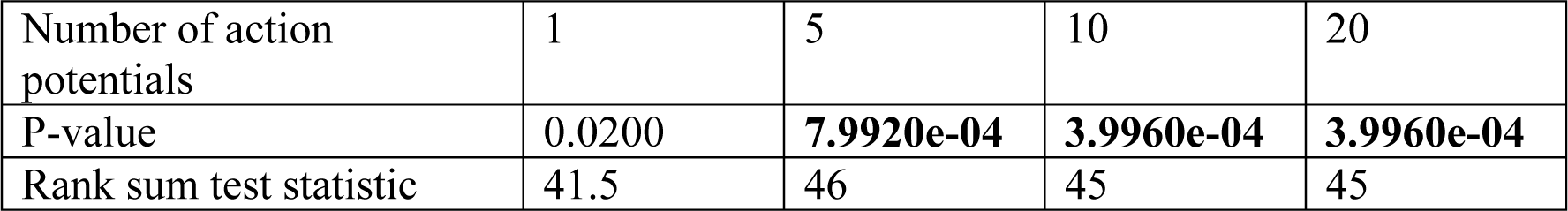

<u>Figure S4B</u>

Wilcoxon rank sum test of the τ_off_ of calcium spikes in slice during electrophysiological inducement of single action potentials between GCaMP6f and SomaGCaMP6f1(n = 3 neurons from 3 slices from 3 mice for GCaMP6f; n = 3 neurons from 3 slices from 3 mice for SomaGCaMP6f1).

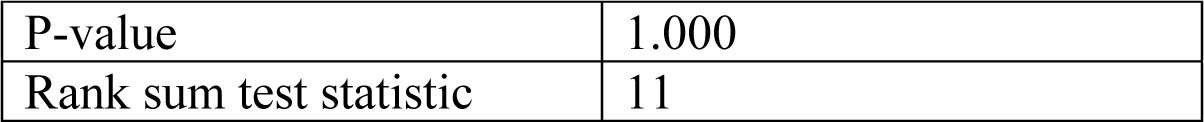

<u>Figure S4C</u>

Wilcoxon rank sum test the τ_off_ of calcium spikes in slice during 4-aminopyridine inducement of single action potentials between GCaMP6f and SomaGCaMP6f1 (n = 5 neurons from 5 slices from 4 mice for GCaMP6f; n = 5 neurons from 4 slices from 3 mice for SomaGCaMP6f1).

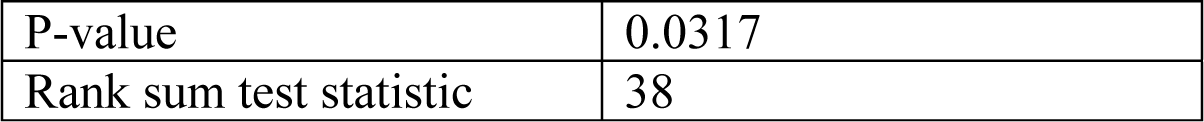

<u>Figure S4D</u>

Wilcoxon rank sum test of the event rate of calcium spikes per minute in slice between GCaMP6f and SomaGCaMP6f1 (n = 8 neurons from 8 slices for GCaMP6f from 4 mice; n = 6 neurons from 6 slices from 3 mice for SomaGCaMP6f1).

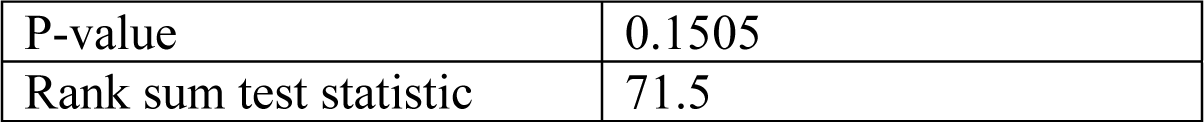

**Supplemental Table 11**: Statistical analysis for **Figure S5** – Temporal dynamics and calcium spike count for GCaMP6f and SomaGCaMP6f1 fish.

<u>Figure S5A</u>

Wilcoxon rank sum test of the τ_on_ of calcium spikes in zebrafish larva. (n = 96 cells from 3 GCaMP6f fish; n = 146 cells from 4 SomaGCaMP6f1 fish).

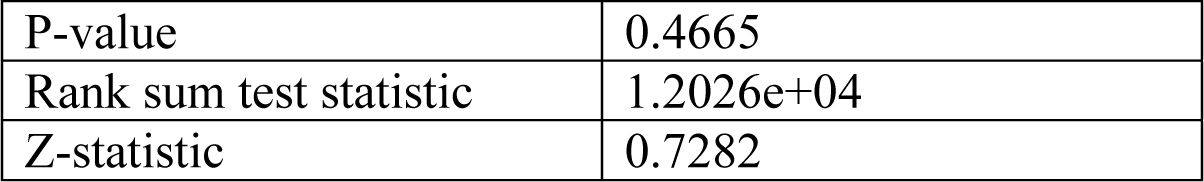

<u>Figure S5B</u>

Wilcoxon rank sum test of the τ_off_ of calcium spikes in zebrafish larva. (n = 96 cells from 3 GCaMP6f fish; n = 146 cells from 4 SomaGCaMP6f1 fish).

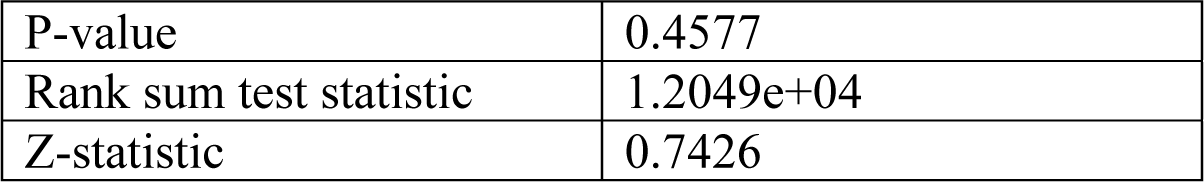

<u>Figure S5C</u>

Wilcoxon rank sum test of the GCaMP-spike rates for neurons expressing either GCaMP6f or SomaGCaMP6f1 (n = 24 cells from 2 GCaMP6f fish; n = 24 cells from 2 SomaGCaMP6f1 fish).

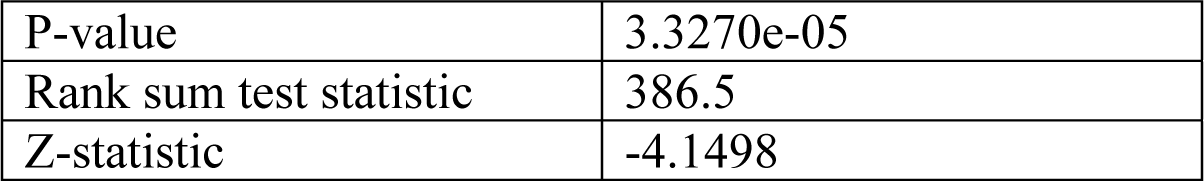

**Supplemental Table 12**: Statistical analysis for **Figure S6**

<u>Figure S6A:</u>

Baseline fluorescence in vivo in the dorsal striatum for GCaMP6f, SomaGCaMP6f1 and SomaGCaMP6f2. Kruskal-Wallis analysis of variance followed by post-hoc test via Steel’s test with GCaMP6f as control group (n = 75 neurons from 5 mice for GCaMP6f; n = 50 neurons from 2 mice for SomaGCaMP6f1; n = 80 neurons from 4 mice for SomaGCaMP6f2).

**Wilcoxon / Kruskal-Wallis Tests (Rank Sums)**

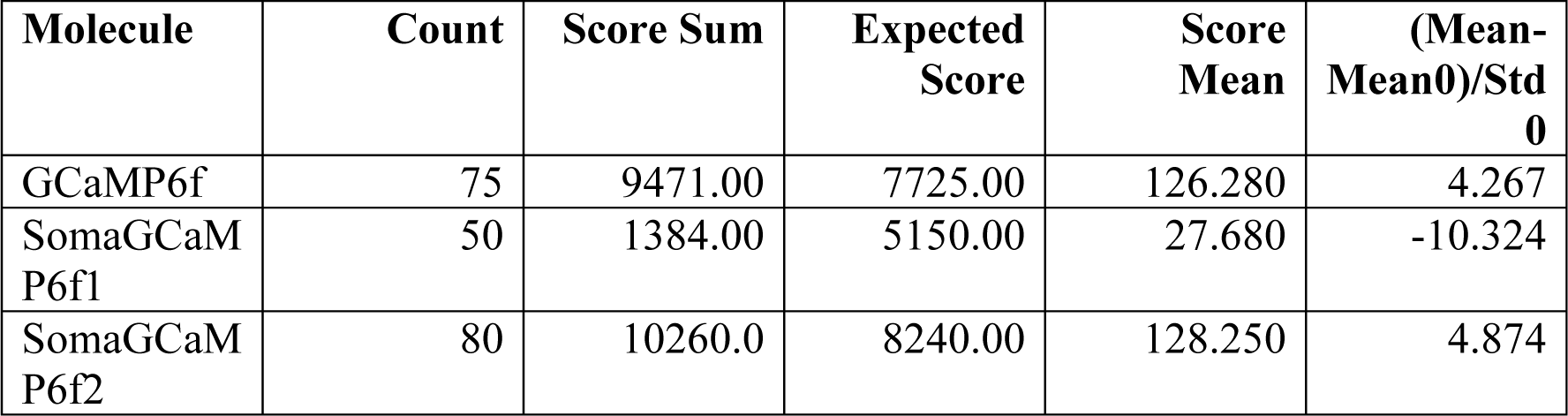

**1-Way Test, ChiSquare Approximation**

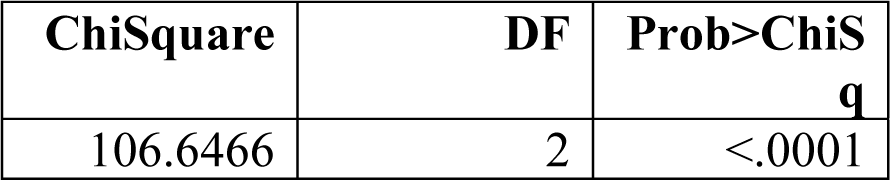

**Nonparametric Comparisons with Control Using Steel Method**

Control Group = GCaMP6f

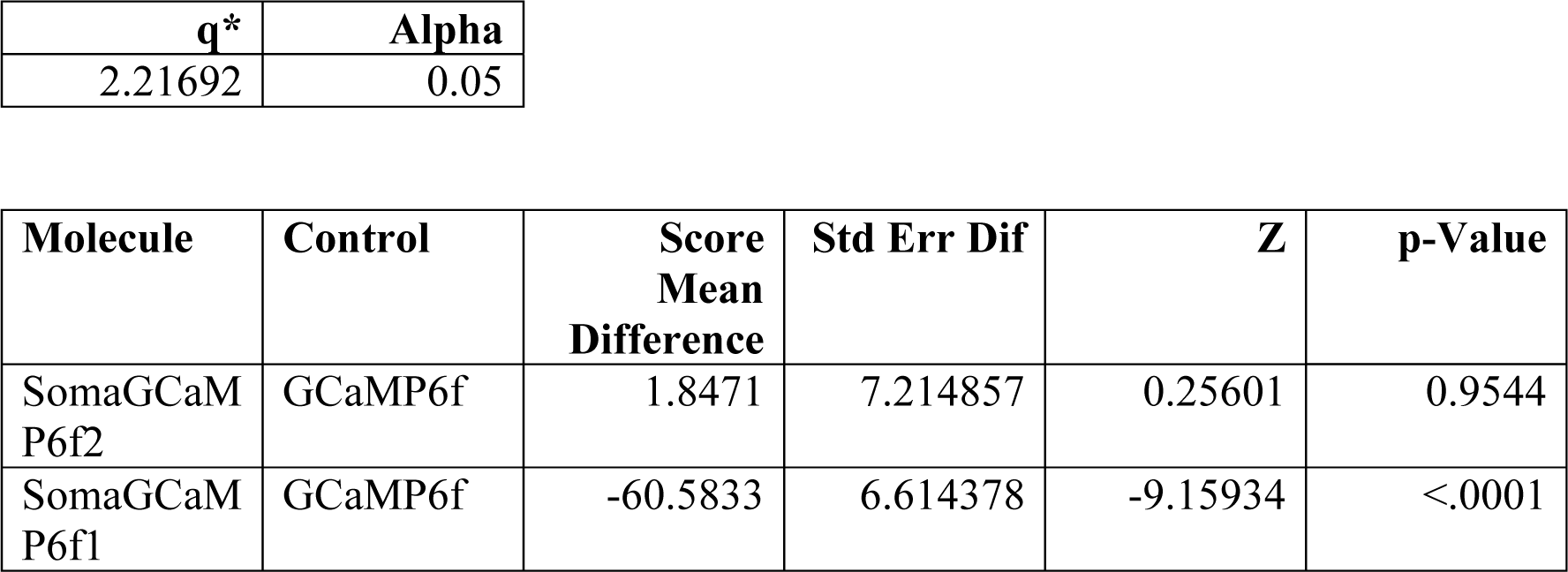

Figure S6B Wilcoxon rank sum tests comparing the rising times and the decay times between GCaMP6f and SomaGCaMP6f2 calcium signals for all detected calcium events (n = 930 neurons from 7 GCaMP6f mice; n = 594 neurons from 4 SomaGCaMP6f2 mice).

Rising times:

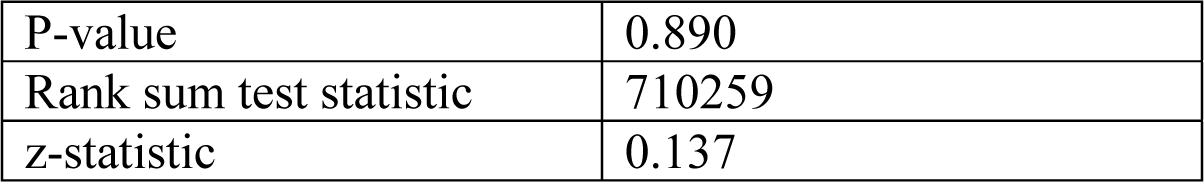

Decay times:

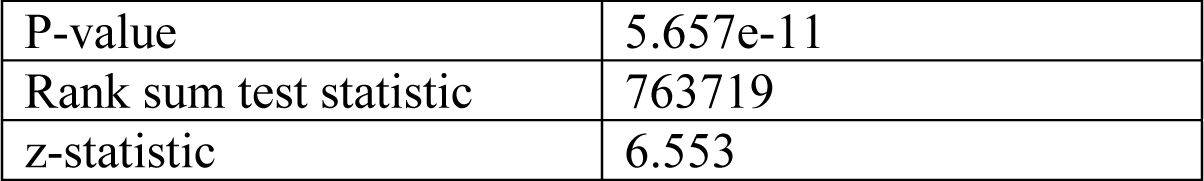

**Supplemental Table 13: Amino acid sequences for protein fragments used in this paper**

**AnkTail-motif (Ankyrin_G_ (1934-2333)):**

REGRIDDEEPFKIVEKVKEDLVKVSEILKKDVCVESKGPPKSPKSDKGHSPEDDWTEFSSEEIREARQAAASHAPSLPERVHGKANLTRVIDYLTNDIGSSSLTNLKYKFEEAKKDGEERQKRILKPAMALQEHKLKMPPASMRPSTSEKELCKMADSFFGADAILESPDDFSQHDQDKSPLSDSGFETRSEKTPSAPQSAESTGPKPLFHEVPIPPVITETRTEVVHVIRSYEPSSGEIPQSQPEDPVSPKPSPTFMELEPKPTTSSIKEKVKAFQMKASSEEEDHSRVLSKGMRVKEETHITTTTRMVYHSPPGGECASERIEETMSVHDIMKAFQSGRDPSKELAGLFEHKSAMSPDVAKSAAETSAQHAEKDSQMKPKLERIIEVHIEKGPQSPCE

**EE-RR:**

LEIEAAFLEQENTALETEVAELEQEVQRLENIVSQYETRYGPLGSLEIRAAFLRRRNTALRTRVAELRQRVQRLRNIVSQYETRYGPL

**AcidP1-BaseP1:**

AQLEKELQALEKENAQLEWELQALEKELAQGSGSAQLKKKLQALKKKNAQLKWKLQ

ALKKKLAQ

**nullsfGFP (mutation to abolish the fluorescence of the original sfGFP is underlined)**

MSKGEELFTGVVPILVELDGDVNGHKFSVRGEGEGDATNGKLTLKFICTTGKLPVPWPT

LVTTLT<u>G</u>GVQCFSRYPDHMKRHDFFKSAMPEGYVQERTISFKDDGTYKTRAEVKFEGD

TLVNRIELKGIDFKEDGNILGHKLEYNFNSHNVYITADKQKNGIKANFKIRHNVEDGSVQLADHYQQNTPIGDGPVLLPDNHYLSTQSVLSKDPNEKRDHMVLLEFVTAAGITHGMDE

LYK

**NLS**

RKRPSDLVHVFSPPRKK

**KGC**

KSRITSEGEYIPLDQIDINV

**ER2**

FCYENEV

**nullCoChR (mutation to abolish photocurrent of the original CoChR is underlined)**

MLGNGSAIVPIDQCFCLAWTDSLGSDTEQLVANILQWFAFGFSILILMFYAYQTWRATC

GWEEVYVCCVELTKVIIEFFHEFDDPSMLYLANGHRVQWLRYAEWLLTCPVILIHLSNL

TGLKDDYSKRTMRLLVSDVGTIVWGATSAMSTGYVKVIFFVLGCIYGANTFFHAAKVYI

ESYHVVPKGRPRTVVRIMAWLFFLSWGMFPVLFVVGPEGFDAISVYGSTIGHTIIDLMS<u>A</u>

NCWGLLGHYLRVLIHQHIIIYGDIRKKTKINVAGEEMEVETMVDQEDEETV

**KA2(1-150)**

MPAELLLLLIVAFANPSCQVLSSLRMAAILDDQTVCGRGERLALALAREQINGIIEVPAKARVEVDIFELQRDSQYETTDTMCQILPKGVVSVLGPSSSPASASTVSHICGEKEIPHIKVGPEETPRLQYLRFASVSLYPSNEDVSLAVS

**KA2(1-150)-Y76A**

MPAELLLLLIVAFANPSCQVLSSLRMAAILDDQTVCGRGERLALALAREQINGIIEVPAKARVEVDIFELQRDSQAETTDTMCQILPKGVVSVLGPSSSPASASTVSHICGEKEIPHIKVGPEETPRLQYLRFASVSLYPSNEDVSLAVS

**KA2(1-100)**

MPAELLLLLIVAFANPSCQVLSSLRMAAILDDQTVCGRGERLALALAREQINGIIEVPAKARVEVDIFELQRDSQYETTDTMCQILPKGVVSVLGPSSSP

**Ank(1-334) (Ankyrin_G_ (1-334))**

MAHAASQLKKNRDLEINAEEETEKKKKHRKRSRDRKKKSDANASYLRAARAGHLEKALDYIKNGVDVNICNQNGLNALHLASKEGHVEVVSELLQREANVDAATKKGNTALHIASLAGQAEVVKVLVTNGANVNAQSQNGFTPLYMAAQENHLEVVRFLLDNGASQSLATEDGFTPLAVALQQGHDQVVSLLLENDTKGKVRLPALHIAARKDDTKAAALLLQNDTNADIESKMVVNRATESGFTSLHIAAHYGNINVATLLLNRAAAVDFTARNDITPLHVASKRGNANMVKLLLDRGAKIDAKTRDGLTPLHCGARSGHEQVVEMLLDRAAP

**AnkCT-motif (Ankyrin_G_ (2334-2622))**

RTDIRMAIVADHLGLSWTELARELNFSVDEINQIRVENPNSLISQSFMLLKKWVTRDGKNATTDALTSVLTKINRIDIVTLLEGPIFDYGNISGTRSFADENNVFHDPVDGWQNETPSGSLESPAQARRLTGGLLDRLDDSSDQARDSITSYLTGEPGKIEANGNHTAEVIPEAKAKPYFPESQNDIGKQSIKENLKPKTHGCGRTEEPVSPLTAYQKSLEETSKLVIEDAPKPCVPVGMKKMTRTTADGKARLNLQEEEGSTRSEPKQGEGYKVKTKKEIRNVEKKTH

**AnkMB-motif (Ankyrin_G_ (1-800))**

MAHAASQLKKNRDLEINAEEETEKKRKHRKRSRDRKKKSDANASYLRAARAGHLEKALDYIKNGVDVNICNQNGLNALHLASKEGHVEVVSELLQREANVDAATKKGNTALHIASLAGQAEVVKVLVTNGANVNAQSQNGFTPLYMAAQENHLEVVRFLLDNGASQSLATEDGFTPLAVALQQGHDQVVSLLLENDTKGKVRLPALHIAARKDDTKAAALLLQNDTNADVESKSGFTPLHIAAHYGNINVATLLLNRAAAVDFTARNDITPLHVASKRGNANMVKLLLDRGAKIDAKTRDGLTPLHCGARSGHEQVVEMLLDRSAPILSKTKNGLSPLHMATQGDHLNCVQLLLQHNVPVDDVTNDYLTALHVAAHCGHYKVAKVLLDKKASPNAKALNGFTPLHIACKKNRIRVMELLLKHGASIQAVTESGLTPIHVAAFMGHVNIVSQLMHHGASPNTTNVRGETALHMAARSGQAEVVRYLVQDGAQVEAKAKDDQTPLHISARLGKADIVQQLLQQGASPNAATTSGYTPLHLAAREGHEDVAAFLLDHGASLSITTKKGFTPLHVAAKYGKLEVASLLLQKSASPDAAGKSGLTPLHVAAHYDNQKVALLLLDQGASPHAAAKNGYTPLHIAAKKNQMDIATSLLEYGADANAVTRQGIASVHLAAQEGHVDMVSLLLSRNANVNLSNKSGLTPLHLAAQEDRVNVAEVLVNQGAHVDAQTKMGYTPLHVGCHYGNIKIVNFLLQHSAKVNAKTKNGYTALHQAAQQGHTHIINVLLQNNASPNELTVNGNTAL

**AnkSB-motif (Ankyrin_G_ (801-1521))**

AIARRLGYISVVDTLKVVTEEIMTTTTITEKHKMNVPETMNEVLDMSDDEVRKASAPEK

LSDGEYISDGEEGEDAITGDTDKYLGPQDLKELGDDSLPAEGYVGFSLGARSASLRSFSS

DRSYTLNRSSYARDSMMIEELLVPSKEQHLTFTREFDSDSLRHYSWAADTLDNVNLVSS

PVHSGFLVSFMVDARGGSMRGSRHHGMRIIIPPRKCTAPTRITCRLVKRHKLANPPPMVE

GEGLASRLVEMGPAGAQFLGPVIVEIPHFGSMRGKERELIVLRSENGETWKEHQFDSKN

EDLAELLNGMDEELDSPEELGTKRICRIITKDFPQYFAVVSRIKQESNQIGPEGGILSSTTV

PLVQASFPEGALTKRIRVGLQAQPVPEETVKKILGNKATFSPIVTVEPRRRKFHKPITMTI

PVPPPSGEGVSNGYKGDATPNLRLLCSITGGTSPAQWEDITGTTPLTFIKDCVSFTTNVSA

RFWLADCHQVLETVGLASQLYRELICVPYMAKFVVFAKTNDPVESSLRCFCMTDDRVD

KTLEQQENFEEVARSKDIEVLEGKPIYVDCYGNLAPLTKGGQQLVFNFYSFKENRLPFSI

KIRDTSQEPCGRLSFLKEPKTTKGLPQTAVCNLNITLPAHKKETESDQDDAEKADRRQSF

ASLALRKRYSYLTEPSMKTVERSSGTARSLPTTYSHKPFFSTRPYQSWTTAPITVPGPAKS

GSLSSSPSNTPSA

**AnkSR-motif (Ankyrin_G_ (1534-1933))**

SPLKSIWSVSTPSPIKSTLGASTTSSVKSISDVASPIRSFRTVSSPIKTVVSPSPYNPQVASGT

LGRVPTITEATPIKGLAPNSTFSSRTSPVTTAGSLLERSSITMTPPASPKSNITMYSSSLPFK

SIITSATPLISSPLKSVVSPTKSAADVISTAKATMASSLSSPLKQMSGHAEVALVNGSVSPL

KYPSSSALINGCKATATLQDKISTATNAVSSVVSAASDTVEKALSTTTAMPFSPLRSYVS

AAPSAFQSLRTPSASALYTSLGSSIAATTSSVTSSIITVPVYSVVNVLPEPALKKLPDSNSF

TKSAAALLSPIKTLTTETRPQPHFNRTSSPVKSSLFLASSALKPSVPSSLSSSQEILKDVAE

MKEDLMRMTAILQTDVPEEKPFQTDLP

**K_V_2.1-motif (K_V_2.1(536-600))**

QSQPILNTKEMAPQSKPPEELEMSSMPSPVAPLPARTEGVIDMRSMSSIDSFISCATDFPE

ATRF

**rSK1-tail (rSK1(351-411))**

QAQKLRTVKIEQGKVNDQANTLADLAKAQSIAYEVVSELQAQQEELEARLAALESRLD

VLGASLQALPSLIAQAICPLPPPWPGPSHLTTAAQSPQSHWLPTTASDCG

**NaV1.6(II-III)**

TVRVPIAVGESDFENLNTEDVSSESDP

**Na_V_1.2(I-II)**

YEEQNQATLEEAEQKEAEFQQMLEQLKKQQEEAQAAAAAASAESRDFSGAGGIGVFSE

SSSVASKLSSKSEKELKNRRKKKKQKEQAGEEEKEDAVRKSASEDSIRKKGFQFSLEGS

RLTYEKRFSSPHQSLLSIRGSLFSPRRNSRASLFNFKGRVKDIGSENDFADDEHSTFEDND

SRRDSLFVPHRHGERRPSNVSQASRASRGIPTLPMNGKMHSAVDCNGVVSLVGGPSALT

SPVGQLLPEGTTTETEIRKRRSSSYHVSMDLLEDPSRQRAMSMASILTNTMEELEESRQK

CPPCWYKFANMCLIWDCCKPWLKVKHVVN

